# Characterization of a novel polyextremotolerant fungus, Exophiala viscosa, with insights into its melanin regulation and ecological niche

**DOI:** 10.1101/2021.12.03.471027

**Authors:** Erin C. Carr, Quin Barton, Sarah Grambo, Mitchell Sullivan, Cecile M. Renfro, Alan Kuo, Jasmyn Pangilinan, Anna Lipzen, Keykhosrow Keymanesh, Emily Savage, Kerrie Barry, Igor V. Grigoriev, Wayne R. Riekhof, Steven D. Harris

**Affiliations:** University of Nebraska-Lincoln, School of Biological Sciences, Lincoln, Nebraska 68588; University of Nebraska-Lincoln, Department of Biochemistry, Lincoln, Nebraska 68588; Iowa State University, Roy J. Carver Department of Biochemistry, Biophysics, and Molecular Biology, Ames, Iowa 50011; University of Nebraska-Lincoln, Department of Agronomy and Horticulture, Lincoln, Nebraska 68588; US Department of Energy Joint Genome Institute, Lawrence Berkeley National Laboratory, Berkeley, California 94720; Department of Plant and Microbial Biology, University of California Berkeley, Berkeley, California 94720; Iowa State University, Department of Plant Pathology and Microbiology, Ames, Iowa 50011; Iowa State University, Department of Entomology Ames, Iowa 50011

**Author notes:** Corresponding Author: Erin C. Carr, Legacy Mailing address: 1901 Vine St. Room E136 Lincoln, NE 68503.

## Abstract

Black yeasts are polyextremotolerant fungi that contain high amounts of melanin in their cell wall and maintain a primarily yeast form. These fungi grow in xeric, nutrient deplete environments which implies that they require highly flexible metabolisms and have been suggested to contain the ability to form lichen-like mutualisms with nearby algae and bacteria. However, the exact ecological niche and interactions between these fungi and their surrounding community is not well understood. We have isolated two novel black yeasts from the genus *Exophiala* that were recovered from dryland biological soil crusts. Despite notable differences in colony and cellular morphology, both fungi appear to be members of the same species, which has been named *Exophiala viscosa (i.e., E. viscosa* JF 03-3 Goopy *and E. viscosa* JF 03-4F Slimy). A combination of whole genome sequencing, phenotypic experiments, and melanin regulation experiments have been performed on these isolates to fully characterize these fungi and help decipher their fundamental niches within the biological soil crust consortium. Our results reveal that *E. viscosa* is capable of utilizing a wide variety of carbon and nitrogen sources potentially derived from symbiotic microbes, can withstand many forms of abiotic stresses, and excrete melanin that can potentially provide UV resistance to the biological soil crust community. Besides the identification of a novel species within the genus *Exophiala*, our study also provides new insight into the regulation of melanin production in polyextremotolerant fungi.

## Background

The production of melanin is a key functional trait observed in many fungi spanning the fungal kingdom (Bell & Wheeler, 1986; Zanne et al., 2020). The diverse protective functions of melanin (e.g., metal resistance, ROS and RNS tolerance, UV resistance) underscores its broad utility in mitigating the impacts of stress despite the potential cost of its synthesis (Schroeder et al., 2020). Fungi are capable of producing three different types of melanin: pheomelanin, allomelanin, and pyomelanin, all of which have their own independent biosynthetic pathways (Pal et al., 2014; Perez-Cuesta et al., 2020). Allomelanin is formed from the polymerization of 1,8-DHN, which requires the use of polyketide synthase for initial steps in production (Perez-Cuesta et al., 2020; Płonka & Grabacka, 2006) (Figure 1). Pyomelanin and pheomelanin share an initial substrate of Tyrosine, but pheomelanin derives from L-DOPA, whereas pyomelanin is created via tyrosine degradation (Perez-Cuesta et al., 2020; Płonka & Grabacka, 2006) (Figure 1). Allomelanin and pheomelanin are often referred to as DHN-melanin and DOPA-melanin respectively, given their chemical precursors.

**Figure 1:**
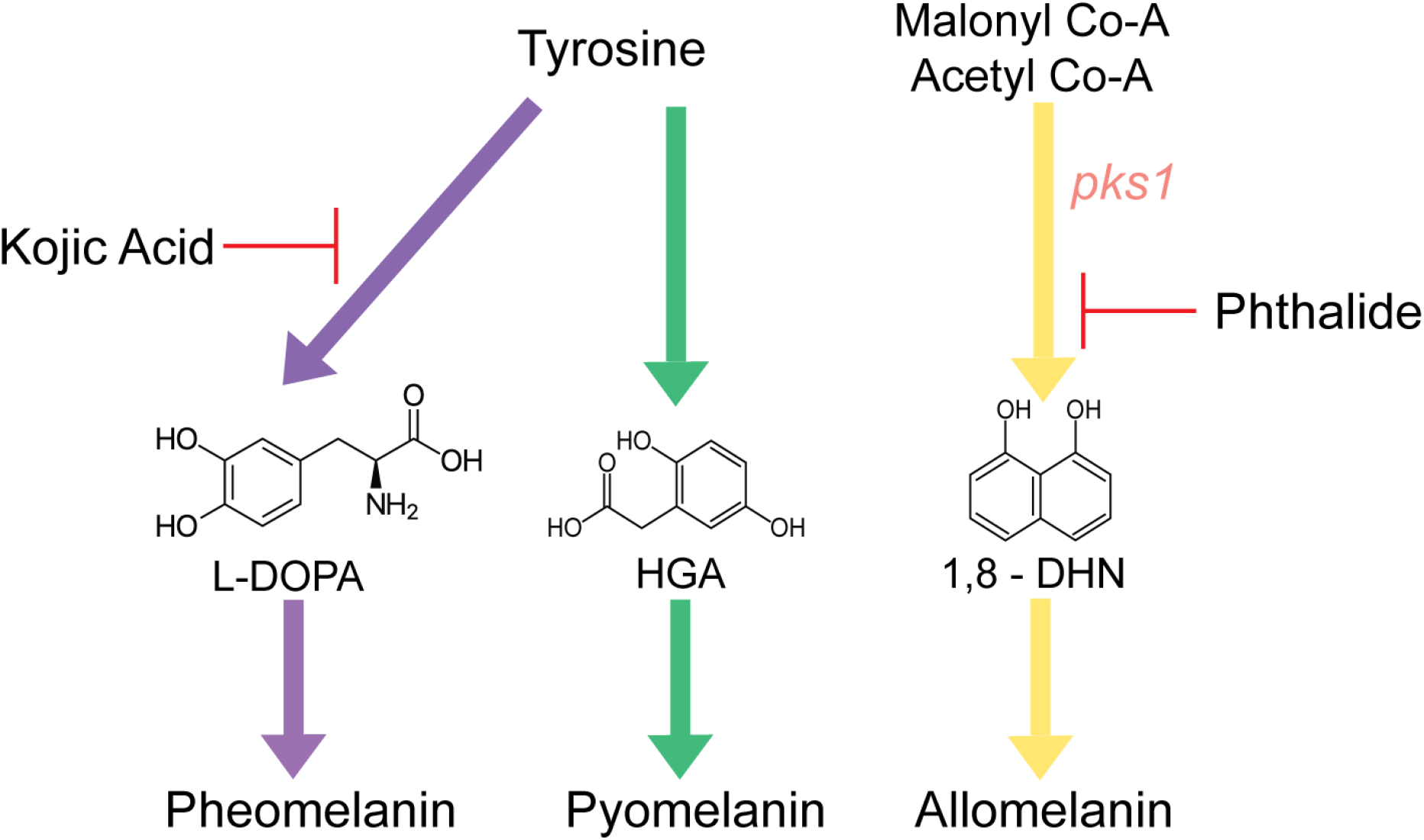
Visual summary of generalized fungal melanin production and the chemicals that block each pathway. Pheomelanin and Pyomelanin both use tyrosine as their starting reagent but have different biosynthesis pathways. Pheomelanin uses L-DOPA as a precursor to the final product and pyomelanin uses HMG as a precursor. Allomelanin’s starting components are malonyl co-A and acetyl co-A, and its immediate precursor is 1,8-DHN. The gene *pks1* is the first gene in the biosynthetic pathway for allomelanin. Kojic acid is a chemical blocker that blocks production of pheomelanin, and phthalide blocks production of allomelanin. Additionally, the chemical precursors of the final melanin product are provided, final melanin compound structures are still unknown and therefore the precursors are used in place of melanin structure itself. Detailed descriptions of these pathways can be found in: Perez-Cuesta et al. and Płonka & Grabacka (2020; 2006).

Unfortunately, due to the unique characteristics of melanins and their association with larger biomolecules, we do not know the complete structure of any type of melanin (Cao et al., 2021). However, given that we do know their chemical constituents, it is possible to draw some inferences about the relationship between structure and function of a particular type of melanin. For instance, out of the three types of melanin fungi can produce, only pheomelanin has a chemical precursor, 5- CysDOPA, with both nitrogen and sulfur in its structure (Płonka & Grabacka, 2006). Notably, all three fungal melanins are synthesized via independent pathways, which enables the targeted use of chemical inhibitors to block one pathway without interfering with the others. For example, previous studies have extensively used the chemical melanin blockers kojic acid and phthalide to block pheomelanin and allomelanin respectively (Pal et al., 2014) (Figure 1). Use of these chemical blockers allowed previous studies to identify the primary melanin biosynthetic pathway employed by individual fungal species (Pal et al., 2014).

Polyextremotolerant fungi are a polyphyletic group that can be divided morphologically into black yeast and microcolonial/meristematic fungi types (Gostinčar et al., 2012). These two sub-types are distinct in their morphology; black yeast fungi are primarily yeasts but can be dimorphic, whereas microcolonial fungi are typically filamentous, pseudohyphal, or produce other unique morphologies such as spherical cells (Ametrano et al., 2017; De Hoog et al., 2003; Gostinčar et al., 2011). However, all polyextremotolerant fungi share the capacity to produce melanin, which presumably accounts for much of their polyextremotolerance. Most polyextremotolerant fungi are in the subdivision Pezizomycotina, residing mainly within Eurotiomycetes and Dothidiomycetes (Gostinčar et al., 2009).

Melanin is arguably the defining feature of polyextremotolerant fungi given that they form unmistakably black colonies. Because of its structure and association with the cell wall, melanin imbues polyextremotolerant fungi with resistance to multiple forms of stress. Most commonly known is ultraviolet (UV) light resistance, as melanin absorbs light in the UV part of the spectrum (Kobayashi et al., 1993). However, melanin is also capable of quenching reactive oxygen species (ROS) and reactive nitrogen species (RNS), providing tolerance to toxic metals, reducing desiccation, and enabling use of ionizing radiation as an energy source (Cordero & Casadevall, 2017; Dadachova et al., 2007; Gessler et al., 2014; Płonka & Grabacka, 2006; Zanne et al., 2020). Collectively, these functions of melanin are thought to enhance the ability of polyextremotolerant fungi to colonize habitats that are otherwise inhospitable to most forms of life.

Polyextremotolerant fungi tend to occupy extreme niches such as rock surfaces, external brick and marble walls, soil surfaces, and even the inside of dishwashers (Gostinčar et al., 2009; Zupančič et al., 2016). Characteristic features of these environments includes the relative paucity of nutrients and the frequent presence of a community biofilm consisting of photosynthetic organisms and/or bacteria (Gostinčar et al., 2012). Strikingly, these microbes are rarely found alone in their habitats, which suggests that multi-species interactions between fungi, bacteria, and photosynthetic organisms underlie the formation and function of these communities. That is, the ability of polyextremotolerant fungi to successfully adapt to their niche must depend on complex yet poorly understood interactions with other microbes.

We have isolated two polyextremotolerant fungi from a biological soil crust (BSC) in British Columbia, Canada. These novel fungi are of the genus *Exophiala*, which has previously been found in BSCs (Bates et al., 2006). Biological soil crusts are unique dryland biofilms that form on the surface of xeric desert soils where little to no plants are able to grow (Belnap, 2003; Belnap et al., 2001). They are notable for their extensive cyanobacteria population, which seeds the initial formation of all BSCs and creates the main source of nitrogen for the rest of the community (Belnap, 2002). Once the initial crust is established, it is then inundated with a consortium of bacteria, fungi, algae, archaea, lichens, and mosses (Bates et al., 2010; Lan et al., 2012; Maier et al., 2016). This community is a permanent fixture on the land they occupy unless physically disturbed, much like other biofilms (Belnap & Eldridge, 2001; Donlan & Costerton, 2002). As a result of the arid conditions to which BSCs are often exposed, their resident microbes are constantly exposed to extreme abiotic factors that must be countered to enable survival (Bowker et al., 2010). Some of these abiotic extremes are: UV radiation (especially at higher altitudes and latitudes) (Bowker et al., 2002), desiccation and osmotic pressures (Rajeev et al., 2013), and temperature fluctuations both daily and annually (Belnap et al., 2001; Bowker et al., 2002; Pócs, 2009). Accordingly, microbes that reside within BSCs must therefore possess adaptations that allow them to withstand these abiotic extremes.

An extensive amount of research has been dedicated to certain members of the BSC community, but one such less studied microbe has been the “free-living” fungal taxa. These fungi are non-lichenized (Teixeira et al., 2017) yet are still thriving in an environment where there are no plants to infect or decompose (Belnap & Lange, 2003), and no obvious source of nutrients aside from contributions of the other members of the BSC community (Belnap & Lange, 2003). This would imply that even though these fungi are not lichenized *per se*, they would have to engage in lichen-like interactions with the BSC community to obtain vital nutrients for proliferation. While previous researchers have postulated the idea of transient interactions between non-lichenized fungi and other microbes (Gostinčar et al., 2012; Grube et al., 2015; Hom & Murray, 2014), confirmation of this in BSCs is potentially challenging given their taxonomic complexity.

Despite the importance of microbial interactions in enabling the successful formation of BSCs in a niche characterized by poor nutrient and water availability, the fungal components of BSCs and their relative functions within the interaction network remain poorly understood. Here, we combine genome sequencing with computational tools and culture-based phenotyping to describe two strains of a new species of black yeast fungi associated with BSCs. We report on their carbon and nitrogen utilization profiles, stress responses, and lipid accumulation patterns. In addition, we characterize their capacity for melanin production and generate valuable insight into mechanisms that might be involved in regulating the synthesis of these compounds so that melanin production contributes to the BSC community.

## Methods

### Fungal Strains and Media

Two strains of a novel species of fungi are described here: *Exophiala viscosa* JF 03-3F Goopy CBS 148801 and *Exophiala viscosa* JF 03-4F Slimy CBS 148802. *E. viscosa* is typically grown in malt extract medium (MEA; see Table 1 for media recipe) at room temperature in an Erlenmeyer flask at 1/10^th^ the volume of the flask, shaking at 160 rpm. Additional strains used in this study were *Saccharomyces cerevisiae* ML440 and BY4741, and *E. dermatitidis* wild type (WT) strain ATCC 34100. *S. cerevisiae* strains are grown in Yeast Peptone Dextrose medium (YPD; Table 1 for media recipe), and *E. dermatitidis* is grown in MEA.

**Table 1:** Medias used and their compositions

| **Media Name** | **Acronym** | **Composition (L<sup>-1</sup>)** |
| --- | --- | --- |
| **Malt Extract Agar Glucose** | MAG | 20 g Dextrose<br>20 g Malt Extract (Sigma 70167)<br>2 g Peptone (Fisher BP1420)<br>1 mL Hutner's Trace Elements<br>1 mL Vitamin Mix<br>15 g Agar |
| **Malt Extract Agar** | MEA | 20 g Dextrose<br>20 g Malt Extract<br>2 g Peptone<br>15 g Agar |
| **Minimal** | MN | 10 g Dextrose<br>50 mL 20x Nitrate salts<br>1 mL Hutner's Trace Elements |
| **Minimal + Vitamins** | MNV | 10 g Dextrose<br>50 mL 20x Nitrate salts<br>1 mL Hutner's Trace Elements<br>1 mL Vitamin Mix |
| **Minimal + N-acetyl Glucosamine** | MN+NAG | 10 g Dextrose<br>50 mL 20x Nitrate salts<br>1 mL Hutner's Trace Elements<br>4.74 g N-Acetyl Glucosamine (21.43 mM) |
| **Potato Dextrose Agar** | PDA | 24 g BD Difco Potato dextrose Broth powder<br>15 g agar (if not in powder) |
| **Spider** | Spider | 20 g Nutrient Broth<br>20 g Mannitol<br>4 g K<sub>2</sub>HPO<sub>4</sub><br>27 g Agar<br>pH adjusted to 7.2 with NaOH |
| **Yeast Extract Peptone Dextrose** | YPD | 20 g Dextrose<br>20 g Peptone<br>10 g Yeast Extract<br>20 g Agar |
| **V8** | V8 | 200 mL V8 Juice<br>2 g CaCO<sub>3</sub><br>15 g Agar |
| **Tyrosine Media** | Tyr | 20 g Dextrose<br>10 g Peptone<br>1 g Yeast Extract<br>20 g Agar |
| **Additives** | **Volume** | **Composition** |
| **20X Nitrate Salts/MN salts** | 1 L | 120 g NaNO<sub>3</sub> (remove for "MN salts")<br>10.4 g KCl<br>10.4 g MgSO<sub>4</sub>·7H<sub>2</sub>O<br>30.4 g KH<sub>2</sub>PO<sub>4</sub> |
| **Hutner's Trace Elements (TE)** | 100 mL | 2.2 g ZnSO<sub>4</sub>-7H<sub>2</sub>O<br>1.1 g H<sub>3</sub>BO<sub>3</sub><br>0.5 g MnCl<sub>2</sub>-4H<sub>2</sub>O<br>0.5 g FeSO<sub>4</sub>-7H<sub>2</sub>O<br>0.17 g CoCl<sub>2</sub>-6H<sub>2</sub>O<br>0.16 g CuSO<sub>4</sub>-5H<sub>2</sub>O<br>0.15 g Na<sub>2</sub>MoO<sub>4</sub>-2H<sub>2</sub>O<br>5 g EDTA (Disodium salt) |
| **Vitamin Mix** | 100 mL | 10 mg biotin<br>10 mg pyridoxin<br>10 mg thiamine<br>10 mg riboflavin<br>10 mg p-aminobenzoic acid (PABA)<br>10 mg nicotinic acid |

### Fungal Isolation and Identification Methods

Fungi were isolated from a BSC in British Columbia (B.C.), Canada, Latitude: 52.94997°, Longitude: -119.41919°. Soil samples were taken from the top 2 cm of the BSC. A 0.1 g portion of the crust was re-suspended in 1 mL of water, ground with a sterile micropestle, and diluted with a dilution factor (DF) of 10 till they reached 10,000x dilution. Each dilution was then spread out onto two different MEA petri plates containing either no antibiotics or a plate containing: Ampicillin (100 mg/L final concentration), Chloramphenicol (50 mg/L), Gentamycin (10 mg/L), and Cycloheximide (100 mg/L). The plates were then incubated in a Percival light incubator at 23 °C with a 12 hr light/dark cycle and examined daily using a dissection microscope to check for small black colonies. Once a potential black colony was seen, half of it was removed and transferred to a new MEA petri plate with no antibiotics. It was vital to examine the plates daily, because even in the presence of antibiotics many unwanted fast-growing microbes would grow on the plates and cover up the slower growing polyextremotolerant fungal colonies. Once a pure culture of each isolate was grown up (approximately 2 weeks), they were preserved in 30% glycerol and stored in the -80 °C freezer.

DNA sequencing of amplified internal transcribed spacer (ITS) sequences was used to initially identify the isolates. DNA was extracted using the Qiagen DNeasy Powersoil DNA extraction kit. Primers used to isolate the ITS region were: ITS1- (5’-TCC GTA GGT GAA CCT GCG G-3’) and ITS4- (5’-TCC TCC GCT TAT TGA TAT GC-3’) (White et al., 1990). A BioRad MJ Mini Personal Thermal Cycler was used, with the program set as: 1) 95 °C for 5:00 (5 minutes), 2) 94 °C for 0:30 (30 seconds), 3) 55 °C for 0:30, 4) 72 °C for 1:30, 5) Return to Step 2 35 times, 6) 72 °C for 5:00. Resulting PCRs were then checked via gel electrophoresis in 1% agar run at 80 V for 1 hr. Isolated ITS regions were sequenced using the Eurofins sequencing facility with their in-house ITS primers as the sequencing primers, and the sequences were subsequently identified using the basic local alignment search tool (BLAST) of the National Center for Biotechnological Information (NCBI) database to look for potential taxa matches.

### DNA Extraction and RNA Extraction for Whole Genome Sequencing and Annotation

A cetyltrimethylammonium bromide (CTAB) based DNA extraction method was performed to obtain high molecular weight DNA for whole genome sequencing. The DNA extraction method used was derived from (Cubero et al., 1999). Changes to the original protocol include: 1) switching polyvinylpolypyrrolidone (PVPP) for the same concentration of polyvinylpyrrolidone (PVP), 2) use of bead beating tubes with liquid nitrogen-frozen cells and the extraction buffer instead of a mortar and pestle for breaking open the cells, and 3) heating up the elution buffer to 65 °C before eluting the final DNA. These changes were made to optimize the protocol for liquid-grown fungal cells instead of lichen material. Cells for the DNA extraction were grown up in 25 mL of liquid MEA in 250 mL Erlenmeyer flasks for 5 days at room temperature with 160 rpm shaking. 1 mL of the grown cells was used for the DNA extraction after washing with water twice. Protocol for CTAB extraction method can be found on protocols.io.

RNA was obtained using the Qiagen RNeasy Mini Kit (Cat. No. 74104). Cells were grown in 25 mL of three different liquid media types (MEA, YPD, and MNV; see Table 1) in 250 mL Erlenmeyer flasks at room temperature for 5 days (as described above), and 1-2 mL of cells were used for the RNA extraction. Cells were washed with DEPC-treated water and flash frozen in liquid nitrogen in 1.5 mL microcentrifuge tubes. RNA extraction was then performed according to the methods by the RNeasy kit.

### Genome Assembly and Annotation

Both genomes and transcriptomes were sequenced using Illumina technology. For transcriptomes, a plate-based RNA sample prep was performed on the PerkinElmer Sciclone next generation sequencing (NGS) robotic liquid handling system using Illuminas TruSeq Stranded mRNA high throughput (HT) sample prep kit utilizing poly-A selection of mRNA following the protocol outlined by Illumina in their user guide: https://support.illumina.com/sequencing/sequencing_kits/truseq-stranded-mrna.html, and with the following conditions: total RNA starting material was 1 μg per sample and 8 cycles of PCR was used for library amplification. The prepared libraries were quantified using KAPA Biosystems’ next-generation sequencing library qPCR kit and run on a Roche LightCycler 480 real-time PCR instrument. The libraries were then multiplexed and prepared for sequencing on the Illumina NovaSeq sequencer using NovaSeq XP v1 reagent kits, S4 flow cell, following a 2x150 indexed run recipe.

BBDuk was used for trimming and filtering sequencing reads (https://sourceforge.net/projects/bbmap/). Raw reads were evaluated for artifact sequence by k-mer matching (k-mer=25), allowing 1 mismatch and detected artifacts were trimmed from the 3’ end of the reads. RNA spike-in reads, PhiX reads and reads containing any Ns were removed. Quality trimming was performed using the phred trimming method set at Q6. Finally, following trimming, reads under the length threshold were removed (minimum length 25 bases or 1/3 of the original read length – whichever is longer). Filtered reads were assembled into consensus sequences using Trinity ver. 2.3.2 (Grabherr et al., 2011). Specific Trinity options that were utilized were: --jaccard_clip, --run_as_paired, -- min_per_id_same_path 95, --normalize_reads.

For genomes, DNA library preparation for Illumina sequencing was performed on the PerkinElmer Sciclone NGS robotic liquid handling system using Kapa Biosystems library preparation kit. 200 ng of sample DNA was sheared to 300 bp using a Covaris LE220 focused-ultrasonicator. The sheared DNA fragments were size selected by double-SPRI and then the selected fragments were end-repaired, A-tailed, and ligated with Illumina compatible sequencing adaptors from IDT containing a unique molecular index barcode for each sample library. The prepared libraries were quantified using KAPA Biosystems’ next-generation sequencing library qPCR kit and run on a Roche LightCycler 480 real-time PCR instrument. The quantified libraries were then multiplexed with other libraries, and the pool of libraries was then prepared for sequencing on the Illumina HiSeq sequencing platform utilizing a TruSeq paired-end cluster kit, v4, and Illumina’s cBot instrument to generate a clustered flow cell for sequencing. Sequencing of the flow cell was performed on the Illumina HiSeq2500 sequencer using HiSeq TruSeq SBS sequencing kits, v4, following a 2x150 indexed run recipe.

An initial assembly of the target genome was generated using VelvetOptimiser version 2.1.7 (3) with Velvet version 1.2.07 (Zerbino & Birney, 2008) using the following parameters; “–s 61 –e 97 –i 4 –t 4, --o “-ins_length 250 -min_contig_lgth 500””. The resulting assembly was used to simulate 28X of a 2x100 bp 3000 +/- 300bp insert long read mate-pair library with wgsim version 0.3.1-r13 (https://github.com/lh3/wgsim) using “-e 0 -1 100 -2 100 -r 0 -R 0 -X 0 -d 3000 -s 30”. 25X of the simulated long read mate-pair was then co-assembled together with 125X of the original Illumina filtered fastq with AllPathsLG release version R49403 (Gnerre et al., 2011) to produce the final nuclear assembly. A simulated long read assembly was required because AllpathsLG requires both a long read and short read insert library and having both types of libraries was beyond the cost of this project. Therefore, a subsample of the original Illumina short read inserts and the newly simulated long read insert data was fed into AllpathsLG. This methodology was shown to produce superior assemblies over other assemblers with a single short insert library. The genome was then annotated using JGI Annotation pipeline (Grigoriev et al., 2014). The JGI Annotation Pipeline, 1) detects and masks repeats and transposable elements, 2) predicts genes using a variety of methods, 3) characterizes each conceptually translated protein using a variety of methods, 4) chooses a ‘best’ gene model at each locus to provide a filtered working set, 5) clusters the filtered sets into draft gene families, and 6) creates a JGI Genome Portal with tools for public access and community-driven curation of the annotation (Grigoriev et al., 2012; Kuo et al., 2014).

The assembly scaffolds were masked by RepeatMasker (http://www.repeatmasker.org/) using both the manually curated Repbase library (Jurka, 2000) and a library of de novo repeats discovered by RepeatScout (Price et al., 2005). Transcript-based gene models were derived by assembling Illumina-sequenced RNA reads into contigs which were then modeled on genomic scaffolds using EST MAP (http://www.softberry.com) and COMBEST (Zhou et al., 2015). Protein-based gene models were predicted using GeneWise (Birney & Durbin, 2000) and FGENESH+ (Salamov & Solovyev, 2000) seeded by BLASTx alignments of genomic sequence against sequences from the NCBI non-redundant protein set nr (Altschul et al., 1990). Ab initio gene models were predicted using GeneMark-ES (Ter-Hovhannisyan et al., 2008) and FGENESH, the latter trained on a set of putative full-length transcripts and reliable protein-based models. GeneWise models were completed using scaffold data to find start and stop codons. RNA contig alignments to the genome were used to verify, complete, and extend the gene models.

All predicted gene models were functionally annotated by the JGI Annotation Pipeline using InterProScan (Zdobnov & Apweiler, 2001), blastp alignments against nr and highly curated databases such as SwissProt (Bairoch et al., 2005), KEGG (Ogata et al., 1999), and Pfam (Bateman et al., 2004). KEGG hits were used to map EC numbers (Bairoch, 2000); InterPro, KEGG, and SwissProt hits were used to map GO terms (Ashburner et al., 2000). In addition, predicted proteins were annotated according to KOG classification (Koonin et al., 2004). Protein targeting predictions were made with signalP (Nielsen et al., 1999) and TMHMM (Krogh et al., 2001). Because the gene-prediction steps usually generated multiple gene models per locus, the Pipeline selected a single representative gene model for each locus based on protein similarity, RNA coverage, and functional annotation, leading to a filtered set of gene models for downstream analysis. All filtered model proteins in a genome were aligned \ to each other and to those of other related genomes by blastp. The alignment scores were used as a distance metric for clustering by MCL (http://www.micans.org/mcl/) into a first draft of candidate multigene families for each genome.

All genomic scaffolds, RNA contigs, gene models and clusters, and functional predictions thereof, may be accessed through MycoCosm Herpotrichiellaceae PhyloGroup Portal (https://mycocosm.jgi.doe.gov/Herpotrichiellaceae/) and the MycCosm Black Yeasts EcoGroup Portal (https://mycocosm.jgi.doe.gov/black_yeasts/). The assemblies and annotations of both genomes are available at the fungal genome portal MycoCosm (Grigoriev et al., 2014); https://mycocosm.jgi.doe.gov) and in the DDBJ/EMBL/GenBank repository under accessions JALDXI000000000 and JALDXH000000000.

### Phylogenetic Analyses

Our single-copy homologous protein sequence approximate maximum likelihood phylogenetic tree was constructed from the protein sequences of 24 taxa which represent a sampling of the family Herpotrichielleaceae in which *E. viscosa* JF 03-3F and JF 03-4F reside. We created this tree using the pipeline described in Kuo et al. (2014). To obtain the largest set of data for phylogenetic analysis available, we defined homologous proteins across all 24 genomes using Best Bidirectional Blast (BBB) pairs via blastp across all proteins of all the genomes from their FilteredModels or ExternalModels files available on Mycocosm, with an e-value 1e-05 cutoff for BBBs. Clusters were compiled using in-house JGI scripts. If there are e.g. N genomes, we first compute BBBs individually for each pair of genomes (in total N*(N-1)/2 pairs) and then using BBBs as an input computed for each pair of genomes. Another script finds BBBs which are present in all N genomes, so these are sets of N genes (one from each species or strain) which are BBB to each other in all N genomes. Resulting defined homologous proteins across all 24 genomes were then used as our sequences for the rest of the pipeline. These sequences were aligned with Mafft v7.123b with no alterations (Katoh et al., 2002). That alignment was then trimmed using Gblocks 0.91b with options -b4=5 -b5=h to have a minimum of 5 positions per block and allowing up to half of those to be gaps (Castresana, 2000). The resulting trimmed alignment was then input into FastTree version 2.1.5 SSE3 with options -gamma -wag (Price et al., 2009). The tree file was then input into FigTree for visual optimization, and edited in Adobe Illustrator to highlight the new species.

The combined Bayesian and maximum likelihood concatenated gene phylogenetic trees were constructed using 18S rDNA, 28S rDNA, and ITS sequences of *Exophiala, Pheoannellomyces,* and *Endocarpon* species available from NCBI GenBank. Accession numbers and strains of fungi used for phylogenetic analyses are listed in Table 2. Specific fungal species and genes were chosen based on relatedness to our new fungus via BLAST alignment, and the availability of all marker genes needed for analyses. The marker gene *RPB1* was also concatenated to the others tested for all species except *Phaeoannellomyces elegans* and *Exophiala crusticola* which did not have an available *RPB1* sequence in NCBI. Alignments of individual gene sets were performed using the program MEGA X (Kumar et al., 2018). Alignment of the sequences was done using MUSCLE aligner (Edgar, 2004). Gap open was set to - 400.00, gap extend set to 0.00, max memory was set to 2048 MB, max iterations was 16, cluster method used was UPGMA, and minimal diagonal length was 24. Trimming was done by hand, first by removing any excess ends, then only essential gaps that formed where one or two taxa contained inserted nucleotides were removed. This allowed for the parts of the ITS and *RPB1* regions that remain highly variable even across the same genus to be kept, whereas when trimming was attempted with Gblocks with all least stringent options selected, only the most conserved regions were maintained resulting in improper phylogenies (e.g., *Exophiala eucalyptorum* 121638 did not cluster with other *Exophiala* species but was eventually removed from the tree for clarity). The alignments were then uploaded to Geneious Prime 2023.0.4 (Kearse et al., 2012). Individual gene sets were then concatenated using the concatenate alignments tool.

**Table 2:**
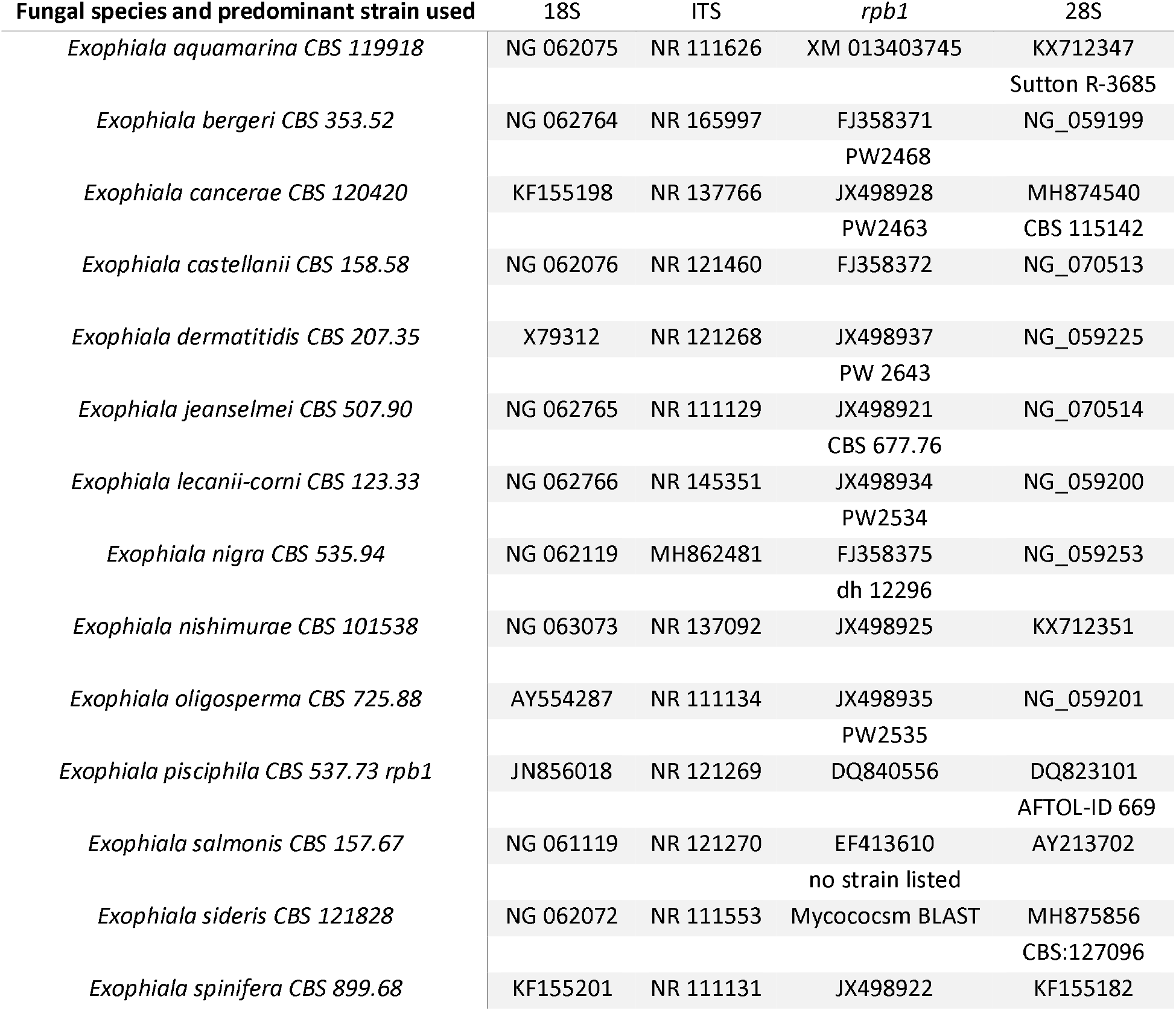
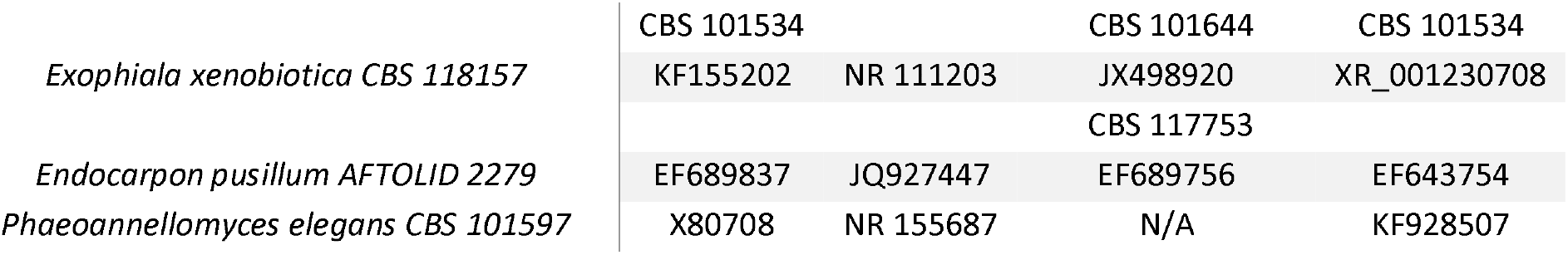
List of NCBI accession numbers of genes and strains of each fungal species used for concatenated phylogenetic analyses. If alternative strain was used for a gene the strain is listed under the accession number.

Phylogenetic estimates were performed using Bayesian inference via the Mr. Bayes plug-in ver. 2.2.4 in Geneious (Huelsenbeck & Ronquist, 2001; Ronquist & Huelsenbeck, 2003). For the Bayesian analysis we used Metropolis-coupled MCMC and the General Time Reversible (GTR) model with gamma distribution with invariant sites (Γ + I) the GTR + Γ + I model, with 5 gamma categories. We conducted the run with four heated chains at a temp, of 0.2 for 500,000 generations, sampling trees every 1,000 generations, with a burn length of 10,000, and a random seed of 20,512. The resulting phylogenetic tree was then used as the seed tree for Maximum Likelihood analyses. A maximum likelihood phylogenetic tree was created in MEGA X using the Maximum Likelihood statistical method and the bootstrap method for the test of phylogeny with 1,000 bootstrap replications. The substitution type used was nucleotide and not a protein coding sequence of DNA, and the model used for the substitution was the GTR + Γ + I with 5 discrete gamma categories. Gaps were included in the data treatment as many individual gaps are created in ITS and *RPB1* sequences after alignment in the variable regions. For tree inference options we used the nearest-neighbor-interchange (NNI) heuristic method, and had the initial tree be automated under the default settings-Neighbor joining (NJ)/bioNJ, the resulting Bayesian tree was used as the initial tree file, and no branch swap filter. The resulting tree in newick (.nwk) form with branch lengths and bootstrap values was then imported into the interactive tree of live (iTOL) program ver. 6.6 for visualization, and edited in Adobe Illustrator to add combined posterior probabilities and bootstrap values and highlight the new species (Letunic & Bork, 2021).

### Assemble Species by Automatic Partitioning for DNA-based Species Delimitation

We used Assemble Species by Automatic Partitioning (ASAP) to confirm our species delimitation that was observed in our phylogenetic analyses (Puillandre et al., 2021). We used the web-based version of ASAP (https://bioinfo.mnhn.fr/abi/public/asap/) using the Kimura (K80) model on our aligned ITS-only, 18S-28S-ITS gene concatenation alignment, and our 18S-28S-ITS-*RPB1* gene concatenation alignment. The resulting lowest ASAP score (smallest p-value and largest relative barcode gap) was chosen to be the most likely delimitation of species of that dataset.

### Mating Type Locus Identification

Mating loci for *E. viscosa* JF 03-3F and *E. viscosa* JF 03-4F were determined using the methods described by (Teixeira et al., 2017). Genes known to flank the MAT loci of most Chaetothyriales species include: *APN2*, *SLA2*, *APC5*, and *COX13*. The protein sequences of these genes from *Aspergillus nidulans* were used to BLASTp against the genomes of the new fungi. These gene sequences were obtained from Aspergillus Genome Database (now called FungiDB https://fungidb.org/) and protein-protein BLASTed using JGI’s Mycocosm. Once the genes were found, analysis of upstream and downstream flanking genes was performed until the mating gene MAT1-1 was located. Genes close to MAT1-1 and within the span of genes listed above were considered part of the MAT locus.

### Phenotyping Experiments

Phenotyping experiments of *E. viscosa* JF 03-3F and *E. viscosa* JF 03-4F were performed to characterize the new species’ capabilities and differences between the strains. Experiments performed were carbon utilization, nitrogen utilization, UV resistance, metal resistance, growth temperature range, budding patterns, lipid profiles, and growth capabilities on various fungal medias as described below.

#### Budding Pattern Determination

Protocols for observing the budding patterns of these new strains were derived from methods in Mitchison-Field et al. (2019) (Figure S1). A 1:1:1 ratio by weight of Vaseline, Paraffin, and Lanolin (VALAP) was combined in a glass bottle and heated to 115 °C to melt completely and kept at room temperature for later use. Heated solid MEA was aliquoted into a 50 mL tube for agar slab making. Isolates were grown in liquid MEA for 5 days prior to inoculation of slides. First, the VALAP was brought back up to 115 °C to melt completely for application. Then 5 μL of the 5-day old cells were diluted in 995 μL of liquid MEA. Agar slabs of MEA were made by microwaving the 50 mL tube of solid MEA until it melted, then pipetting 1 mL of the hot agar into a 1 cm x 2 cm mold formed out of cut strips of silicone and laid down in a sterile petri dish. This agar slab was allowed to solidify and was then cut in half to be 1 cm x 1 cm. Both the microscope cover slip and the microscope slide were wiped down with 70% ethanol to ensure clean and sterile growth conditions for the cells. 6 μL of the diluted cells was pipetted onto the center of the sterile slide, then one square of the agar slab was carefully placed on top of the cells in media, 8 μL of MEA was pipetted onto the top of the agar slab, and the coverslip was placed over the agar slab. Using a small paintbrush, the melted VALAP was carefully painted onto the gap between the coverslip and the microscope slide to seal off the coverslip. Finally, a 23-gauge needle was used to poke holes in the solidified VALAP to allow for gas exchange. The slide was then placed with the microscope slide facing down onto the inverted microscope EVOS fl set to the 40x objective. Once an adequate number of cells was observed in frame, the cells were allowed to settle for 2 hours before imaging began. Images were then automatically taken every 30 mins for 72 hours. Videos of the budding pattern were created using Adobe Premiere Pro.

#### Growth of E. viscosa JF 03-3F and E. viscosa JF 03-4F on Different Medias

Nine different fungal media were used to observe the growth of these novel fungi. These media have been used for identification purposes and will be useful for future identification of this species from other locations. Media used in this experiment were: MAG, MEA, MN, MNV, MN+NAG, PDA, Spider, YPD, YPD+TE, and V8 (Table 1). Both isolates were first grown in 25 mL of liquid MEA in a 250 mL Erlenmeyer flask at room temperature, shaking at 200 rpm for 5 days. Then 1 mL of each fungus was aliquoted and washed 3 times with water. Washed cells were then applied to the media in three ways: 5 μL spotting (pipetting 5 μL of cells onto the plate), toothpick poking (poking a sterile toothpick tip into the suspended cells and then onto a plate), and metal loop streaking (placing sterile metal loop into suspended cells, then spreading cells onto plate in a decreasing manner). Cells were then allowed to grow on the plates for 14 days. This provided us with different plating techniques that could potentially yield different morphologies. Descriptions of colony color morphologies follow color nomenclature from Ridgway (1912).

#### Carbon Utilization

Carbon utilization of each isolate was determined using a BioMerieux ID C32 carbon utilization strip (Cat. No. 32200-1004439110). These strips have 30 different carbon sources in individual wells, one well with no carbon source for a negative control, and one well with esculin ferric citrate. The esculin (β-glucose-6,7-dihydroxycoumarin) ferric citrate assay is a colorimetric assay originally used to identify Enterobacteria via production of a dark color in positive cultures for esculin degradation (Edberg et al., 1977), and was co-opted to identify *Cryptococcus neoformans* (Edberg et al., 1980). Therefore, the dark coloration that forms is due to the nature of the colorimetric assay, not due to extreme amounts of melanized fungal growth. The inoculated strips were kept in a plastic box container with a lid and lined with moist paper towels to reduce drying. The initial inoculum of cells was prepared in 25 mL of MEA shaking at room temp for 5 days. Inoculation of the strips was done according to the instructions provided by the vendor, and each strain was inoculated in three separate strips for triplicate replication. Cells were diluted to the kit requirement of McFarland standard #3 (McFarland, 1907) before starting the inoculum. Growth in the ID strip lasted 10 days before evaluation. Growth in each well was observed and evaluated by eye. Each well was compared to the negative control (no carbon) and the positive control (dextrose). Nuances of the fungal growth on individual carbon sources required a more gradual scale, and so scores were adjusted to form a range of 1-5 to allow for more accurate average calculation between the three replicates. For this scale: ”1” was no growth or equal to the kits “-“, ”5” was the most growth and equivalent to the kit’s “+”; numbers in between allowed us to average out the replicates to ensure we had a sliding scale of utilization rather than multiple variable “V” utilizations without a clear idea of how variable the utilization was. Since triplicates were performed, we averaged the growth results in each carbon source, and then determined the fungus’ carbon utilization profile from the averages. The scale used goes as follows: no growth “-“ = 1.0-1.9, variable growth “v” = 2.0-3.6, growth “+” = 3.7-5.0.

#### Wavelength for Optical Density Determination

The proper wavelength for optical density (OD) readings of these new strains was determined by creating a standard curve. We performed a full spectrum analysis of serially diluted cells of these fungi at their exponential growth phase. Cells were allowed to grow in 25 mL of MEA in a 250 mL Erlenmeyer flask for 5 days. Then 200 μL of culture was removed, washed 3x with water, and diluted 1:1 in water (100 μL of cells to 100 μL of MEA) 9 times to reach a maximum dilution of 1:256 (DF = 512). OD was measured at wavelengths from 300 nm – 700 nm in 10 nm increments. Then the R^2^ values for: OD to percent difference at 25% dilution (DF=4) and below, OD to percent difference at 50% dilution (DF=2) and below, OD to cell count at 2.19x10^6^ cells/mL (DF=8) and below, OD to cell count at 4.38x10^6^ cells/mL (DF=4) and below, and OD to the consistency of the actual cell counts percent difference at 12.5% dilution (DF=8) and below, were compared across all wavelengths to determine the most optimal wavelength. The most optimal wavelength would have the highest R^2^ values for all comparisons except OD to the consistency of the actual cell counts percent difference at 12.5% dilution and below which would have the lowest R^2^ value. Optical density reads were done on the BioTek Synergy H1 hybrid spectrophotometer.

#### Nitrogen Utilization

Nitrogen utilization tests were performed using ten different nitrogen conditions in MN base media. 100 mM was the concentration used for all compounds that contained one nitrogen atom per molecule: proline, ammonium tartrate dibasic, serine, sodium nitrate, glycine, glutamate, and aspartate; 50 mM was the concentration used for urea because it has two atoms of nitrogen per molecule; 1% w/v of Peptone was used as a positive control and no nitrogen was added as a condition for a negative control (Table 3). This concentration 100 mM was used because our standard growth media YPD contains 20 g/L of peptone, which contains 14% total nitrogen by weight. 14% total nitrogen per gram of peptone is 1.4 g of nitrogen per 100 mL of water, which if we use 1% peptone as a positive control that equals 100 mM total nitrogen in 1% of peptone. Liquid minimal media (MN) with MN salts (not 20x Nitrate salts) was used with the varying nitrogen sources to ensure that no alternative nitrogen source would be available to the fungi. Fungi were first grown up in liquid MEA for 5 days at room temperature to reach maximum density. Then, 1 mL of cells was removed and washed three times with water. 99 μL of each nitrogen source-containing medium was added to six wells in a 96-well plate, for six replicates, and 1 μL of the washed cells was added to each well. 100 μL of each medium was also added to one well each without cells to blank each condition, because the different nitrogen sources created different colors of medium. Daily growth was measured from day 0 to day 7 at 420 nm (the resulting optimal wavelength for OD reads), using the BioTek Synergy H1 hybrid spectrophotometer.

**Table 3:** Nitrogen sources, concentrations, and providers

| Nitrogen Source | Concentration | Catalog number |
| --- | --- | --- |
| No Nitrogen | N/A | N/A |
| Peptone | 1% w/v | Fisher Brand: BP1420 |
| L-Proline | 100 mM | Sigma: P-0380 |
| Ammonium tartrate | 100 mM | Sigma: A2956 |
| L-Serine | 100 mM | Sigma: S4500 |
| Sodium Nitrate | 100 mM | Fisher Brand: S343 |
| Glycine | 100 mM | Fisher BioReagents: BP381 |
| L-Glutamic acid | 100 mM | Sigma: G1251 |
| L-Aspartic acid | 100 mM | Sigma: A9256 |
| Urea | 50 mM | Alfa Aesar: A12360 |

#### Optimal Growth Temperature and Range of Growth Temperatures

To determine the temperature resistance range and optimal growth temperature for each isolate, we grew both fungi at 4 °C, 15 °C, 23 °C (i.e., ambient room temperature), 28 °C, 37 °C, and 42 °C. Isolates were first grown up in 25 mL of MEA for 5 days at room temperature to maximum density. Then 1 mL of cells was removed, and a 10x serial dilution was made from 0x to 100,000x, using pre-filled 1.5 mL tubes with 900 µL of MEA and adding 100 µL of the previous tubes each time. Then 5 µL of each serial dilution was spotted onto a square MEA plate which allowed us to determine the carrying capacity of each isolate at the different temperatures. Plates were kept at their respective temperatures for 7 days before observations were made, however the 37 °C and 42 °C plates were only kept at those temperatures for two days and the incubators required cups of water inside of them to prevent the plates from dehydrating. Plates grown in 42 °C and 37 °C were then allowed to grow at room temp for up to a week to determine if the isolates died at these temperatures or if their growth was just arrested.

#### UV Resistance

Resistance to UV light was observed to determine if these black fungi, with highly melanized cell walls and constant exposure to sunlight in their natural habitat, were UV resistant. To determine this, we used the UVP HL-2000 HybriLinker UV crosslinker as our source of UV light, which has a UV wavelength of 254 nm. Lower wavelengths (100-280 nm) are of the UV-C range, they are considered ionizing radiation and are the most detrimental to living organisms, but are completely blocked by the ozone layer (Molina & Molina, 1986; Schreier et al., 2015). Therefore, using this wavelength we are able to push our organisms beyond the UV limits found in their natural habitat and test extreme amounts of UV exposure.

The fungi were inoculated in 25 mL of MEA in a 250 mL Erlenmeyer flask and let grow under shaking conditions at 200 rpm for 5 days at room temperature to reach maximum density. 100 µL of this culture was then spread out onto 6 MEA plates, using a glass spreader. Three plates were kept as the control growth, to compare to the three other plates which were exposed to the UV light. Experimental plates were placed inside of the UV crosslinker with their lids taken off. Then the plates were exposed to 120 seconds of UV light from a distance of 9.5 cm to the light source at 10,000 μJ/cm^2^ (254 nm) (Frases et al., 2007). We then wrapped all plates in aluminum foil and placed them in the Percival light incubator set at 23 °C for 2 days. Placing UV-exposed cells in complete dark after exposure is essential for limiting repair via photoreactivation (Weber, 2005). After 2 days the plates were removed from the aluminum foil and left in the incubator for 5 more days before final observations were made. To determine whether a particular isolate was resistant to UV exposure, the growth of the fungi exposed to UV was compared to the control growth in imageJ, by comparing the pixel areas of the plates.

#### Metal Resistance

Metal resistance is a relatively universal trait in many polyextremotolerant fungi. Due to the under-studied nature of this particular characteristic in BSCs and fungi, we decided to test if any of our isolates were resistant to any heavy metals which would indicate possible bioremediation capacity. In order to test metal resistance, we used the antibiotic disk method by aliquoting metal solutions onto paper discs and observing zones of clearance. Metals and concentrations used are listed in Table 4. For testing, 5 µL of each metal solution was aliquoted onto a dry autoclaved Whatman filter paper disc which was created using a standard paper hole puncher. These discs were then allowed to air dry and kept at 4 °C for up to a week. Initial growth of the fungi (*E. viscosa* JF 03-3F, *E. viscosa* JF 03-4F, *E. dermatitidis*, and *S. cerevisiae* ML440) was done in 25 mL of MEA, shaking at 200 rpm for 5 days at room temperature. We then spread 100 μL of each fungus onto 100 mm sized MEA plates using a glass spreader to create a lawn. Using flame sterilized tongs our metal paper discs were placed onto the center of the petri dish on top of the fungal lawn and lightly pressed down to ensure the metal disc was touching the plate completely. These plates were then placed in the Percival light incubator at 23 °C with a 12 hr light/dark cycle for up to 2 weeks. Once a zone of clearing was clearly visible amongst the fungal growth (1-2 weeks), the zone of clearing was then measured in cm. Generally, large zones of clearing indicated sensitivity to the metal, whereas zones of reduced size were consisted with resistance to the metal.

**Table 4:** Metals used and their concentration

| Metal | Concentration | Catalog Number |
| --- | --- | --- |
| Fe(III)SO<sub>4</sub> | 0.6 M | Fisher: I146 |
| CoCl<sub>2</sub> | 0.5 M | Sigma: C-2644 |
| NiCl<sub>2</sub> | 1.5 M | Sigma-aldrich: 223387 |
| CuCl<sub>2</sub> | 1.5 M | Sigma: 203149 |
| CdCl<sub>2</sub> | 10 mM | Fisher: 7790-84-3 |
| AgNO<sub>3</sub> | 0.47 M | Alfa Aesar: 7761-88-8 |

#### Lipid Profiles

Comparison of the lipid production of *S. cerevisiae*, *E. dermatitidis*, *E. viscosa* JF 03-3F, and *E. viscosa* JF 03-4F was performed in the presence of fermentable vs. non-fermentable sugars in high and low nitrogen. To test these conditions, we grew all four fungi in four different media types. 1) MEA; 2) MEA + 20 g/L of peptone instead of 2 g/L; 3) MEA with the dextrose replaced with the same weight amount of glycerol; 4) MEA with glycerol instead of dextrose and 20 g/L of peptone instead of 2 g/L. All four fungi were first inoculated in 25 mL of liquid MEA in a 250 mL Erlenmeyer flask and shaken at 200 rpm for 5 days at room temperature to reach peak density. Then 100 μL was inoculated into 5 mL of each media in a size 25 mm tube, placed in a roller drum and allowed to grow at room temperature for 5 days.

To observe their lipid profile, we performed a standard Bligh Dyer lipid extraction (Bligh & Dyer, 1959). Equal wet weight of each organisms’ cells was pelleted and re-suspended in 2 mL of methanol inside of 16 mm glass tubes. Tube openings were covered in Duraseal before applying the lid of the tube, then samples were boiled for 5 minutes and let cool for 10 minutes. Then 2 mL of chloroform and 1.6 mL of 0.9% NaCl were added, and the tubes were vortexed to fully mix. Tubes were then centrifuged at 5000 rpm for 5 minutes to separate the layers. The bottom lipid layer of the suspension was removed and placed in a new glass tube which was then dehydrated using nitrogen gas till the samples became fully dried. Dehydrated samples were then re-suspended with 100 μL of a 9:1 ratio of chloroform : methanol to run the thin layer chromatography (TLC) with.

For all samples except the *S. cerevisiae*, 7 μL of the lipid suspension was used to dot the TLC. For *S. cerevisiae,* 10 μL of the lipid suspension was needed. The solvent systems used for TLC were chloroform : methanol : glacial acetic acid: water 85:12.5:12.5:3 for the polar lipid solvent system, and petroleum ether : diethyl ether : acetic acid 80:20:1 for the neutral lipid solvent system. The TLC plates were loaded with 7 or 10 μL of the re-suspended samples, and they were placed in the polar solvent system for approximately 30 minutes (half-way up the plate) before transferring to the neutral lipid solvent system in a separate container till the solvent front reached just a few cm below the top of the plate. The plate was then removed and dried for 15 minutes, until the solution on the plate was no longer volatile by smell, and the plate was placed in the presence of iodine (Sigma-Aldrich cat. No. 207772) in a glass chamber for 5 minutes until all the lipids were visible. The plates were then immediately placed in plastic covers and scanned and photographed for visualization and documentation.

### Melanin Biosynthesis and Regulation Experiments

#### Melanin Biosynthesis Gene Annotation

Melanin biosynthesis in fungi occurs via three different pathways: the 1,8-DHN pathway which creates allomelanin, the L-DOPA pathway which creates pheomelanin, and the tyrosine degradation (HGA) pathway which creates pyomelanin (Cao et al., 2021; Gessler et al., 2014) (Figure 1). Most fungal species only contain one melanin biosynthetic pathway, but there are many species in Pezizomycotina, particularly in the genera *Aspergillus* and *Exophiala*, which are capable of producing two or all three forms of melanin (Perez-Cuesta et al., 2020; Teixeira et al., 2017). For that reason, we decided to manually annotate the *E. viscosa* JF 03-3F and *E. viscosa* JF 03-4F genome sequences to determine if they too possessed multiple melanin biosynthetic pathways. In all cases, the relevant *A. niger* and *A. fumigatus* proteins were used as queries (Chen et al., 2014; Teixeira et al., 2017). Protein sequences for each gene were found using the Aspergillus genome database (AspGD) (now called FungiDB https://fungidb.org/) and were tested using protein-protein BLAST against the filtered model proteins database of *E. viscosa* JF 03-3F *and E. viscosa* JF 03-4F on Mycocosm. Since *A. niger* contains paralogs for some melanin biosynthetic genes, all genes listed in Teixeira et al. and Chen et al. (2014; 2017) were used as queries for BLAST searches. Additionally, the enzyme sequences for scytalone dehydratase (Arp1) and 1,3,8-trihydroxynapthelene reductase (Arp2) are not homologous between *A. niger* and *A. fumigatus* (Jørgensen et al., 2011), therefore both species’ protein sequences were used for BLASTing against *E. viscosa* JF 03-3F and *E. viscosa* JF 03-4F. Once the melanin biosynthetic genes in *E. viscosa* JF 03-3F and *E. viscosa* JF 03-4F were identified, their highest matching protein sequences were then reverse BLASTed to the *A. niger* genome to determine the reciprocal best hit to ensure true homology. Similarity scores (>40%), e-values (<10^-6^), and bit scores (>50) of the protein-protein BLASTs followed standards for homology laid out by Pearson (2013).

#### Regulation of Melanin Production Using Chemical Blockers

Once it was established that both isolates contain the potential for production of all three fungal melanins, we wanted to determine which melanin was actively being produced using chemical blockers of the 1,8-DHN and L-DOPA melanin production pathways. DHN-melanin blocker phthalide and the L-DOPA melanin blocker kojic acid, were both used to determine which melanin was being actively produced. Stock solutions were made according to (Pal et al., 2014): phthalide was diluted in 70% ethanol, and kojic acid in DMSO. Three separate experiments were performed using these melanin blockers, to determine which method would provide the most informative results.

The first was the disc diffusion method whereby Whatman filter paper discs were autoclaved and impregnated with 5 μL of either 10 mM of phthalide or 10 mg/mL of kojic acid. Impregnated filter paper discs were then placed on top of freshly spread lawns of either isolates on both MEA and YPD. Lawns were of 50:50 diluted 5-day old cells grown in MEA, and 100 µL of this dilution was spread onto the petri plates with a glass spreader. These plates were then grown at 23 °C with 12 hr light/dark cycles for 5 days. Additionally, both a kojic acid disc and phthalide discs were placed on fungal lawns ∼4 cm apart simultaneously to observe their specific melanin-blocking capabilities on the same cells.

Next, we tried adding the melanin blockers directly to the medium as was done in Pal et al. (2014). Since melanin is more universally distributed in *Exophiala* cells compared to *Aspergillus* cells, we decided to use the highest concentration of both kojic acid and phthalide that was used by Pal et al. (2014), which was 100 mM of each compound. This concentration of each compound was added to both solid YPD and MEA after autoclaving, chemicals were added both individually and combined. These plates were then used for two forms of growth experiments, serial dilutions and lawn growth. We spread a lawn of either fungus onto YPD and MEA with and without kojic acid, phthalide, and both compounds at 100 mM each. And we performed a 10x serial dilution of both *E. viscosa* JF 03-3F and *E. viscosa* JF 03-4F up to 10,000x diluted, and spotted 5 μL of each dilution onto MEA plates with and without Kojic acid, Phthalide, and both compounds. We let both growth experiments grow at 23 °C for 5 days with a 12 hr light/dark cycle.

#### Evaluating the Cause of Melanin Excretion

To assess the role of organic nitrogen levels in melanin excretion, we initially switched the concentration of peptone added to YPD and MEA media such that the new media were YPD + 0.2% peptone, and MEA + 2% peptone. *E. viscosa* JF 03-3F and *E. viscosa* JF 03-4F were grown in liquid MEA for 5 days with shaking at room temperature and were plated onto these new media using the same technique as described above for growth comparison on different media. To determine if a more gradual increase in peptone would correlate with a gradual excretion of melanin, we took the base media of MEA (solid) and changed the concentration of peptone to 0.1%, 0.5%, 1%, 1.5%, 2%, 2.5%, 3%, 3.5%, 4%, and 5%. We then spotted 5 μL of both strains onto the plates after performing a 10x serial dilution up to 10,000x dilution. The plates were grown at 23° for 10 days with a 12 hr light/dark cycle.

Additionally, we tested if tyrosine levels (a direct precursor for L-DOPA and HGA-derived melanin) were the source of the increased melanin excretion in the increased peptone experiment. Growth of 5 μL spots of *E. viscosa* JF 03-3F and *E. viscosa* JF 03-4F were grown on "Tyrosine media” (Jalmi et al., 2012) (2% w/v dextrose, 1% w/v peptone, 0.1% w/v yeast extract (Table 1)): without added tyrosine, with 0.5 g/L tyrosine, with 1 g/L tyrosine, and MN-nitrate 2% w/v dextrose with 0.5 g/L tyrosine as the sole nitrogen source. Colonies were allowed to grow for 19 days and then observations were made to see if melanin excretion was induced by different tyrosine levels.

#### Melanin Extraction and Spectrophotometric Measurements

Extraction of melanin from a variety of sources has been performed using two different approaches: chemical extraction and enzymatic extraction (Pralea et al., 2019). We performed both chemical and enzymatic extractions on our fungi’s melanin. The enzymatic extraction method that was used came from Rosas et al. (2000), and the chemical extraction method, which has been used more extensively in previous works, was derived from Pal et al. (2014). Pal et al.’s method for extraction and purification of melanin from culture filtrate was adapted and used for all future melanin extractions, mainly supernatant melanin extractions. Adjustments to the Pal et al. method included: the 6M HCl precipitation took multiple days instead of overnight for melanin to precipitate, and the protocol was stopped when 2M NaOH was added to the extracted melanin. We did not continue on to re- precipitation and drying of the melanin as this product did not reprecipitate in any solvents used.

Exact methods are as follows. 10 mL of culture was centrifuged at 3,000x g for 5 minutes, and the resulting supernatant was filter sterilized through a 2 μm filter to ensure all cells were removed. The filtered supernatant was then transferred into a 50 mL centrifuge tube, and 40 mL of 6M HCl was added to the tube. The filtrate was then allowed to precipitate out for up to two weeks. Precipitated solutions were then centrifuged at 4000x g for 3 minutes, and the resulting supernatant was discarded. The pellet was washed with 2 mL of dd H_2_O, vortexed, centrifuged, and the supernatant discarded. Then 3 mL of 1:1:1 chloroform : ethyl acetate : ethanol was added to the samples and vortexed vigorously to ensure re-distribution of the melanin. The tubes were then centrifuged at 4,000x g for 1 minute, and any resulting clear layers (top and or bottom) were discarded, leaving behind the dark layer. 2 mL of water was added to the sample for washing, and the tubes were centrifuged at 4,000x g for 1 minute, and the entire supernatant was discarded. Finally, 1 mL of 2M NaOH was added to each sample to allow for a standard volume added even if the melanin amount and therefore the final volume varied.

Extracted melanin samples suspended in 1 mL of 2M NaOH were then diluted 5 μL into 195 μL of 2M NaOH into a 96-well plate, with a 200 μL 2M NaOH blank well. These diluted samples were then read using the BioTek Synergy H1 hybrid spectrophotometer. The settings were for a full spectrum read from 230 nm to 700 nm, with 10 nm steps. However, the machine could not read OD values above 4.0, and therefore only data from 300 nm to 700 nm was used.

#### Confirming Active Melanin Excretion from Living Cells and Determining its OD Amount in the Supernatant

To confirm that *E. viscosa* JF 03-3F and *E. viscosa* JF 03-4F were actively excreting melanin, as opposed to dead cells lysing and releasing it, we grew up both strains and took daily aliquots for melanin extraction. We use the term excrete/excreting instead of secrete/secreting for the terminology of melanin being released into the media by the fungi because we have no evidence currently that this melanin release mechanism is an active process (i.e.: via Golgi, efflux pumps, or autophagy) warranting active secretion. Additionally, we wanted to compare the melanin excretion capabilities of these strains to *E. dermatitidis* for a baseline comparison. All three fungi were grown up in liquid MEA shaking at room temperature for 5 days. Then 2 mL of cells were washed with water three times. 500 μL of washed cells were then inoculated into 100 mL of MEA and YPD in 500 mL flasks. We let the cells grow at 23 °C shaking at 200 rpm for 7 days, removing 11 mL of cells and supernatant daily and pipetting them into 15 mL centrifuge tubes. The tubes of cells were then centrifuged at 3000x g for 5 minutes, the supernatant was removed, filter sterilized through a 2 μm filter, and placed into a new tube. We filter sterilized the supernatant to ensure that no cells remained in the supernatant, therefore all of the melanin extracted came only from excreted melanin. Melanin was then extracted using the chemical method explained above. Resulting pure melanin for all samples was read with the full spectrum as stated above, and both standard OD and log scale graph were created to confirm the presence of melanin with the proper R^2^ value above 0.9 (Pralea et al., 2019).

### Albino Mutant Creation for Genetic Analysis of Melanin Production

#### Creation of EMS Mutants and Annotation of their Mutations

Genetic modification of *E. viscosa* has not been established yet. Therefore, random mutagenesis was performed using ethyl methanesulfonate (EMS) to induce G:C to A:T mutations EMS mutagenesis was performed using the method by Winston (2008). Albino mutants and other interesting pigmentation or morphological mutants were isolated from the resulting mutagenesis, and their DNA was extracted using the CTAB DNA extraction protocol described above. DNA was then sent to the former Microbial Genome Sequencing Center (MiGS; https://www.migscenter.com/; 355 Fifth Avenue, Suite 810 Pittsburgh, PA 15222, now called SeqCenter (https://www.seqcenter.com/ ; 91 43rd Street, Ste. 250 Pittsburgh, PA 15201) for genome re-sequencing and base-called against the wild-type genome sequences. Resulting mutations were then manually annotated using the JGI Mycocosm genome “search” tool to identify affected genes.

#### Recovery of Melanin Production in Albino Mutants: Modern use of the Classical Beadle and Tatum Biochemical-Genetics Experiment

Following recovery of an albino mutant, we attempted to restore melanin production via chemical induction of the other melanin biosynthetic pathways. We did this using hydroxyurea (HU), L-DOPA, 1,8-DHN, and tyrosine. Hydroxyurea has been shown to enhance melanin production in *E. dermatitidis*, and L-DOPA is needed for certain fungi to produce melanized structures, including the albino mutant form of *E. dermatitidis* WdPKS1 (Dadachova et al., 2007; Paolo et al., 2006; Schultzhaus et al., 2020). Both YPD and MEA medium was made up and 20 mM of HU, 1 mM of L-DOPA, or 1 mM of 1,8-DHN was added to the medium after autoclaving. 5 μL of albino and wildtype control cells were spotted onto these media with added compounds, and they were grown for 10 days at 23 °C 12 hr light/dark cycle.

Additionally, we also tested the presence of tyrosine, the starting compound for L-DOPA and HGA melanins on the albino mutant. Growth of 5 μL spots of EMS 2-11 was grown on "Tyrosine media” (Jalmi et al., 2012) (2% w/v dextrose, 1% w/v peptone, 0.1% w/v yeast extract (Table 1)): without added tyrosine, with 0.5 g/L tyrosine, with 1 g/L tyrosine, and MN-nitrate 2% w/v dextrose with 0.5 g/L tyrosine as the sole nitrogen source. Colonies were allowed to grow for 19 days and then observations were made to see if melanization was recovered by different tyrosine levels.

### Data Availability

All data generated is available through the JGI Mycocosm website under *E. viscosa* JF 03-3F and *E. viscosa* JF 03-4F, or at NCBI under accession numbers JALDXI000000000 for *E. viscosa* JF 03-3F and JALDXH000000000 for *E. viscosa* JF 03-4F. Transcriptomes for these fungi can be found at NCBI using the SRA numbers: SRX5076261 for *E. viscosa* JF 03-3F, and SRX5077344 for *E. viscosa* JF 03-4F. All accession numbers from all databases for both strains are listed in Table 5. The strains of *E. viscosa* JF 03-3F CBS 148801 and *Exophiala viscosa* JF 03-4F CBS 148802 have also been deposited to the Westerdijk institute in a metabolically inactive state for access to the scientific community, but all strains used in this study can also be obtained from the corresponding author upon request. Finally, if any additional details or questions are required for the replication of any experiments listed above, one may reach out to the corresponding author.

**Table 5:** Accession numbers for all databases containing E. viscosa JF 03-3F and JF 03-4F’s data.

| **Database ID numbers** | ***Exophiala viscosa* JF 03-3F** | ***Exophiala viscosa* JF 03-4F** |
| --- | --- | --- |
| **WGS Accession GenBank** | JALDXI000000000 | JALDXH000000000 |
| **WGS Accession version used** | JALDXI010000001.1 | JALDXH000000001.1 |
| **Transcriptome Accession** | PRJNA501636 | PRJNA501637 |
| **BioProject** | PRJNA500555 | PRJNA500554 |
| **BioSample** | SAMN10366861 | SAMN10366859 |
| **SRA** | SRS4089839 | SRS4090708 |
| **DOE JGI GOLD project ID** | Gp0220461 | Gp0220460 |
| **CBS strain number** | 148801 | 148802 |
| **MycoBank number** | MB845198 | MB845199 |

## Results

### Fungal Isolation, ITS Alignment, and General Morphological Observations

Two fungi were isolated from BSCs located near Jackman Flats Provincial Park in B.C., Canada (Latitude: 52.94997°, Longitude: -119.41919°, Altitude: 738). Biological soil crust samples were diluted in water, crushed with a plastic micropestle, and plated out onto MEA with antibiotics and cycloheximide, and grown at 23 °C for 2 weeks to obtain the isolates. Once colonies began to form, they were picked and struck out onto regular MEA to obtain pure cultures. DNA extractions were performed on liquid cultures of the isolates that had been grown in liquid MEA for 5 days at 23 °C and 160 rpm of shaking. PCR amplification of the isolates’ ITS region was performed and then sequenced using Sanger sequencing. The resulting ITS sequences placed the fungal isolates within the genus *Exophiala*, and later apparent sequencing errors created nucleotide differences in their ITS sequences which initially placed them as different species. Further whole genome sequencing using Illumina based sequencing allowed for more accurate sequences of the ITS region for these isolates.

Using the Illumina-based ITS sequences of the isolates, it was determined that they have 100% identity to each other’s ITS sequence (Supplementary File 1). BLAST results to their ITS sequences using the UNITE simple analysis (Nilsson et al., 2019) resulted in a 97.57% identity to “*Exophiala nigra* strain CBS 535.95” accession number: MH862481.1, which was performed on September 29^th^, 2022. UNITE’s fungal species identification delineation typically separates species on ITS sequence similarity with ≥98% as the cut-off (Kõljalg et al., 2013), indicating that via ITS sequence alone, our *Exophiala* isolates (introduced here as *E. viscosa* JF 03-3F and JF 03-4F) were a separate species from *E. nigra*. However, since that date, the UNITE database has been further curated with more strict parameters and now only contains 19 *Exophiala* species, removing *E. nigra* and *E. sideris* as species in Taxon name searches. Repeating this analysis on December 12^th^, 2022, resulted in a BLAST result of *Capronia acutiseta* UDB035414 86.16% identity as the highest match.

Although both *E. viscosa* JF 03-3F and *E. viscosa* JF 03-4F are two strains of the same species indicated via their marker gene alignment, their cellular morphology and budding patterns differ drastically as described below. Both strains form smooth, shiny, sooty black colonies on PDA after 9 days of growth at 23 °C. When grown on high organic nitrogen media such as YPD, they will begin to excrete melanin into the agar by day 7 when grown at 23 °C, and by day 3 in liquid YPD (Figures 4L, 5L, & 14). On MEA, they are greyish brown in color, have the consistency of sticky black tar (Videos S1 & S2), and an iridescent rainbow shine which forms on their colonies after a week at 23 °C (Figures 4K, 5K & S3). These fungi also fail to easily disperse when suspended in water but can be readily pelleted. These observations and the prominence of lipid bodies within the cells (Figure S4) suggested that lipid-derived compounds could cause their sticky, water repelling, iridescent nature. These isolates grow to their maximum density in 7 days at 23 °C in 25 mL of MEA in a 250 mL Erlenmeyer flask shaken at 160 rpm. Both *E. viscosa* JF 03-3F and *E. viscosa* JF 03-4F have been preserved in a metabolically inactive state at the Westerdijk institute.

#### Phylogenetic Placement of Exophiala viscosa Amongst the Genus Exophiala

Phylogenetic analyses were performed using the marker genes 18S rDNA (1,600 bp), 28S rDNA (548 bp), and ITS rDNA (547 bp), with a combined sequence length of 2,685 base pairs. These genes were chosen because while they are predominant marker genes, they are also the only genes available on NCBI for the species *Phaeoannellomyces elegans*, which has been identified as closely related to this clade regardless of its nomenclature (Moussa et al., 2017). Bayesian analysis of the multi-locus gene tree yielded the best scoring tree, with the final log likelihood value of -9,031.1. Estimated base frequencies were: A = 0.242, C = 0.23, G = 0.275, and T = 0.253; with substitution rates of: AC = 0.161, AG = 0.151, AT = 0.143, CG = 0.067, CT = 0.412, GT = 0.065. The shape parameter of the gamma distribution of rate variation alpha was 0.141, and the total tree length was 0.574. The final average standard deviation of split frequencies at the end of the total 500,000 MCMC generations was 0.002955.

The multi-locus combined Bayesian inference and Maximum likelihood tree placed *E. viscosa* within the monophyletic clade composed of *E. nigra*, *E. sideris*, *E. bergeri*, and *P. elegans* with a posterior probability of 100% (Figure 2). In this combined Bayesian and Maximum likelihood tree, *P. elegans* is placed with 100% posterior probability and a bootstrap value of 100 as related to *E. sideris* and distinct from *E. nigra*. Additionally, *E. nigra* is further separated from *E. viscosa*, placing *E. nigra* on its own node beneath *E. bergeri* with 100% posterior probability and 100 bootstrap value (Figure 2). Both strains of the new fungus *E. viscosa* form a monophyletic node separate from *E. nigra*, *E. sideris*, *E. bergeri,* and *P. elegans* with a posterior probability of 100%, providing supporting that these strains are a novel species.

**Figure 2:**
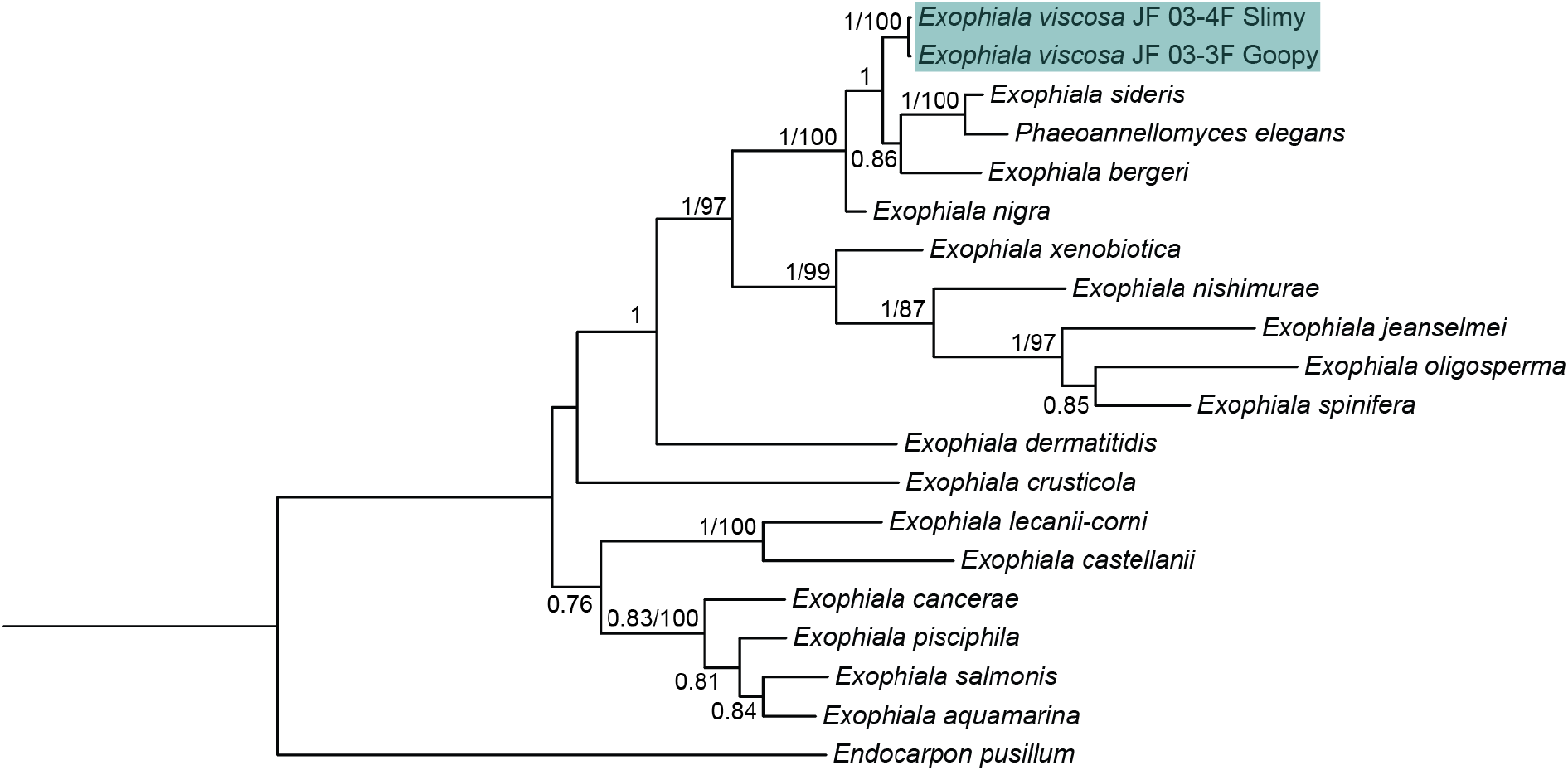
Rooted Bayesian phylogenetic tree of the concatenation of the 18S rDNA, 28S rDNA, and ITS rDNA regions of available *Exophiala* species with the addition of *E. viscosa* JF 03-3F and *E. viscosa* JF 03- 4F to identify the phylogenetic location of these new species within the genus *Exophiala*. The tree with the highest log likelihood is shown. The numbers above the branches are the Bayesian posterior probability/maximum likelihood bootstrap values, with posterior probabilities > 75% and bootstrap values > 80 shown. Location of *E. viscosa* JF 03-3F and *E. viscosa* JF 03-4F within *Exophiala* places them closest to *E. bergeri, P. elegans, and E. sideris*.

One additional marker gene sequence is available for *E. nigra* which is not available for *P. elegans*, and that is *RPB1*. When *RPB1* (464 bp) is added to the gene concatenation, creating the combination of 18S, 28S, ITS, and *RPB1* (3,156 bp total), *E. nigra* is again placed on its own node below *E. bergeri* maintaining a separate position to *E. viscosa* and the *E. sideris* clade with 100% posterior probability and 79 bootstrap value (Figure S2). This tree also shows *E. sideris* as the closest relative to *E. viscosa* with 100% posterior probability and a bootstrap value of 100. Within the three gene concatenation (18S, 28S, and ITS), the percent identical base pairs between *E. viscosa* and *E. sideris* are 98.436%, and 99.07% identical bases between *E. viscosa* and *E. nigra*. However, when *RPB1* is added to the concatenation, the percent identical bases between *E. sideris* and *E. viscosa* remain similar at 98.551%; whereas percent identical bases between *E. viscosa* and *E. nigra* becomes 97.03%. Percent identity matches between all species represented in the phylogenetic trees of Figure 2 and Figure S2 across all genes and gene concatenations are provided in Supplemental File 1. Further phylogenetic analyses were performed on all *Exophiala* species and related fungi who have been whole genome sequenced, and whose genomes are available in JGI’s Mycocosm. With their whole genomes, we were able to perform single-copy homologous protein sequence approximate maximum likelihood phylogenetic analyses, providing additional support for *E. viscosa*’s intraspecific variation from specifically the fungus *E. sideris*. Our approximate maximum likelihood single-copy homologous protein phylogenetic tree shows with 100 bootstrap value, that *E. sideris* is separate from *E. viscosa* (Figure 3).

**Figure 3:**
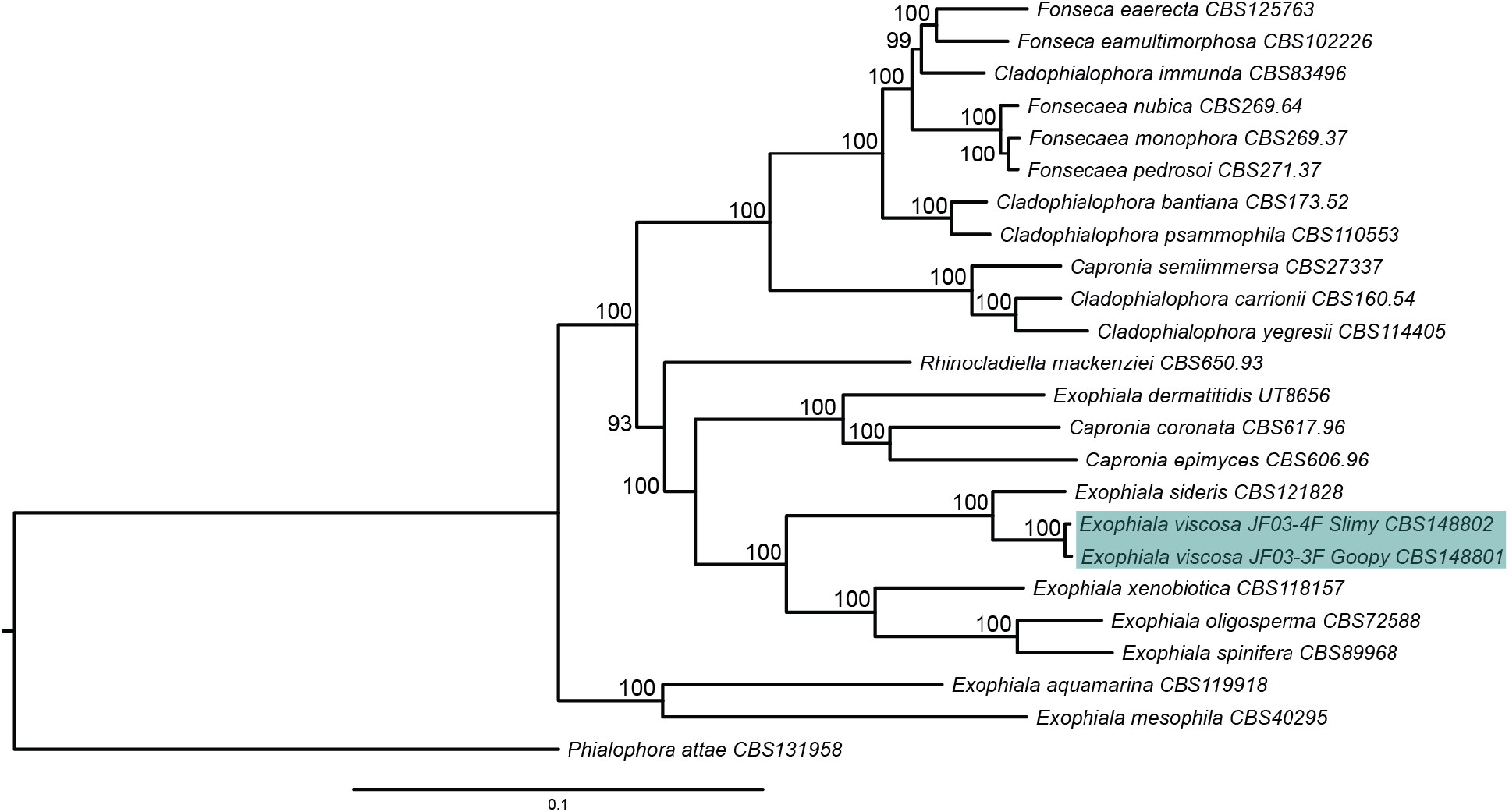
Single-copy homologous protein sequence approximate maximum likelihood phylogenetic tree of taxa of the family Herpotrichielleaceae all of which have been whole genome sequenced and annotated. *E. viscosa* JF 03-3F and *E. viscosa* JF 03-4F are shown to be most closely related to *E. sideris*. Node values represent bootstrap values.

ASAP values of the marker genes allowed us to confirm our phylogenetic inferences of separating *E. viscosa* from the other species in its clade. For the ITS-only ASAP analysis, the output provided an ASAP score of 1.0 to 18 distinct species with a distance threshold of 1.32% (Supplemental File 2). Both strains of *E. viscosa* were considered one species, and *E. sideris* and P. elegans were considered one species. When ASAP was performed on the concatenated 18S-28S-ITS alignment, it maintained that there should be 18 distinct species (ASAP score of 2.0), with both strains of *E. viscosa* as one species and *E. sideris* and *P. elegans* as one species. However, with the three-gene concatenation, the distance threshold decreased to 0.75%. Finally, the concatenation of 18S-28S-ITS-*RPB1* provided an ASAP score of 2.5 which suggested that there be 16 species with a distance threshold of 2.5%, combining both strains of *E. viscosa* into one species, and combining *E. aquamarina* and *E. pisciphila* into one species (*E. crusticola* and *P. elegans* are not in this dataset). All three ASAP analyses suggest that both strains of *E. viscosa* remain as one species, and that they are a distinct species from the other *Exophiala* spp.

#### Morphological Differences Between E. viscosa Strains JF 03-3F and JF 03-4F

While these two strains of *E. viscosa*, JF 03-3F and JF 03-4F, are very similar to each other in genomic content and have 100% similarity across the four marker genes 18S, 28S, ITS, and *RPB1*, they vary in gene synteny, they contain multiple single nucleotide polymorphisms across their genomes, and their morphologies differ. *E. viscosa* JF 03-3F is found primarily as a yeast morphology on all media tested, with ellipsoidal cells 4.4 μm ± 0.34 μm wide and 5.36 μm ± 0.37 μm long (n=20 cells measured; in MEA). *E. viscosa* JF 03-4F is found primarily in a pseudohyphal form on all medias tested, with more elongated cells 3.6 μm ± 0.38 μm wide and 6.0 μm ± 0.67 μm long (n=20 cells measured; in MEA). *E. viscosa* JF 03-3 will form pseudohyphae when grown for longer than 2 weeks and only on solid media, whereas *E. viscosa* JF 03-4F readily forms pseudohyphae in liquid media and on solid media as a primary growth from and will form a few true hyphae after 3 weeks of growth on MEA (Figures 5N, 6, & Video S3). As seen in Figures 6 & S3D, the cellular morphology of *E. viscosa* JF 03-4F resembles that of *Hortaea werneckii* as observed by Mitchison-Field et al. (2019). Budding of *E. viscosa* JF 03-3F cells occur at 180° or 90° from the mother cell’s origin, whereas JF 03-4F is almost exclusively budding 180° from the mother cell. When *E. viscosa* JF 03-4F is grown up to maximum density in liquid culture, pipetting becomes difficult due to large clumps of cells formed by its more filamentous growth pattern (Figures 5N & 6). However, *E. viscosa* JF 03-3F at maximum density in liquid culture does not form large clumps, and therefore creates no pipetting issues. Notably, *E. viscosa* JF 03-4F possesses a looser pellet and is less refractory to re-suspension than *E. viscosa* JF 03-3F. *E. viscosa* JF 03-4F also produces a fuller, thicker biofilm at the air-liquid interface than *E. viscosa* JF 03-3F, as seen in Figures 4I & 5I. *E. viscosa* JF 03-3F is also a much more prolific melanin excreter than *E. viscosa* JF 03-4F (Figures 14, 15, & 16).

**Figure 4:**
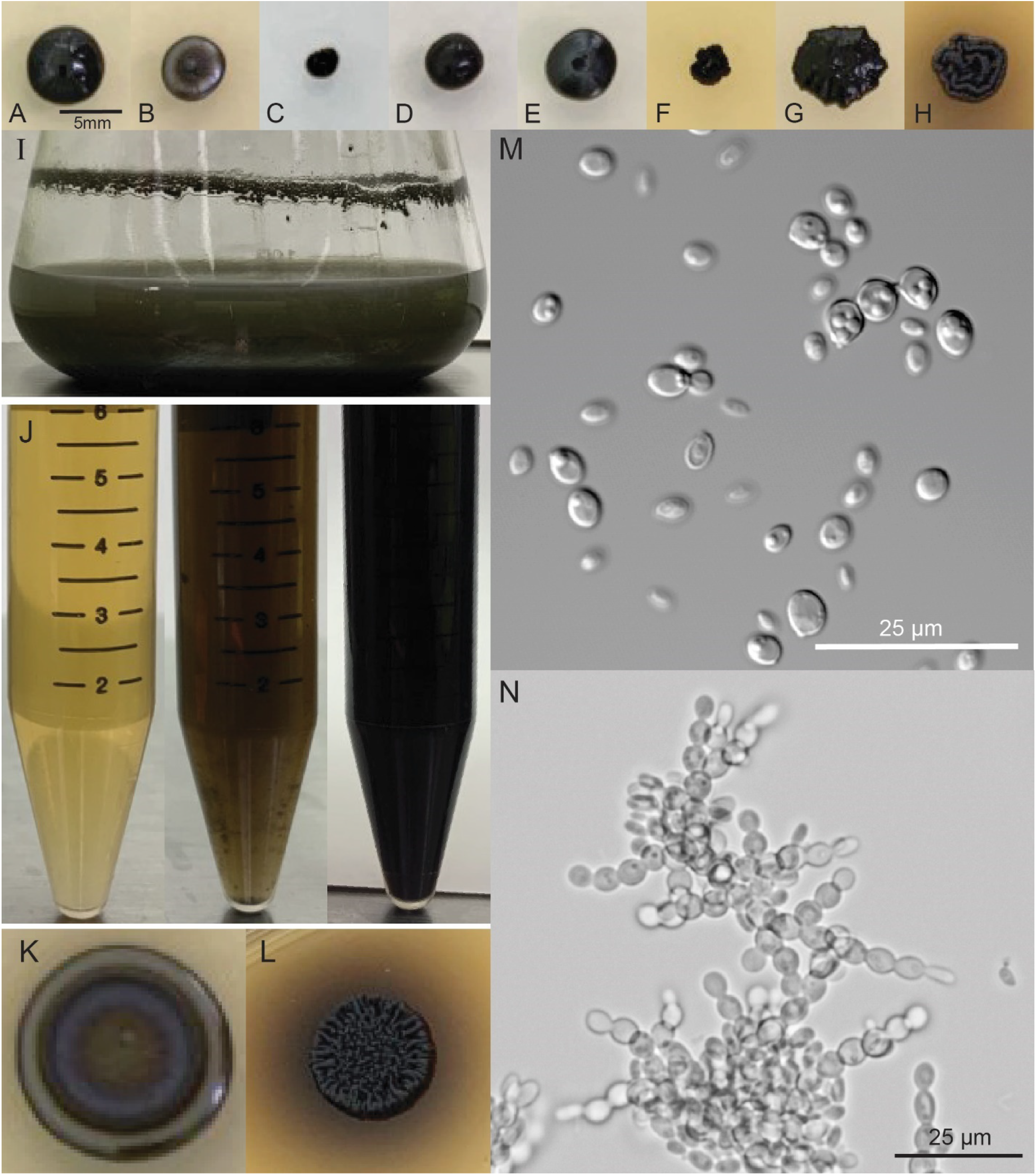
*E. viscosa* JF 03-3F’s description summary figure. All colonies: day 14 grown at 23° C) A) Colony on MAG; B) Colony on MEA; C) Colony on MN-NAG; D) Colony on MNV; E) Colony on PDA; F) Colony on Spider media; G) Colony on V8; H) Colony on YPD; I) Biofilm at air-liquid Interface of growth in MEA day 3; J) Supernatant when grown in YPD; Day 0, 3, 6; K) Colony growth from 5 µL of liquid culture on MEA after 14 days with rainbow sheen; L) Colony growth from 5 µL of liquid culture on YPD after 14 days with dark excretions; M) Cellular growth in liquid MEA, shaking at room temp. day 5; N) Cellular growth on solid MEA, room temp. hour 65, using VALAP method.

**Figure 5:**
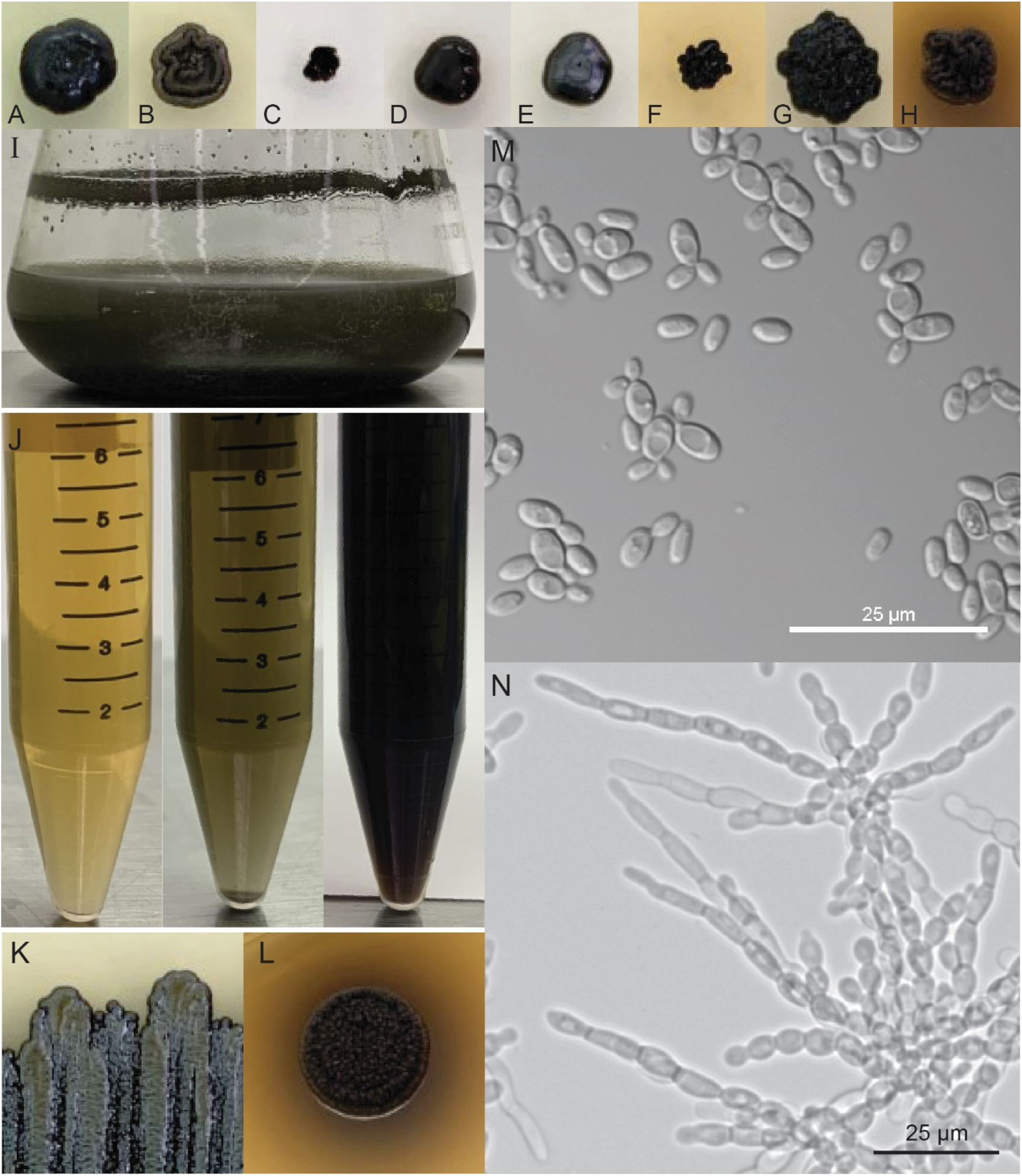
*E. viscosa* JF 03-4F’s description summary figure. All colonies: day 14 grown at 23° C A) Colony on MAG; B) Colony on MEA; C) Colony on MN-NAG; D) Colony on MNV; E) Colony on PDA; F) Colony on Spider; G) Colony on V8; H) Colony on YPD; I) Biofilm at air-liquid interface of growth in MEA day 3; J) Supernatant when grown in YPD; Day 0, 3, 6; K) Colony growth from 5 µL of liquid culture on MEA after 14 days with rainbow sheen; L) Colony growth from 5 µL of liquid culture on YPD after 14 days with dark excretions; M) Cellular growth in liquid MEA, shaking at room temp. day 5; N) Cellular growth on solid MEA, room temp. hour 60, using VALAP method.

**Figure 6:**
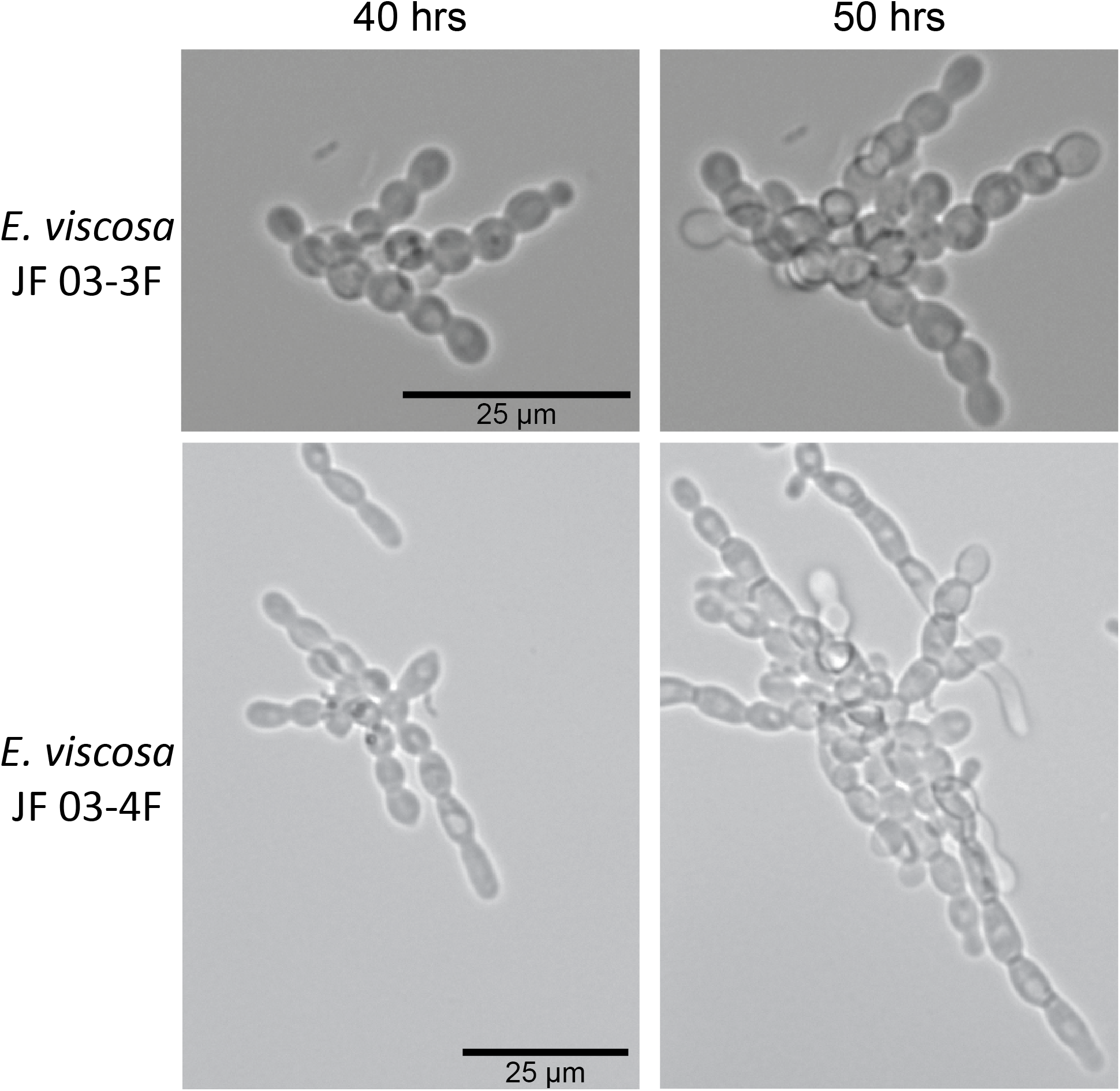
Budding styles of *E. viscosa* JF 03-3F and *E. viscosa* JF 03-4F pseudohyphae. Rate of budding is higher in *E. viscosa* JF 03-4F than in *E. viscosa* JF 03-3F, as seen at the 50 hrs mark. Budding style is also different between the species, *E. viscosa* JF 03-3F buds both distal polarly and proximal at close to a 90° angle, whereas *E. viscosa* JF 03-4F buds almost exclusively as a distal polar. *E. viscosa* JF 03-4F’s cells also elongate with every bud, forming almost hyphal-like structures at the 50 hr timepoint (scale bars apply to side-by-side images).

#### Recognition of Novel Taxon Exophiala viscosa

The new species *Exophiala viscosa* is defined based on molecular phylogenetic analyses of the four marker regions: 18S, 28S, ITS, and *RPB1* resulting in the holotype and the additional strain forming one monophyletic node separate from *E. sideris, P. elegans, E. bergeri* and *E. nigra* which are its closest relatives, with the genes ITS and *RPB1* providing the most support for intergenic variation (Figures 2 & S2). Additional phylogenetic support results from the single-copy gene tree of whole genome sequenced *Exophiala* spp. and related fungi, which further supports intraspecific variation of *E. viscosa* from *E. sideris* (Figure 3). Whole genome sequencing and then single-copy gene phylogenetic analyses of *E. nigra* and *P. elegans* would have been beneficial for further elucidation of this fungus, but that was not feasible at this time. Additionally, ASAP values of all three gene/gene concatenations tested were in consensus that *E. viscosa* is a separate species from *E. sideris*, *E. bergeri*, *P. elegans*, and *E. nigra*.

Morphological comparison of *E. viscosa* to *E. nigra* from images in Moussa et al. show differences in colony and cell morphology (2017). Colonies of *E. nigra* on MEA showed no rainbow sheen and remained olivaceous black, with irregular undulate colony edges (Moussa et al., 2017). Whereas *E. viscosa*’s colonies are smooth to curled, are dark grayish brown in color, and have a rainbow sheen. *E. nigra* colonies on PDA were also irregular and undulate on the edges and umbonate or lumpy on the elevation morphology (Moussa et al., 2017), whereas *E. viscosa* has smooth and circular edges with a flat elevation on PDA (Figures 4E & 5E). Cellular morphology images of *E. nigra* on MEA showed pseudohyphae to hyphal formation with conidia forming on the tips of hyphae (Moussa et al., 2017). Pseudohyphae are observed in *E. viscosa* on MEA, but hyphae are rarely observed except in the strain JF 03-4F, and no conidia were observed forming at the tips of pseudohyphae or hyphae on MEA even after three weeks. No characteristic asexual or sexual structures of *Exophiala* species were observed in either strain on any medias or conditions tested on *E. viscosa*.

The other closest relative of *E. viscosa* is *E. sideris*. Multiple strains of *E. sideris* were isolated from primarily arsenate and other metal-contaminated locations, and the holotype strain is isolated from *Sorbus aucuparia* berries in The Netherlands (Seyedmousavi et al., 2011). These specific locations and environments are distinct from those in which *E. viscosa* has so far been recovered. The differences between a BSC environment and a berry or rock surface would be the permanence of the location (in reference to the berry surface) and the abundance and make-up of microbial neighbors (i.e.: cyanobacteria, algae, specific lichens, and bacteria) (Couradeau et al., 2019; Nunes da Rocha et al., 2015). Additionally, while arsenate was not a metal tested on *E. viscosa*, the other metals tested on both strains of this fungus did not indicate increased metal resistance, and showed less resistance to CoCl_2_, AgNO_3_, and CuCl_2_ than *S. cerevisiae* (Table 9). Neither sexual nor asexual structures were observed in *E. viscosa* on all medias and conditions tested including on PDA after 30 days, whereas *E. sideris* readily forms conidia on PDA (Seyedmousavi et al., 2011). Attempts at inducing asexual or sexual development for both *E. viscosa* strains were performed using established methods: Modified Leonian’s agar from Untereiner (1994), Untereiner & Naveau (1999), and Potato dextrose agar via methods from Seyedmousavi et al. (2011), but were not successful.

The isolates described here are the first described instance of this fungus being cultured. This is only the second instance of an *Exophiala* species being isolated from a BSC, the first being *E. crusticola* (Bates et al., 2006). The ITS sequences of *E. viscosa* and *E. crusticola* have 82.904% identity (Supplemental File 1), and therefore they are not the same species, which is supported by the phylogenetic tree in Figure 2. It is also the first instance of an *Exophiala* species being isolated from a BSC in Canada, as opposed to the Western United States where *E. crusticola* was isolated. In terms of other instances of *E. viscosa*’s presence in biological samples, according to BLAST results of *E. viscosa*’s ITS region on NCBI there has been one instance of another 100% identical fungal DNA from a rocks surface with the accession number AY843177. Unfortunately, this fungal DNA is from an unpublished article according to NCBI. Additional fungal ITS ribosomal DNA that closely matched *E. viscosa*’s via BLAST on NCBI with 99.26% identity were: MZ017038, MZ016619, MZ016618, all of which were DNA samples labeled “Fungal sp.” from *Pinus ponderosa* either roots or needles from Arizona (USA) (Bowman & Arnold, 2021).

#### Taxonomy and Species Descriptions of E. viscosa

*Exophiala viscosa* JF 03-3F Goopy E.C. Carr, S.D. Harris, **sp. nov.**

MycoBank MB845198

**Figure 4**

*Typification:* CANADA. BRITISH COLUMBIA: Adjacent to Fraser-Fort George H, Jackman Flats Provincial Park (Lat.: 52.95; Long.: -119.416; Alt.: 783), in biological soil crust sample, sampled June 2015, *S. Harris*, isolated July 2015, *E. Carr*. Holotype culture strain JF 03-3F Goopy: CBS 148801. Additional strains JF 03-4F Slimy: CBS 148802.

*GenBank*: Whole genome = JALDXI000000000.1

NCBI Taxonomy ID: 2486360

*Etymology*: *Exophiala viscosa* was colloquially referred to as “Goopy” due to the nature of its yeast morphology lending to a very goopy colony. Accordingly, we have formally named it *Exophiala viscosa* for the Latin term of viscous.

#### Sexual and asexual morph: not observed

##### Culture characteristics

Hyphae not observed in liquid culture, aerial pseudohyphae in colonies observed by day 30 in Malt Extract medium (MEA), primarily found in yeast form in liquid MEA and Yeast Peptone Dextrose (YPD) mediums (23 °C) but will form pseudohyphae as well. Cells in MEA liquid culture are ellipsoid in shape, 3.9 μm ± 0.55 μm wide and 5.66 μm ± 0.62 μm long (n=40 cells measured). Cell walls start off thin and become thicker over time. In pseudohyphal form on solid MEA, budding occurs at 180° or 90° from the mother cell’s origin, or at acute angles from the previous bud site. Cells either maintain size and shape of yeast form as pseudophyphal cells bud out for four buds then begin to elongate, or the buds begin to elongate from the first bud and continue to elongate with each linear budding. Colony morphology all at 23 °C with a 12 hour light/dark cycle: on MEA colonies are initially shiny dark grayish brown and smooth to curled, then develops a rainbow sheen with a colony size of 5.2- 6.6 mm by day 14, additionally dark supernatant is observed in liquid MEA culture starting on day 3; on Malt Extract Agar with vitamins and trace elements (MAG) colonies are sooty black and glossy with smooth colony edges forming a dome shape, and 6.3-8.1 mm growth is observed on day 14; on Minimal media (MN; nitrate with salts) + N-acetyl Glucosamine as the carbon source, the colony is matte sooty black smooth edged and grows to 2.7-2.8 mm in 14 days; on MN with vitamins added (MNV), the colony is glossy raisin black with smooth edges and smooth top, colonies grow to 4.7-5.3 mm in 14 days; on Potato Dextrose Agar (PDA) colonies are glossy sooty black with smooth edges and grow to 5.6-5.8 mm in 14 days; on Spider media, the colonies are sooty black and irregularly shaped with a size of 3.0-4.5 mm in 14 days, additionally a red pigment Is observed surrounding the colonies in the media; on V8 medium colonies are glossy sooty black and irregularly shaped, growth reaches 7.5-8.1 mm in 14 days; on YPD colonies are blackish brown (1) and matte with round to irregular side morphologies and undulate top morphology, dark excretions are observed coming from the colonies and excessively dark supernatant is observed in liquid culture starting on day 2, colony growth reaches 6.1-6.2 mm in 14 days. Color descriptions used above come from Ridgway (1912).

##### Cardinal temperatures

Growth is capable at temperatures from 4 °C to 28 °C, and optimal growth is at 23 °C, growth is not capable at or above 37 °C.

##### Additional specimen examined

CANADA. BRITISH COLUMBIA: Fraser-Fort George H, Jackman Flats Provincial Park, in biological soil crust sample, sampled June 2015, *S. Harris*, isolated July 2015, *E. Carr*. Culture *Exophiala viscosa* Slimy JF 03-4F CBS 148802.

*Notes*: Support for *E. viscosa* as a novel species stems from molecular multi-locus phylogenetic analyses both for marker genes and single-copy genes, morphological comparisons of micro- and macromorphology to phylogenetically related species, ecology, and biogeography of the novel species. *E. viscosa* is phylogenetically placed on all analyses as related to but separate from *E. sideris* with 100% posterior probability in all Bayesian analyses, and bootstrap values above 95 in all Maximum likelihood analyses (Figures 2, 3, & S2). The highest matching species to *E. viscosa*’s ITS sequence was *E. nigra*, however, marker gene phylogenies place it as polyphyletic to *E. viscosa* (Figures 2 & S2). Morphologically, *E. viscosa* is distinct in that it excreted melanin into its surrounding environment, especially on high organic nitrogen substrates such as YPD or on MEA if grown for longer than 2 weeks, which is not noted in other *Exophiala* species. Additionally, it forms a rainbow sheen when grown on MEA, which is not observed in *E. sideris* or *E. nigra* (Moussa et al., 2017; Seyedmousavi et al., 2011). Sexual or asexual structures were not observed in *E. viscosa* on all medias and conditions tested, whereas *E. sideris* readily forms conidia on PDA (Seyedmousavi et al., 2011). Both strains of *E. viscosa* are also the first instances of *Exophiala* species being isolated from Canadian soils and biological soil crusts, and the only other *Exophiala* species that has been previously isolated from a biological soil crust was *E. crusticola* (Bates et al., 2006).

#### Genome Description and Gene Content

Genome sequencing, assembly, and annotation was performed by DOE JGI and their genomes and transcriptomes are deposited on the Mycocosm website under their genus species names. All accession numbers for both strains are listed in Table 5. The genome assembly sizes of these two novel *Exophiala* strains are: 28.29 Mbp for *E. viscosa* JF 03-3F and 28.23 Mbp for *E. viscosa* JF 03-4F (Table 6). These sizes are similar to other yeasts in the genus *Exophiala* and are only a bit smaller than their closest relative *E. sideris* (29.51 Mbp). Their predicted gene contents are 11,344 for *E. viscosa* JF 03-3F and 11,358 for *E. viscosa* JF 03-4F (Table 6). *E. sideris* appears to possess longer genes, transcripts, exons, and introns compared to *E. viscosa* JF 03-4F and *E. viscosa* JF 03-3F, which could also contribute to the gene number to genome size differences (Table 6). *E. viscosa* JF 03-3F contains the highest GC% amongst the *Exophiala* species listed in Table 6. *E. viscosa* JF 03-3F’s GC content is even higher than *E. viscosa* JF 03-4F by 2.65%.

**Table 6:** Genome descriptions of the novel Exophiala species and related taxa. Information gathered from JGI’s Mycocosm.

|  | *E. viscosa* JF 03-3F<br>CBS148801 | *E. viscosa* JF 03-4F<br>CBS148802 | *E. sideris*<br>CBS121828 | *E. spinifera*<br>CBS89968 | *E. xenobiotica*<br>CBS118157 | *E. oligosperma*<br>CBS72588 | *E. dermatitidis*<br>UT8565 |
| --- | --- | --- | --- | --- | --- | --- | --- |
| **Accession #s** | PRJNA500555 | PRJNA500554 | PRJNA325799 | PRJNA325800 | PRJNA325801 | PRJNA325798 | PRJNA225511 |
| **Genome Assembly statistics** |  |  |  |  |  |  |  |
| **Genome Assembly size (Mbp)** | 28.29 | 28.23 | 29.51 | 32.91 | 31.41 | 38.22 | 26.35 |
| **Number of contigs** | 35 | 27 | 69 | 143 | 64 | 287 | 10 |
| **Number of scaffolds** | 30 | 17 | 5 | 28 | 15 | 143 | 10 |
| **Number of scaffolds >= 2Kbp** | 25 | 17 | 5 | 20 | 11 | 129 | 10 |
| **Scaffold N50** | 4 | 5 | 2 | 4 | 3 | 5 | 4 |
| **Scaffold L50 (Mbp)** | 2.24 | 2.89 | 7.9 | 3.79 | 5.04 | 3.39 | 3.62 |
| **Number of gaps** | 5 | 10 | 64 | 115 | 49 | 144 | 0 |
| **% of scaffold length in gaps** | 0.00% | 0.00% | 0.10% | 0.10% | 0.10% | 0.80% | 0.00% |
| **Three largest Scaffolds (Mbp)** | 5.39, 4.56, 2.49 | 4.57, 3.58, 2.93 | 9.94, 7.90, 7.15 | 6.18, 3.93, 3.92 | 5.55, 5.20, 5.04 | 4.47, 4.29, 4.12 | 4.25, 4.22, 3.71 |
| **GC content (%)** | 51.91 | 49.26 | 49.73 | 49.42 | 51.89 | 50.99 | 51.74 |
| **Gene statistics** |  |  |  |  |  |  |  |
| **Number of genes** | 11344 | 11358 | 11120 | 12049 | 13187 | 13234 | 9562 |
| **Gene length (bp, Average)** | 1840 | 1844 | 2044 | 1593 | 2072 | 2090 | 2237 |
| Gene length<br>(bp, Median) | 1666 | 1671 | 1823 | 1415 | 1845 | 1879 | 1923 |
| Transcript<br>length (bp,<br>Average) | 1740 | 1746 | 1933 | 1483 | 1941 | 1949 | 2122 |
| Transcript<br>length (bp,<br>Median) | 1564 | 1574 | 1710 | 1311 | 1710 | 1735 | 1794 |
| Exon length<br>(bp, Average) | 705 | 707 | 776 | 621 | 738 | 743 | 896 |
| Exon length<br>(bp, Median) | 410 | 414 | 452 | 339 | 430 | 426 | 513 |
| Intron length<br>(bp, Average) | 70 | 69 | 74 | 81 | 80 | 85 | 85 |
| Intron length<br>(bp, Median) | 56 | 56 | 56 | 61 | 57 | 61 | 62 |
| Protein length<br>(aa, Average) | 488 | 489 | 500 | 494 | 492 | 488 | 501 |
| Protein length<br>(aa, Median) | 428 | 430 | 434 | 437 | 431 | 429 | 429 |
| Exons per gene<br>(Average) | 2.47 | 2.47 | 2.49 | 2.39 | 2.63 | 2.62 | 2.37 |
| Exons per gene<br>(Median) | 2 | 2 | 2 | 2 | 2 | 2 | 2 |

Although the morphology and development of *E*. *viscosa* is seemingly much less complex compared to other Eurotiomycete fungi (e.g., *Aspergillus nidulans*), there are few differences in the complement of genes that are assigned to these processes when these fungi are compared. For example, compared to *A*. *nidulans*, differences include losses of a velvet regulator (VosA) as well as a class of hydrophobins (RodA, DewA), along with the gains of additional cryptochrome/photolyases (CryA) and an NADPH oxidase (NoxB) (Supplemental File 3). This pattern suggests that *E*. *viscosa* might harbor additional morphological or developmental complexity that has yet to observed. Notably, results from automated and manual annotation of the genome sequences reveal no striking differences in numerous traits (e.g., morphogenetic or developmental functions, gene clusters associated with secondary metabolism, CAZymes, transcription factors, cytochrome p450 complement) compared to other sequenced and annotated genomes from the members of Chaetothyriales (Teixeira et al., 2017). Additional details of both species’ genome sequences, including results from automated annotation (e.g., GO, KEGG, PFAM domains, etc.) can be found on their respective pages on the JGI Mycocosm website (https://mycocosm.jgi.doe.gov/mycocosm/home).

#### Mating Type

*Exophiala* species are one of the many fungal genera whose mating capabilities remain incompletely understood and vary across the genus (Teixeira et al., 2017). The closest species to *E. viscosa* JF 03-3F and *E. viscosa* JF 03-4F with its mating type identified is *E. sideris,* within which both mating types have been characterized as a single fused gene (Teixeira et al., 2017). Given the known order of genes in regions flanking the *MAT* locus in *Exophiala* species, we used comparative approaches to determine the mating identities of the sequenced *E. viscosa* JF 03-3F and *E. viscosa* JF 03-4F isolates, and to look for possible evidence of homothallism that is also presumed in *E. sideris*. Homologues of the genes *APN2*, *SLA2*, *APC5*, *COX13*, and *MAT1* (*MAT1-1/alpha*) in *E. viscosa* JF 03-3F and *E. viscosa* JF 03-4F were identified via BLAST searches. We found that *APN2* and *SLA2* flank the *MAT* locus of both species, and they also contain the Herpotrichielleaceae-specific mating gene *MAT1-1-4* (Figure S5). These results are not surprising, in that this is the exact same order of these genes in *E. sideris*. Interestingly, as Teixeira et al. indicate, in *E. sideris* the *MAT* gene is a fused *MAT1-1/MAT1-2* gene with the HMG domain in the middle splitting the α-box in half (2017). When the MAT protein sequences of *E. viscosa* JF 03-3F and *E. viscosa* JF 03-4F are aligned with the *MAT* gene of *E. sideris*, we see 83.79% homology between their protein sequences indicating that these fungi also contain a fused *MAT* gene (Figure S5). We were able to confirm that the HMG domain was inserted within the α-box of *E. viscosa* via NCBI’s conserved domains visualizer, where residues 75 to 206 were the HMG domain within the *MAT1-1* protein sequence. This fused *MAT* gene (*MAT1-1-1*) presumably confers homothallic mating behavior, although this has not been confirmed experimentally in any of these three fungi. Additionally, neither *E. viscosa* JF 03-3F nor *E. viscosa* JF 03-4F contain an additional gene between *APN2* and the *MAT1-1-4* as is found in *E. sideris* (Teixeira et al., 2017). Furthermore, *COX13* and *APC5* are about 7,000 bp downstream of the MAT locus in both species, but *COX13* is on the reverse strand in *E. viscosa* JF 03-4F compared to being on the forward strand in *E. viscosa* JF 03-3F (Figure S5).

### Phenotyping of *E. viscosa* JF 03-3F and *E. viscosa* JF 03-4F

To further understand the morphology, growth capabilities, and stress responses of *E. viscosa* JF 03-3F and *E. viscosa* JF 03-4F, we performed multiple phenotyping experiments. The intent of these experiments was to provide a broader perspective on the potential adaptations that would support the ability of these fungi to successfully colonize BSCs and other extreme niches.

#### Budding Patterns

Due to similarities in their colony morphologies, the extent to which this was reflected in their respective budding patterns was assessed. Using protocols adapted from Mitchison-Field et al. (2019) we were able to perform more detailed time-lapse microscopy of their budding patterns (Figure S1). From this we were able to observe dramatic differences in the budding types of *E. viscosa* JF 03-3F and *E. viscosa* JF 03-4F.

*E. viscosa* JF 03-3F buds with round cells in initially a distal fashion where the new bud forms 180° from the mother cell origination site, but also forms new buds at a ∼90° angle from where the mother bud was formed in a proximal manner (Figure 6; Video S4). *E. viscosa* JF 03-4F on the other hand forms elongated cells in a distal manner, forming longer chains of cells instead of clusters, with axial buds forming at later timepoints (Figure 6; Video S2). These morphological differences in budding patterns influences the way these two strains grow in a shaking flask. For example, *E. viscosa* JF 03-4F forms more elongated cells and buds distally which although it does not result in the formation of hyphae, still creates larger clumps of cells which are not easily pipetted. However, because *E. viscosa* JF 03-3F forms rounder cells and buds with both distal and proximal patterns, it does not form extensive clumps in a shaking flask and is more easily pipetted. *E. viscosa* JF 03-4F also forms more extensive biofilms at the liquid-air interface than *E. viscosa* JF 03-3F, likely also due to the elongated cell morphology (Figures 4I & 5I).

#### Growth of E. viscosa JF 03-3F and E. viscosa JF 03-4F on Different Media

Growth of *E. viscosa* JF 03-3F and *E. viscosa* JF 03-4F on a variety of different media was done to evaluate growth requirements and their impact on pigmentation (Figures 7 & 8). The media used are described in Table 1, and include MAG, MEA, MN+NAG, MNV, PDA, Spider, YPD, YPD+TE, and V8. The addition of Vitamin mix (Table 1) to any medium, but specifically to MAG and MNV, caused the colonies of both isolates to become much shinier, blacker (vs. browner), and more yeast morphology. Growth on MEA caused the formation of a rainbow iridescent sheen (Figures 7, 8, 4K and 5K), which is not seen on any other medium. Spider medium and YPD caused the formation of a colored halo around colonies of both strains. However, the halo around colonies grown on YPD is much darker and extends further than on Spider medium, and *E. viscosa* JF 03-3F showed a more extensive halo than *E. viscosa* JF 03-4F. When trace elements (TE) are added to YPD, the formation of a dark halo is inhibited in both strains. The ability of both strains to grow on V8 medium implies that they can use cellulosic material as a carbon source. Overall, colony growth and pigmentation were similar across of media types for both strains (Figures 7 & 8).

**Figure 7:**
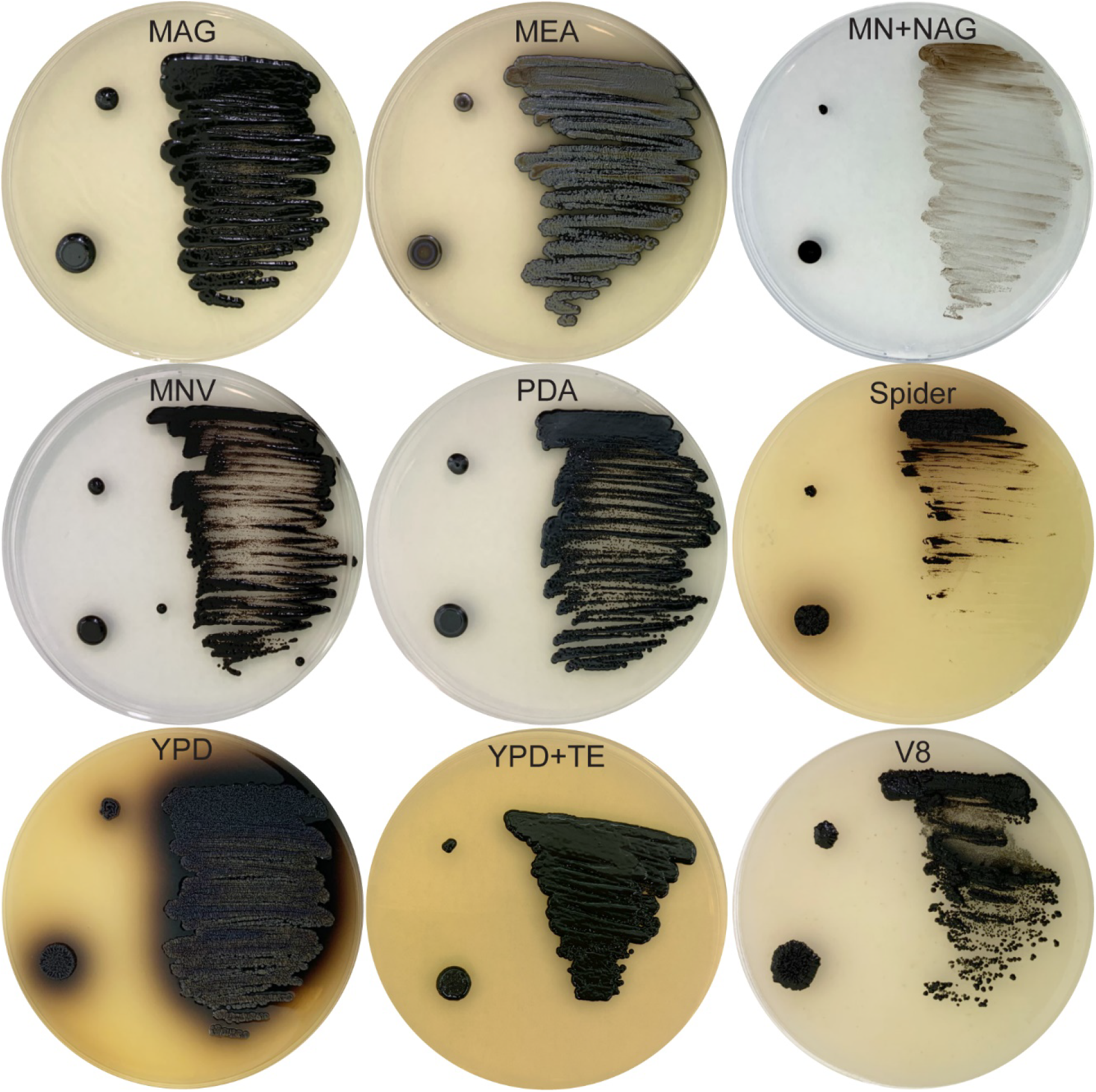
Growth of *E. viscosa* JF 03-3F on different media types grown at 22 °C for 14 days, plated as a toothpick prick (top left), 5 μL spot (bottom left), and loop streak patterns (pictures reduced in contrast and lightened to see details; see **Figure S15** for originals). *E. viscosa* JF 03-3Fwas capable of growth on all medias tested, but growth on MN+NAG showed the least amount of growth. PDA, MAG, and MNV allowed for very shiny and dark black colony growth. Growth on V8 medium shows potential for saprotrophic growth on plant material. Colorful red excretion was observed on Spider media and a lot of dark brownish-black excretion on YPD. Addition of trace elements (TE) to YPD prevents excretion of melanin into the media. Growth on MEA resulted in a rainbow iridescent shine on both the streaked-out growth and the 5 μL spot. Images adjusted for brightness and contrast for easier visualization, original images in **Figure S15**.

**Figure 8:**
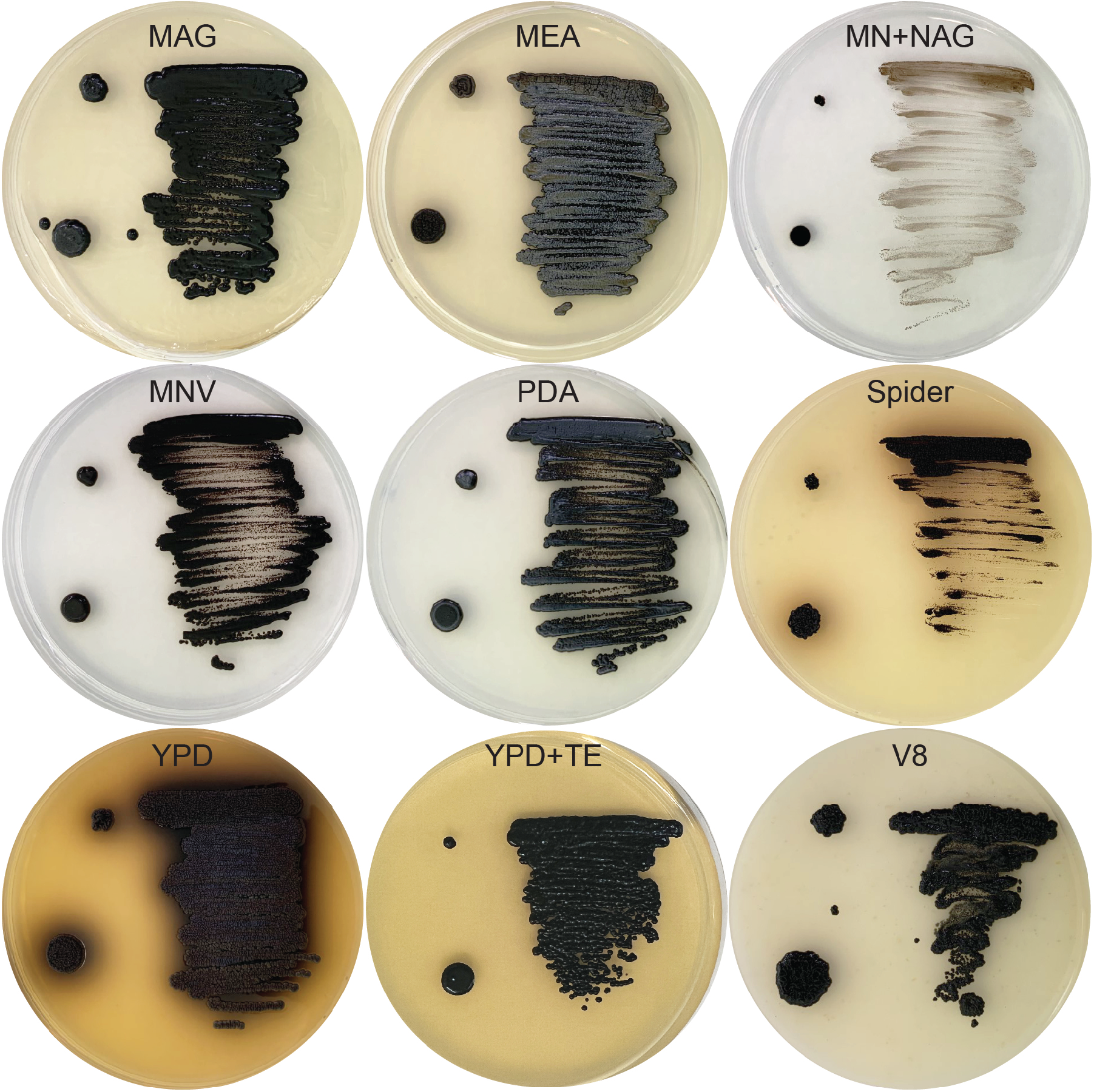
Growth of *E. viscosa* JF 03-4F on different media types grown at 22 °C for 14 days, plated as a toothpick prick (top left), 5 μL spot (bottom left), and loop streak patterns (pictures reduced in contrast and lightened to see details; see supplementary for originals). *E. viscosa* JF 03-4F was capable of growth on all medias tested, but growth on MN+NAG showed the least amount of growth. PDA, MAG, and MNV allowed for very shiny and dark colonies to form. Growth on V8 medium shows potential for saprotrophic growth. Colorful excretions were observed on both Spider media and YPD. Addition of trace elements (TE) to YPD prevents excretion of melanin into the media. Growth on MEA shows an iridescent shine on the streaked-out growth. Images adjusted for brightness and contrast for easier visualization, original images in **Figure S15**.

#### Optimal Wavelength for OD Measurements

Serial dilutions (1:1 to DF=512) of both *E. viscosa* JF 03-3F and *E. viscosa* JF 03-4F were read in a wavelength range of 300-700 nm at 10 nm intervals. The OD results of those dilutions were then compared to: the percent difference at 25% dilution and below, the percent difference at 50% and below, cell counts at 2.19x10^6^ cells/mL and below, cell counts at 4.38x10^6^ cells/mL and below, and the consistency of the actual percent difference at 12.5% dilution and below. The resulting R^2^ values showed that the optimal wavelength range for reading ODs for these strains is between 410-420nm (Supplemental Table 1). To reduce the potential effect pure melanin in the supernatant might be having on the OD results, we decided to choose the higher wavelength of 420 nm for our OD reads.

#### Carbon and Nitrogen Utilization

Carbon source utilization was determined using Biomerieux C32 carbon strips, which are typically used for identification of human pathogens (Tragiannidis et al., 2012). Following the protocols provided by the vendor, we were able to show that *E. viscosa* JF 03-3F and *E. viscosa* JF 03-4F can utilize a wide range of carbon sources. Triplicates were performed on these strips to ensure results were uniform and representative. Overall, a variety of carbon sources supported robust growth of both strain (e.g., D-glucose, L-sorbose, D-galactose, N-acetyl glucosamine, D-sorbitol, D-xylose, glycerol, L-rhamnose, L-arabinose, D-cellobiose, and maltose), and there are only a few quantitative differences in utilization patterns (Figure 9; Tables 7 & 8). Carbon sources that could not support growth of either strain included D-raffinose, D-melibiose, methyl-aD-glucopyranoside, and D-lactose. Genome annotation did not provide an obvious explanation for this observation (Supplemental File 3). Both strains were resistant to cycloheximide and were capable of producing a black color in the presence of esculin ferric citrate (positive for esculin metabolism) (Figure 9). Notably, for both *E. viscosa* JF 03-3F and *E. viscosa* JF 03-4F, growth on some carbon sources, particularly sorbose, levulinic acid, and N-acetyglucosamine, lead to enhanced pigmentation (Figure 9).

**Figure 9:**
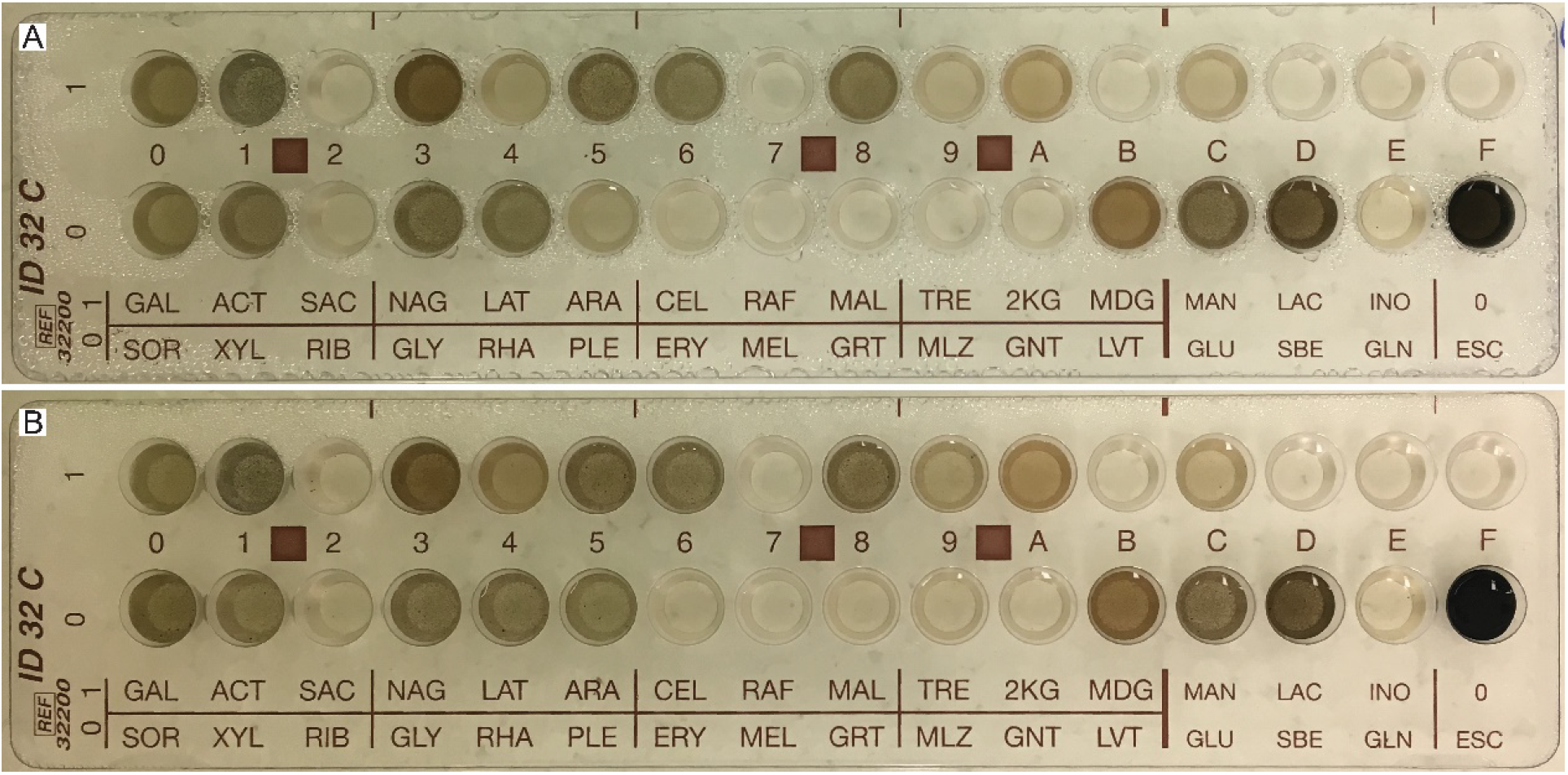
Growth of (A) *E. viscosa* JF 03-3F and (B) *E. viscosa* JF 03-4F on C32 strips for determining carbon utilization. Both species were capable of using the same carbon sources, though some variations are seen. *E. viscosa* JF 03-4F was better at growing on palatinose (PLE), trehalose (TRE), potassium 2- ketogluconate (2KG), and mannitol (MAN) than *E. viscosa* JF 03-3F. *E. viscosa* JF 03-3F was not better at growing on any carbon sources than *E. viscosa* JF 03-4F.

**Table 7:** Carbon source utilization values of E. viscosa JF 03-3F

| *E. viscosa* JF 03-3F |  | # 1 | # 2 | # 3 | Average | Score |
| --- | --- | --- | --- | --- | --- | --- |
| *D-galactose* | GAL | 4 | 5 | 4 | 4.3 | + |
| *D-sorbitol* | SOR | 4 | 4 | 4 | 4.0 | + |
| *actidione (cycloheximide)* | ACT | 5 | 4 | 4 | 4.3 | + |
| *D-xylose* | XYL | 4 | 4 | 4 | 4.0 | + |
| *D-saccharose (sucrose)* | SAC | 1 | 1 | 1 | 1.0 | - |
| *D-ribose* | RIB | 2 | 2 | 2 | 2.0 | V |
| *N-acetyl-glucosamine* | NAG | 4 | 5 | 4 | 4.3 | + |
| *glycerol* | GLY | 4 | 4 | 4 | 4.0 | + |
| *lactic acid* | LAT | 3 | 3 | 3 | 3.0 | V |
| *L-rhamnose* | RHA | 4 | 4 | 4 | 4.0 | + |
| *L-arabinose* | ARA | 4 | 4 | 4 | 4.0 | + |
| *palatinose* | PLE | 3 | 3 | 3 | 3.0 | V |
| *D-cellobiose* | CEL | 4 | 4 | 4 | 4.0 | + |
| *erythritol* | ERY | 2 | 2 | 2 | 2.0 | V |
| *D-raffinose* | RAF | 1 | 1 | 1 | 1.0 | - |
| *D-melibiose* | MEL | 1 | 1 | 1 | 1.0 | - |
| *D-maltose* | MAL | 4 | 4 | 4 | 4.0 | + |
| *sodium glucuronate* | GRT | 2 | 2 | 2 | 2.0 | V |
| *D-trehalose* | TRE | 3 | 3 | 3 | 3.0 | V |
| *D-melezitose* | MLZ | 2 | 2 | 2 | 2.0 | V |
| *potassium 2-ketogluconate* | 2KG | 4 | 3 | 4 | 3.7 | + |
| *potassium gluconate* | GNT | 2 | 2 | 2 | 2.0 | V |
| *methyl-αD-glucopyranoside* | MDG | 1 | 1 | 1 | 1.0 | - |
| *levulinic acid (leuvulinate)* | LVT | 4 | 3 | 4 | 3.7 | + |
| *D-mannitol* | MAN | 3 | 3 | 3 | 3.0 | V |
| *D-glucose* | GLU | 5 | 5 | 5 | 5.0 | + |
| *D-lactose* | LAC | 1 | 1 | 1 | 1.0 | - |
| *L-sorbose* | SBE | 5 | 5 | 5 | 5.0 | + |
| *inositol* | INO | 2 | 2 | 2 | 2.0 | V |
| *glucosamine* | GLN | 1 | 1 | 1 | 1.0 | - |
| *no substrate* | No | 1 | 1 | 1 | 1.0 | - |
| *esculin ferric citrate* | ESC | + | + | + | + | Black + |

**Table 8:** Carbon source utilization values of E. viscosa JF 03-4F

| *E. viscosa* JF 03-4F |  | # 1 | # 2 | # 3 | Average | Score |
| --- | --- | --- | --- | --- | --- | --- |
| *D-galactose* | GAL | 4 | 4 | 4 | 4.0 | + |
| *D-sorbitol* | SOR | 5 | 5 | 5 | 5.0 | + |
| *actidione (cycloheximide)* | ACT | 4 | 4 | 4 | 4.0 | + |
| *D-xylose* | XYL | 4 | 4 | 4 | 4.0 | + |
| *D-saccharose (sucrose)* | SAC | 2 | 2 | 2 | 2.0 | V |
| *D-ribose* | RIB | 2 | 2 | 3 | 2.3 | V |
| *N-acetyl-glucosamine* | NAG | 4 | 4 | 4 | 4.0 | + |
| *glycerol* | GLY | 4 | 4 | 4 | 4.0 | + |
| *lactic acid* | LAT | 3 | 3 | 3 | 3.0 | V |
| *L-rhamnose* | RHA | 4 | 4 | 4 | 4.0 | + |
| *L-arabinose* | ARA | 4 | 4 | 4 | 4.0 | + |
| *palatinose* | PLE | 4 | 4 | 4 | 4.0 | + |
| *D-cellobiose* | CEL | 4 | 4 | 4 | 4.0 | + |
| *erythritol* | ERY | 1 | 2 | 2 | 1.7 | V |
| *D-raffinose* | RAF | 1 | 1 | 1 | 1.0 | - |
| *D-melibiose* | MEL | 1 | 1 | 1 | 1.0 | - |
| *D-maltose* | MAL | 4 | 4 | 4 | 4.0 | + |
| *sodium glucuronate* | GRT | 2 | 2 | 2 | 2.0 | V |
| *D-trehalose* | TRE | 4 | 3 | 4 | 3.7 | + |
| *D-melezitose* | MLZ | 2 | 2 | 2 | 2.0 | V |
| *potassium 2-ketogluconate* | 2KG | 3 | 3 | 4 | 3.3 | V |
| *potassium gluconate* | GNT | 2 | 2 | 2 | 2.0 | V |
| *methyl-αD-glucopyranoside* | MDG | 1 | 1 | 1 | 1.0 | - |
| *levulinic acid (leuvulinate)* | LVT | 3 | 4 | 3 | 3.3 | V |
| *D-mannitol* | MAN | 3 | 3 | 4 | 3.3 | V |
| *D-glucose* | GLU | 5 | 5 | 5 | 5.0 | + |
| *D-lactose* | LAC | 1 | 1 | 1 | 1.0 | - |
| *L-sorbose* | SBE | 5 | 5 | 5 | 5.0 | + |
| *inositol* | INO | 2 | 2 | 2 | 2.0 | V |
| *glucosamine* | GLN | 2 | 2 | 1 | 1.7 | V |
| *no substrate* | 0 | 1 | 1 | 1 | 1.0 | - |
| *esulin ferric citrate* | ESC | + | + | + | + | Black + |

We were particularly interested in patterns of nitrogen utilization for *E. viscosa* JF 03-3F and *E. viscosa* JF 03-4F given their isolation from a nutrient depleted niche with extensive nitrogen-fixing bacterial populations. Nine different nitrogen sources were tested: five amino acids (aspartate, glutamate, glycine, proline, serine), ammonium tartrate, sodium nitrate, urea, peptone (mixed short chain amino acids) as a positive control, and no nitrogen as a negative control. Both strains are incapable of utilizing aspartate and glutamate as nitrogen sources; *E. viscosa* JF 03-3F having a p-value of 0.012 (n=6) for aspartate and 0.0005 (n=6) for glutamate; *E. viscosa* JF 03-4F having a p-value of 0.0003 (n=6) for aspartate and .00005 (n=6) for Glutamate (all compared to peptone) (Figure 10). E. viscosa JF 03-3F and E. viscosa JF 03-4F differed in their preferred nitrogen sources. Both preferred proline, p-value for *E. viscosa* JF 03-3F 0.024 (n=6), p-value for E. viscosa JF 03-4F is 0.0012 (n=6), over peptone. However, they differ in that *E. viscosa* JF 03-3F most prefers urea p-value 0.009 (n=6), while *E. viscosa* JF 03-4F had additional significantly preferred nitrogen sources including serine (p-value 0.0064 (n=6)), ammonium (p-value 0.0007 (n=6)), and nitrate (p-value 0.022 (n=6) (Figure 10).

**Figure 10:**
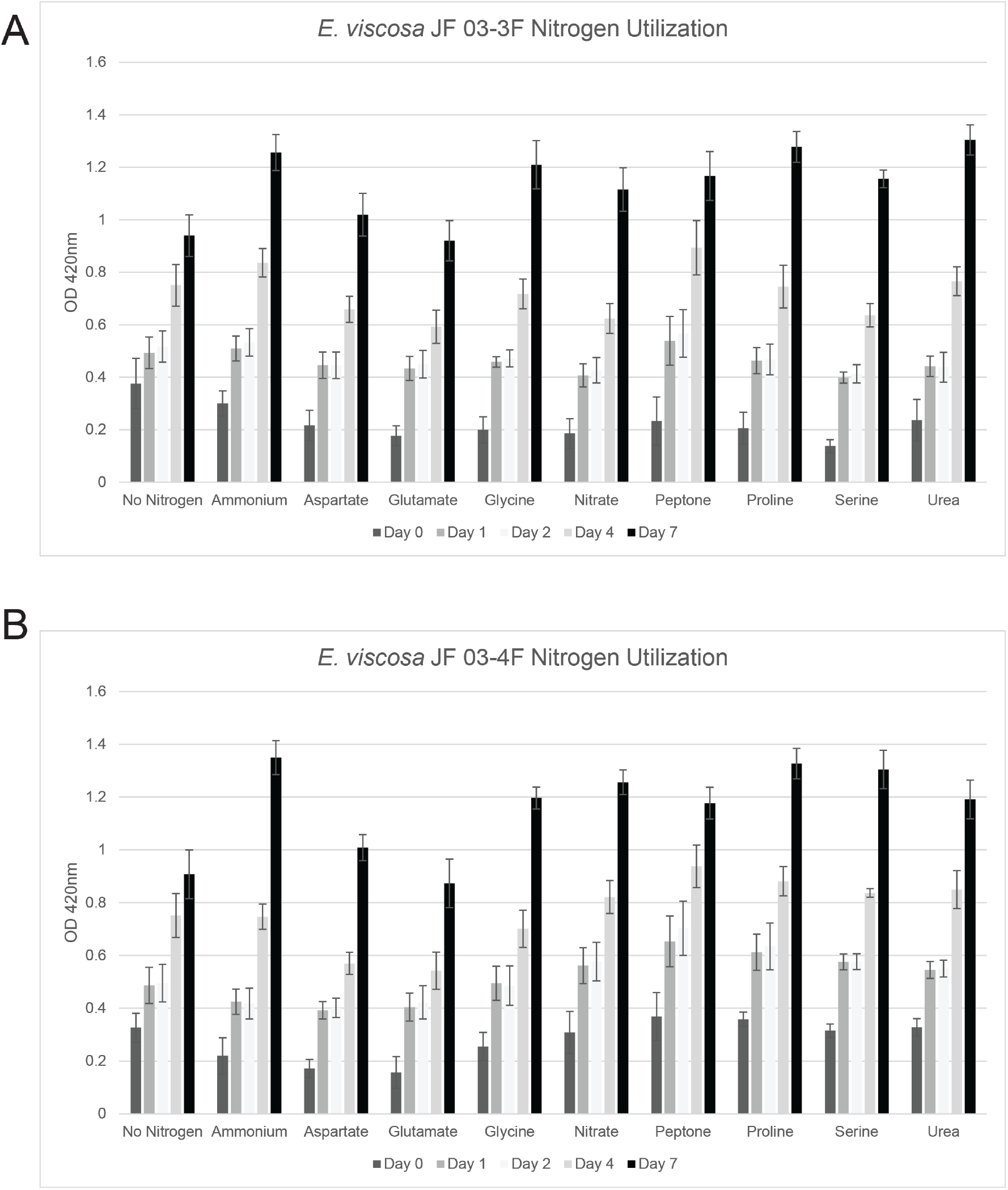
Nitrogen source utilization of (A) *E. viscosa* JF 03-3F and (B) *E. viscosa* JF 03-4F in liquid culture. Neither species was capable of using aspartate or glutamate as a nitrogen source, as their growth amount were equivalent to no nitrogen. All other nitrogen sources tested were used by both species with varying preference. Ammonium was the preferred nitrogen source for *E. viscosa* JF 03-4F (p-value = 0.0007 compared to peptone (n=6)), and *E. viscosa* JF 03-3F preferred urea for a nitrogen source (p-value = 0.009 compared to peptone (n=6)). *E. viscosa* JF 03-3F was unable to use aspartate or glutamate as nitrogen sources, but *E. viscosa* JF 03-4F was able to use aspartate as a nitrogen source (p-value = 0.03 compared to no nitrogen (n=6)). * = p-value of 0.05 > (n=6) compared to peptone positive control, ** = p-value of 0.01 > (n=6) compared to peptone positive control, + = p-value of 0.05 > (n=6) compared to no nitrogen negative control.

#### Optimal Growth Temperature and Range of Growth Temperatures

*E. viscosa* JF 03-3F and *E. viscosa* JF 03-4F were isolated from an environment that experiences wide seasonal and diurnal temperature changes. As such, we determined the optimal growing temperature for these strains, as well as the limits at which they are capable of survival (cardinal temperatures). Both isolates were serial diluted and spotted onto MEA plates to determine the carrying capacity at different temperatures. Both *E. viscosa* JF 03-3F *and E. viscosa* JF 03-4F were capable of growth at 4 °C, 15 °C, 23 °C, and 27 °C, but could not grow at 37 °C and 42 °C (Figure 11). Optimal growth temperature of both *E. viscosa* JF 03-3F and *E. viscosa* JF 03-4F was 23 °C (Figure 11). Growth at 4 °C was slow, but after a year the isolates both strains formed extensive colonies and flooded the agar with melanin (Figure S6). Although neither strain grew at 37 °C, they retained viability as they were able to resume growth following return to 23 °C after two days of exposure to 37 °C (Figure S6). In contrast, the same experiment showed that incubation at 42 °C is lethal.

**Figure 11:**
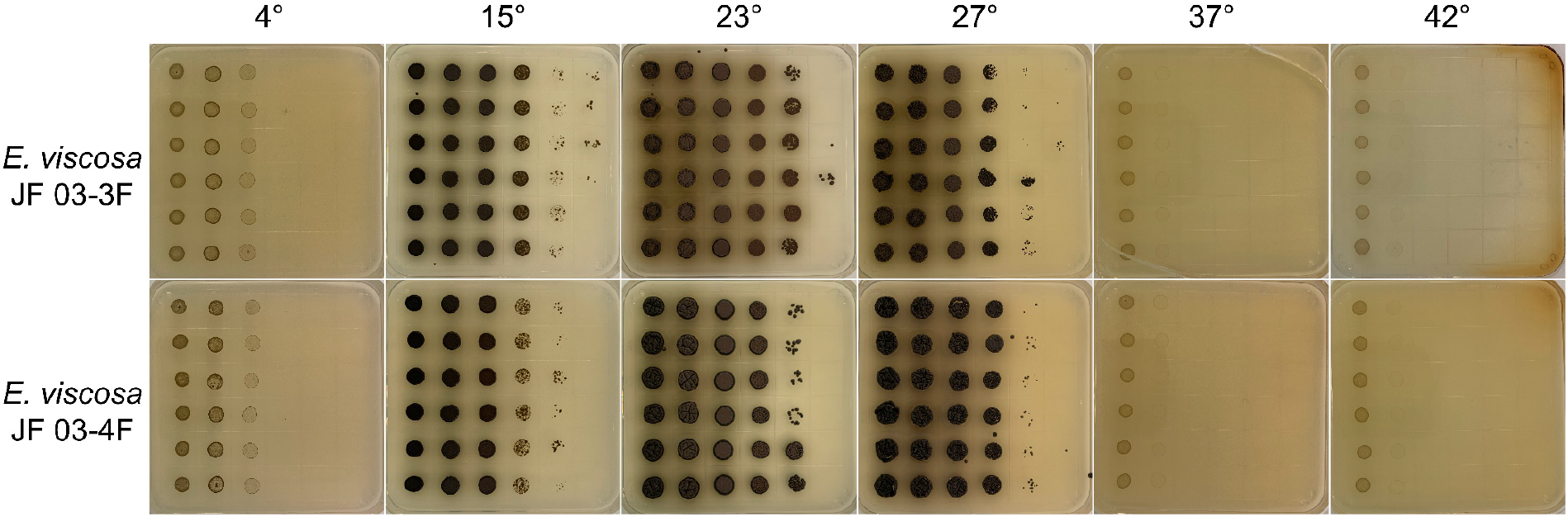
Growth of *E. viscosa* JF 03-3F and *E. viscosa* JF 03-4F at varying temperatures. Neither species was capable of growth at or above 37 °C, implying they are not human pathogens. Both species optimally grew at 23 °C, but were capable of growth down to 4 °C.

#### UV and Metal Resistance

Melanized fungi are recognized for their resistance to UV light, and the possibility of using ionizing radiation as an energy source (Dadachova et al., 2007). To assess the response of *E. viscosa* JF 03-3F and *E. viscosa* JF 03-4F to UV radiation, they were each exposed to a dose (120 seconds of 10,000 μJ/cm^2^ at 254 nm) that was lethal to *S. cerevisiae* and *E. dermatitidis* (data not shown). The same level of exposure did not kill either *E. viscosa* JF 03-3F or *E. viscosa* JF 03-4F (Figure 12) but did significantly reduce the number of viable colonies. Strikingly, surviving colonies showed no evidence of induced mutagenesis based on the absence of altered morphologies or pigmentation (Figure 12).

**Figure 12:**
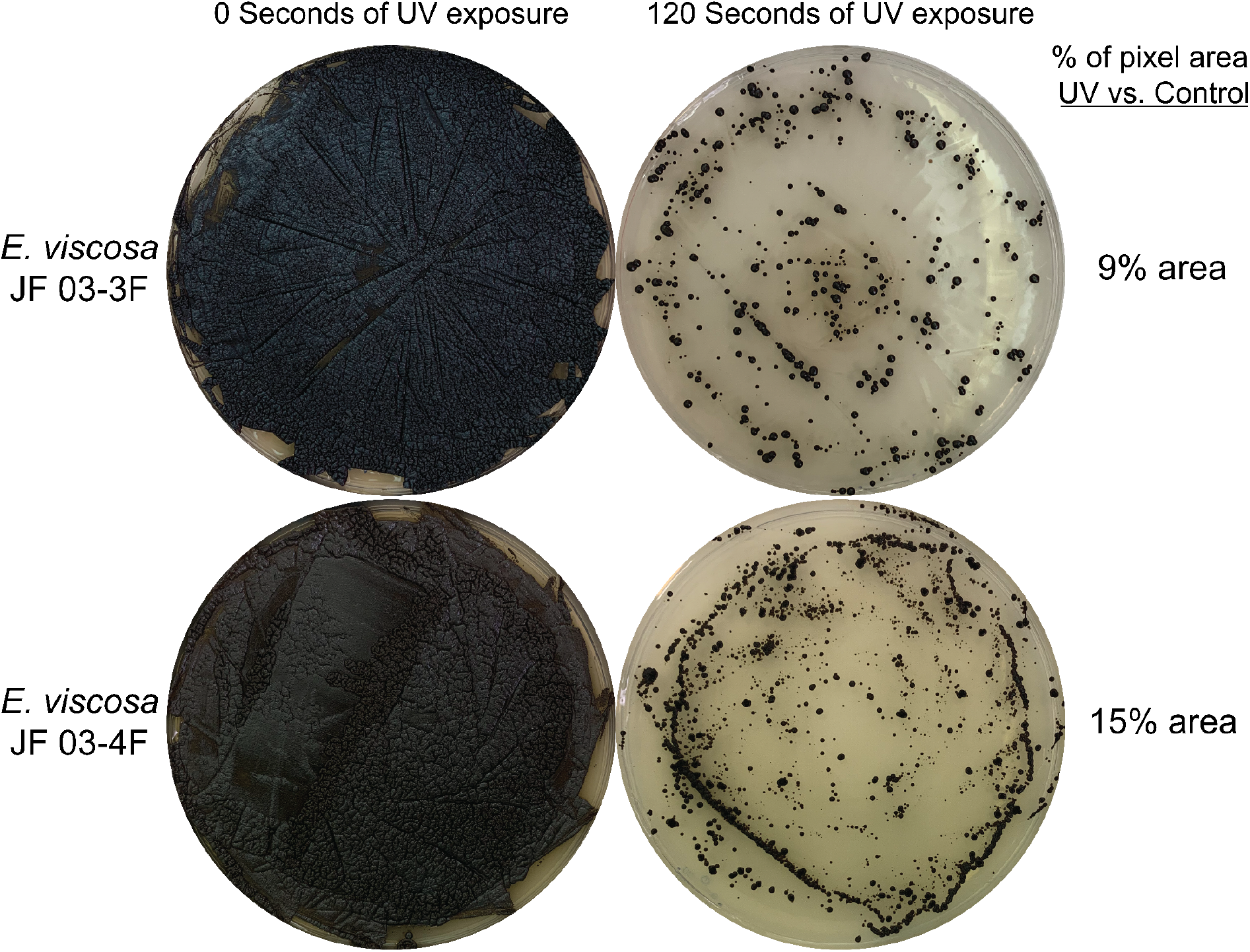
Difference in growth of *E. viscosa* JF 03-3F and *E. viscosa* JF 03-4F with and without exposure to UV light. *E. viscosa* JF 03-3F showed slightly less resistance to the UV exposure than *E. viscosa* JF 03- 4F. Neither species was mutated from 120 seconds of UV exposure, normally *S. cerevisiae* and *E. dermatitidis* are incapable of growth after the same amount of UV exposure (data not shown).

Polyextremotolerant fungi have been previously noted as having increased metal resistances as a result of their melanized cell wall and other adaptations to harsh environments (Gadd & de Rome, 1988). To test if these two new *Exophiala* spp. possess increased metal resistances when compared to *E. dermatitidis* and *S. cerevisiae*, we impregnated Whatman filter discs with various metals of different concentrations. Diameters of zones of clearing revealed no evidence for enhanced metal resistance to the metals tested (Table 9). On the other hand, both strains appear to be moderately more sensitive to NiCl_2_ and CdCl_2_ (Table 9).

**Table 9:** Diameter of the zone of clearing of E. viscosa JF 03-3F, E. viscosa JF 03-4F, E. dermatitidis, and S. cerevisiae with various metals.

| Species | FeSO<sub>4</sub> 1 M | CoCl<sub>2</sub> 0.5 M | AgNO<sub>3</sub> 0.47 M | NiCl<sub>2</sub> 1.5 M | CuCl<sub>2</sub> 1.5 M | CdCl<sub>2</sub> 10 mM |
| --- | --- | --- | --- | --- | --- | --- |
| *E. viscosa*<br>JF 03-3F | 1.6 cm | 1.5 cm | 1.4 cm | 3.9 cm | 1.3 cm | 4.5 cm |
| *E. viscosa*<br>JF 03-4F | 1.8 cm | 1.8 cm | 1.4 cm | 4 cm | 1.5 cm | 3.9 cm |
| *E. dermatitidis* | 1.3 cm | 1 cm | 1.5 cm | 2 cm | 2.5 cm | 1.2 cm |
| *S. cerevisiae* | 1 cm | 1.9 cm | 2 cm | 2.1 cm | 2 cm | 2.5 cm |

#### Lipid Profiles

Both *E. viscosa* JF 03-3F and *E. viscosa* JF 03-4F appear to possess abundant lipid bodies (Figure S4). This observation along with the unique sticky morphology of both strains led us to the idea that they might contain unique or copious amounts of lipids. We performed a Bligh and Dyer lipid extraction followed by thin layer chromatography (TLC) to observe patterns of lipid accumulation in *E. viscosa* JF 03-3F and *E. viscosa* JF 03-4F grown on media that contained fermentable or non-fermentable sugars, both with high nitrogen and low nitrogen. These iterations creating four unique media types would allow us to cover as much of a spread of lipid changes as possible. *S. cerevisiae* and *E. dermatitidis* were also similarly analyzed. Results from this lipid extraction showed that our two new strains did not seem to produce any unique or novel amounts or types of lipids when compared to *S. cerevisiae* and *E. dermatitidis* (Figure S7).

### Melanin Production and Regulation in *E. viscosa* JF 03-3F and *E. viscosa* JF 03-4F

#### Melanin Biosynthesis Gene Annotation

A defining feature of black yeasts is their pigmentation caused by the accumulation of melanin (Bell & Wheeler, 1986). Given the presumed importance of melanin to a polyextremotolerant lifestyle, we are interested in understanding how melanin production is regulated. A first step in this process is to determine the types of melanin that *E. viscosa* JF 03-3F and *E. viscosa* JF 03-4F are capable of producing. To accomplish this, the sequenced and automatically annotated genomes of *E. viscosa* JF 03-3F and *E. viscosa* JF 03-4F were manually annotated using protein sequences for all three melanin biosynthetic pathways in *Aspergillus niger* (Teixeira et al., 2017). The list of *A. niger* (and two *A. fumigatus*) proteins and their homologs in both *E. viscosa* JF 03-3F and *E. viscosa* JF 03-4F are summarized in Table 10. Manual annotation and reverse BLASTP of the melanin biosynthesis pathway genes showed that both *E. viscosa* JF 03-3F and *E. viscosa* JF 03-4F have the genetic capability to produce all three forms of known fungal melanins: DOPA-melanin (pheomelanin), DHN-melanin (allomelanin), and L-tyrosine/HGA derived pyomelanin. Two *A. fumigatus* proteins, hydroxynaphthalene reductase (Arp2) and scytalone dehydratase (Arp1) were also separately BLASTPed against the sequences of *E. viscosa* JF 03-3F and *E. viscosa* JF 03-4F because those protein sequences are not homologous between *A. niger* and *A. fumigatus* (Jørgensen et al., 2011). Both *E. viscosa* JF 03-3F and *E. viscosa* JF 03-4F were found to have homologous proteins to both *A. fumigatus* and *A. niger*’s Arp1 and Arp2 (Table 10).

**Table 10:**
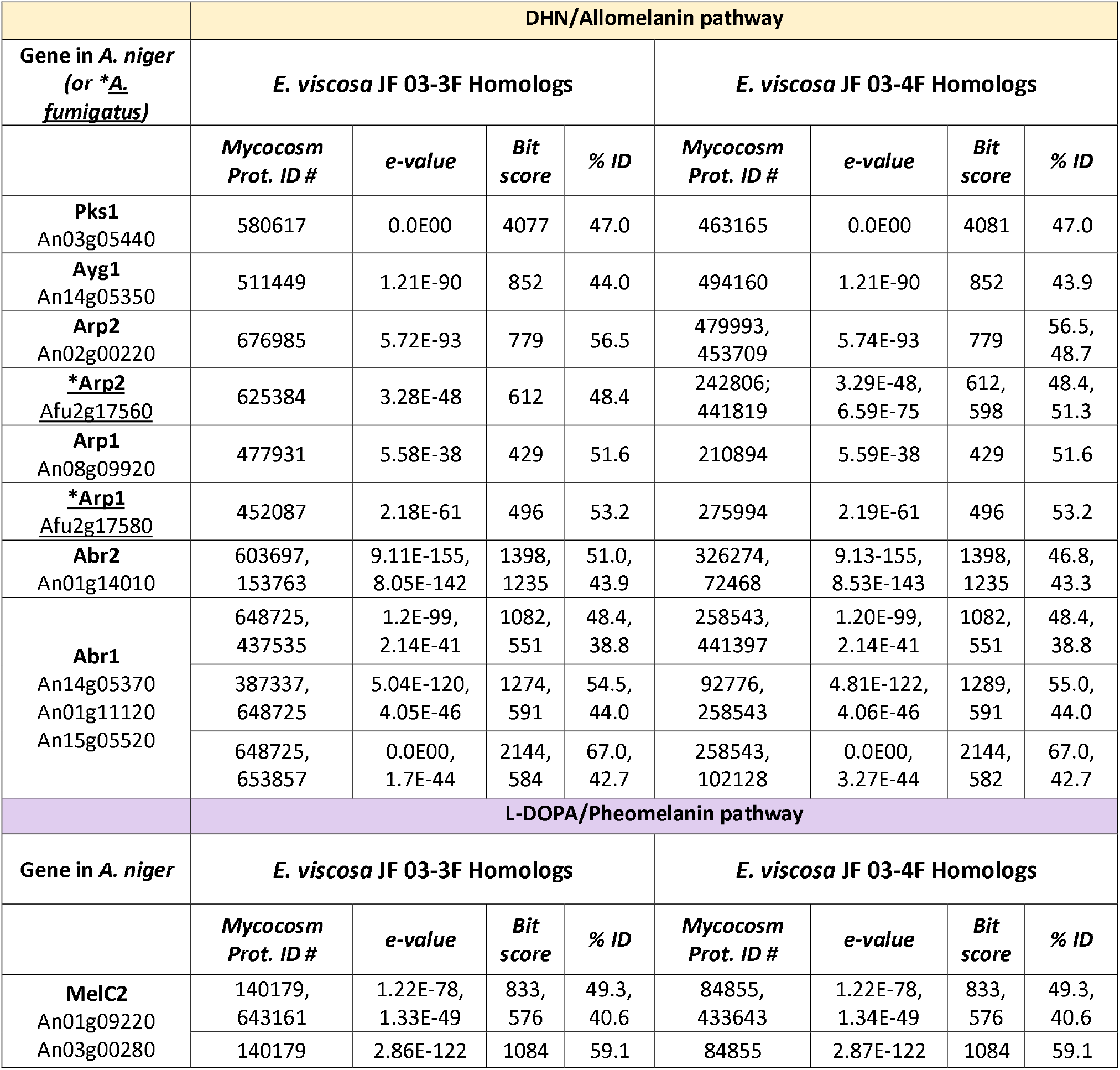
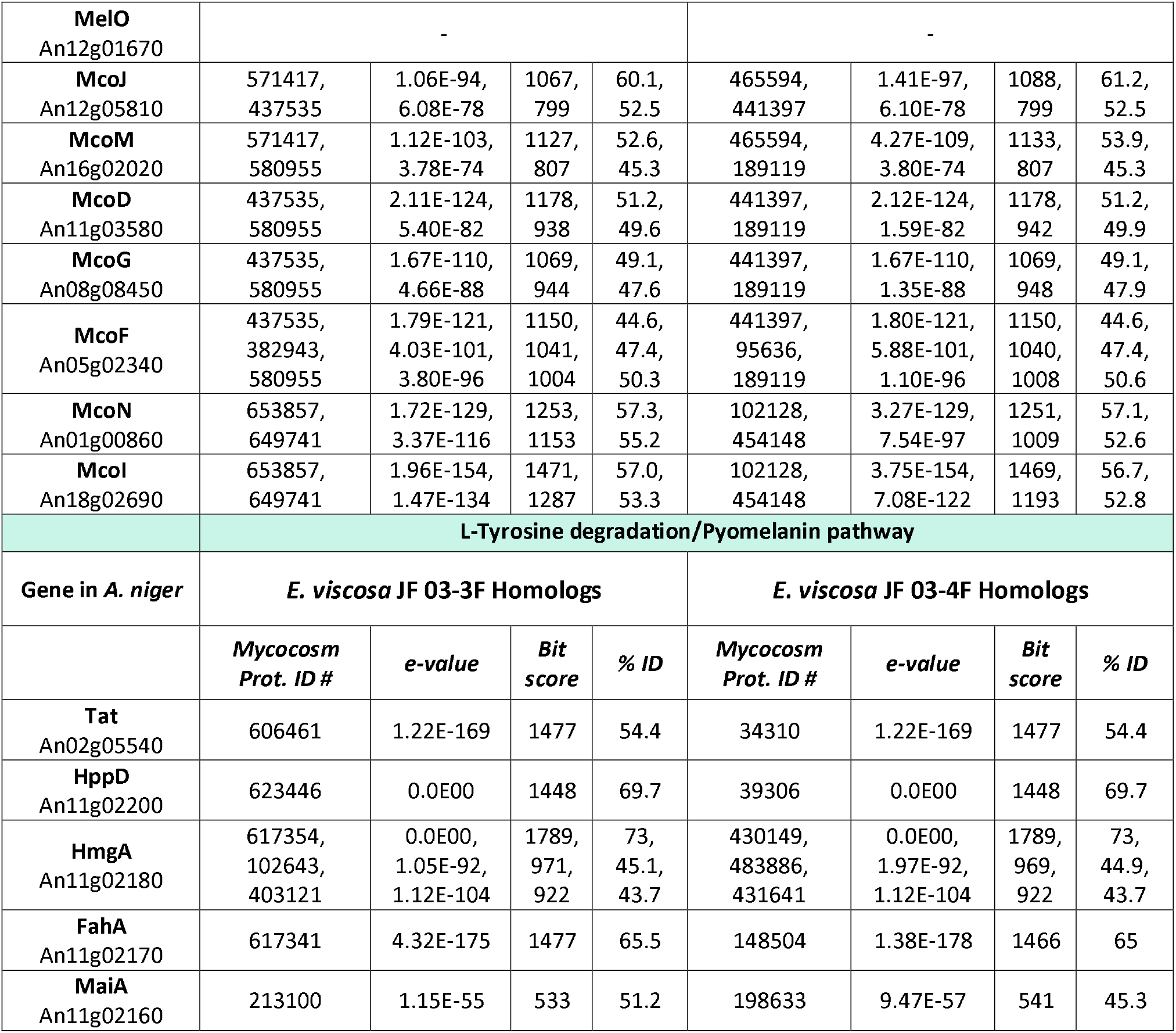
Annotation of melanin biosynthetic genes for E. viscosa JF 03-3F and JF 03-4F

|  | DHN/Allomelanin pathway |  |  |  |  |  |  |  |
| --- | --- | --- | --- | --- | --- | --- | --- | --- |
| Gene in *A. niger*<br>(or *\*A. fumigatus*) | *E. viscosa* JF 03-3F Homologs |  |  |  | *E. viscosa* JF 03-4F Homologs |  |  |  |
|  | *Mycocosm*<br>Prot. ID # | *e-value* | *Bit*<br>score | % ID | *Mycocosm*<br>Prot. ID # | *e-value* | *Bit*<br>score | % ID |
| **Pks1**<br>An03g05440 | 580617 | 0.0E00 | 4077 | 47.0 | 463165 | 0.0E00 | 4081 | 47.0 |
| **Ayg1**<br>An14g05350 | 511449 | 1.21E-90 | 852 | 44.0 | 494160 | 1.21E-90 | 852 | 43.9 |
| **Arp2**<br>An02g00220 | 676985 | 5.72E-93 | 779 | 56.5 | 479993,<br>453709 | 5.74E-93 | 779 | 56.5,<br>48.7 |
| **\*Arp2**<br><u>Afu2g17560</u> | 625384 | 3.28E-48 | 612 | 48.4 | 242806;<br>441819 | 3.29E-48,<br>6.59E-75 | 612,<br>598 | 48.4,<br>51.3 |
| **Arp1**<br>An08g09920 | 477931 | 5.58E-38 | 429 | 51.6 | 210894 | 5.59E-38 | 429 | 51.6 |
| **\*Arp1**<br><u>Afu2g17580</u> | 452087 | 2.18E-61 | 496 | 53.2 | 275994 | 2.19E-61 | 496 | 53.2 |
| **Abr2**<br>An01g14010 | 603697,<br>153763 | 9.11E-155,<br>8.05E-142 | 1398,<br>1235 | 51.0,<br>43.9 | 326274,<br>72468 | 9.13-155,<br>8.53E-143 | 1398,<br>1235 | 46.8,<br>43.3 |
| **Abr1**<br>An14g05370<br>An01g11120<br>An15g05520 | 648725,<br>437535 | 1.2E-99,<br>2.14E-41 | 1082,<br>551 | 48.4,<br>38.8 | 258543,<br>441397 | 1.20E-99,<br>2.14E-41 | 1082,<br>551 | 48.4,<br>38.8 |
|  | 387337,<br>648725 | 5.04E-120,<br>4.05E-46 | 1274,<br>591 | 54.5,<br>44.0 | 92776,<br>258543 | 4.81E-122,<br>4.06E-46 | 1289,<br>591 | 55.0,<br>44.0 |
|  | 648725,<br>653857 | 0.0E00,<br>1.7E-44 | 2144,<br>584 | 67.0,<br>42.7 | 258543,<br>102128 | 0.0E00,<br>3.27E-44 | 2144,<br>582 | 67.0,<br>42.7 |
|  | L-DOPA/Pheomelanin pathway |  |  |  |  |  |  |  |
| Gene in *A. niger* | *E. viscosa* JF 03-3F Homologs |  |  |  | *E. viscosa* JF 03-4F Homologs |  |  |  |
|  | *Mycocosm*<br>Prot. ID # | *e-value* | *Bit*<br>score | % ID | *Mycocosm*<br>Prot. ID # | *e-value* | *Bit*<br>score | % ID |
| **MelC2**<br>An01g09220<br>An03g00280 | 140179,<br>643161 | 1.22E-78,<br>1.33E-49 | 833,<br>576 | 49.3,<br>40.6 | 84855,<br>433643 | 1.22E-78,<br>1.34E-49 | 833,<br>576 | 49.3,<br>40.6 |
|  | 140179 | 2.86E-122 | 1084 | 59.1 | 84855 | 2.87E-122 | 1084 | 59.1 |
| **MeIO**<br>An12g01670 | - |  |  |  | - |  |  |  |
| **McoJ**<br>An12g05810 | 571417,<br>437535 | 1.06E-94,<br>6.08E-78 | 1067,<br>799 | 60.1,<br>52.5 | 465594,<br>441397 | 1.41E-97,<br>6.10E-78 | 1088,<br>799 | 61.2,<br>52.5 |
| **McoM**<br>An16g02020 | 571417,<br>580955 | 1.12E-103,<br>3.78E-74 | 1127,<br>807 | 52.6,<br>45.3 | 465594,<br>189119 | 4.27E-109,<br>3.80E-74 | 1133,<br>807 | 53.9,<br>45.3 |
| **McoD**<br>An11g03580 | 437535,<br>580955 | 2.11E-124,<br>5.40E-82 | 1178,<br>938 | 51.2,<br>49.6 | 441397,<br>189119 | 2.12E-124,<br>1.59E-82 | 1178,<br>942 | 51.2,<br>49.9 |
| **McoG**<br>An08g08450 | 437535,<br>580955 | 1.67E-110,<br>4.66E-88 | 1069,<br>944 | 49.1,<br>47.6 | 441397,<br>189119 | 1.67E-110,<br>1.35E-88 | 1069,<br>948 | 49.1,<br>47.9 |
| **McoF**<br>An05g02340 | 437535,<br>382943,<br>580955 | 1.79E-121,<br>4.03E-101,<br>3.80E-96 | 1150,<br>1041,<br>1004 | 44.6,<br>47.4,<br>50.3 | 441397,<br>95636,<br>189119 | 1.80E-121,<br>5.88E-101,<br>1.10E-96 | 1150,<br>1040,<br>1008 | 44.6,<br>47.4,<br>50.6 |
| **McoN**<br>An01g00860 | 653857,<br>649741 | 1.72E-129,<br>3.37E-116 | 1253,<br>1153 | 57.3,<br>55.2 | 102128,<br>454148 | 3.27E-129,<br>7.54E-97 | 1251,<br>1009 | 57.1,<br>52.6 |
| **McoI**<br>An18g02690 | 653857,<br>649741 | 1.96E-154,<br>1.47E-134 | 1471,<br>1287 | 57.0,<br>53.3 | 102128,<br>454148 | 3.75E-154,<br>7.08E-122 | 1469,<br>1193 | 56.7,<br>52.8 |
| **L-Tyrosine degradation/Pyomelanin pathway** |  |  |  |  |  |  |  |  |
| **Gene in *A. niger*** | ***E. viscosa* JF 03-3F Homologs** |  |  |  | ***E. viscosa* JF 03-4F Homologs** |  |  |  |
|  | ***Mycocosm***<br>**Prot. ID #** | ***e-value*** | ***Bit***<br>***score*** | **% ID** | ***Mycocosm***<br>**Prot. ID #** | ***e-value*** | ***Bit***<br>***score*** | **% ID** |
| **Tat**<br>An02g05540 | 606461 | 1.22E-169 | 1477 | 54.4 | 34310 | 1.22E-169 | 1477 | 54.4 |
| **HppD**<br>An11g02200 | 623446 | 0.0E00 | 1448 | 69.7 | 39306 | 0.0E00 | 1448 | 69.7 |
| **HmgA**<br>An11g02180 | 617354,<br>102643,<br>403121 | 0.0E00,<br>1.05E-92,<br>1.12E-104 | 1789,<br>971,<br>922 | 73,<br>45.1,<br>43.7 | 430149,<br>483886,<br>431641 | 0.0E00,<br>1.97E-92,<br>1.12E-104 | 1789,<br>969,<br>922 | 73,<br>44.9,<br>43.7 |
| **FahA**<br>An11g02170 | 617341 | 4.32E-175 | 1477 | 65.5 | 148504 | 1.38E-178 | 1466 | 65 |
| **MaiA**<br>An11g02160 | 213100 | 1.15E-55 | 533 | 51.2 | 198633 | 9.47E-57 | 541 | 45.3 |

#### Regulation of Melanin Production Using Chemical Blockers

Because *E. viscosa* JF 03-3F and *E. viscosa* JF 03-4F possess the genetic capability to produce all three forms of fungal melanin, we asked whether they were all being produced simultaneously or if one was preferred for production versus excretion. We used chemical blockers for both L-DOPA-melanin and DHN-melanin to determine the predominant type of melanin produced on MEA and YPD. Kojic acid blocks the production of L-DOPA melanin, whereas phthalide inhibits the synthesis of DHN-melanin; both are effective at doses of 1, 10, and 100 μg/mL in various fungi (Pal et al., 2014).

First, we used 100 mM of phthalide and 100 mg/mL of kojic acid in a filter disc assay. Placing drug-impregnated filter discs on freshly spread lawns of cells either individually or with both drugs combined did not block melanin production even though the concentrations of the drugs were higher than that of previous studies (Figure S8). Then using Pal et al.’s (2014) method of melanin blocking, we added the highest dosage (100 µg/mL) of phthalide and kojic acid individually and combined to agar-solidified MEA, and both spread a lawn of cells and spotted serially diluted cells onto the plates. Neither assay resulted in blockage of melanin production in *E. viscosa* JF 03-3F or *E. viscosa* JF 03-4F (Figures 13 & S8). However, addition of 100 µg/mL of phthalide alone did result in their apparent excretion of a dark substance into MEA (Figure 13). Overall, chemical inhibition of L-DOPA-melanin or DHN-melanin production did not qualitatively reduce colony pigmentation, possibly suggesting that the tyrosine-derived pyomelanin is still being produced.

**Figure 13:**
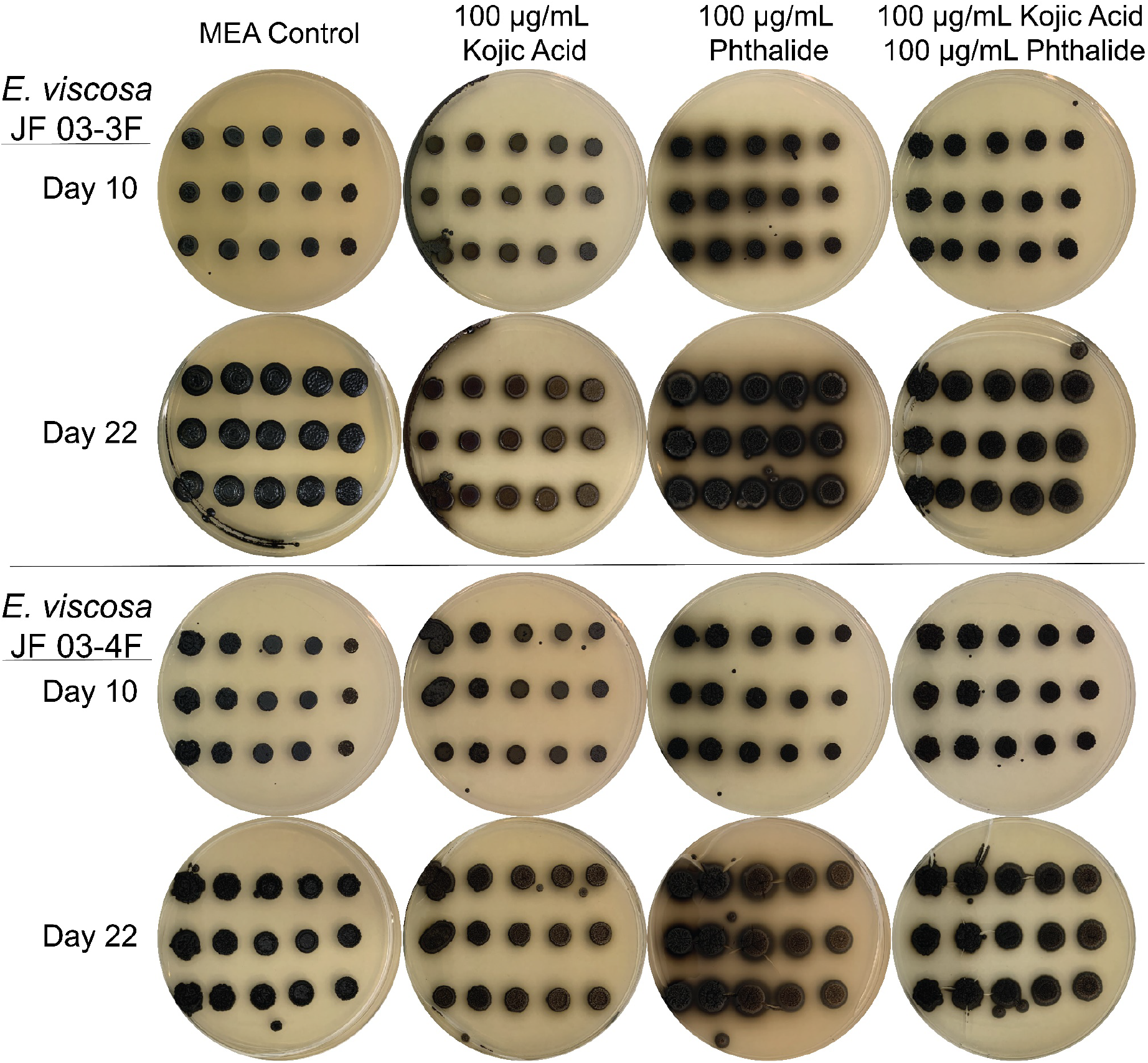
Growth of serial diluted (0x-10,000x with 10x steps) *E. viscosa* JF 03-3F and *E. viscosa* JF 03-4F in the presence of melanin blockers kojic acid and phthalide. Neither chemical melanin blocker was capable of blocking melanin production in either fungus, even when both chemical blockers were used simultaneously. Additionally, use of phthalide on *E. viscosa* JF 03-3F induced melanin secretion on a medium where this does not usually occur. The melanin halo around *E. viscosa* JF 03-3F’s colonies on medium containing phthalide was replaced with hyphal growth after 22 days.

#### Evaluating the Cause of Melanin Excretion

The appearance of a dark pigment surrounding colonies of *E. viscosa* JF 03-3F and *E. viscosa* JF 03-4F under specific conditions raised the idea that these fungi are able to excrete melanin into their local environment. The presence of dark pigments in the supernatants of liquid cultures lent further support to this.

Initial studies were performed to determine which media triggered the release of melanin. Both *E. viscosa* JF 03-3F and *E. viscosa* JF 03-4F were capable of releasing the most melanin and in the shortest growth time on YPD, with *E. viscosa* JF 03-3F seemingly excreting more than *E. viscosa* JF 03-4F (Figures 7 & 8). Because YPD and MEA only differ by the presence of yeast vs. malt extract as well as the percentage of peptone (2% in YPD, 0.2% in MEA), we first determined if the peptone differences impacted melanin excretion. By switching the peptone amounts in YPD and MEA, we demonstrated that *E. viscosa* JF 03-3F acquired the ability to excrete melanin on MEA if it was supplemented with 2% peptone (Figure S9). To extend this observation, we added progressively higher amounts of peptone to MEA media (i.e., 10 different concentrations of peptone ranging from 0.1% to 5%). We observed that *E. viscosa* JF 03-3F starts excreting melanin on MEA at a peptone amount of 2%, and *E. viscosa* JF 03-4F starts excreting melanin at about 4% peptone (Figure 14).

**Figure 14:**
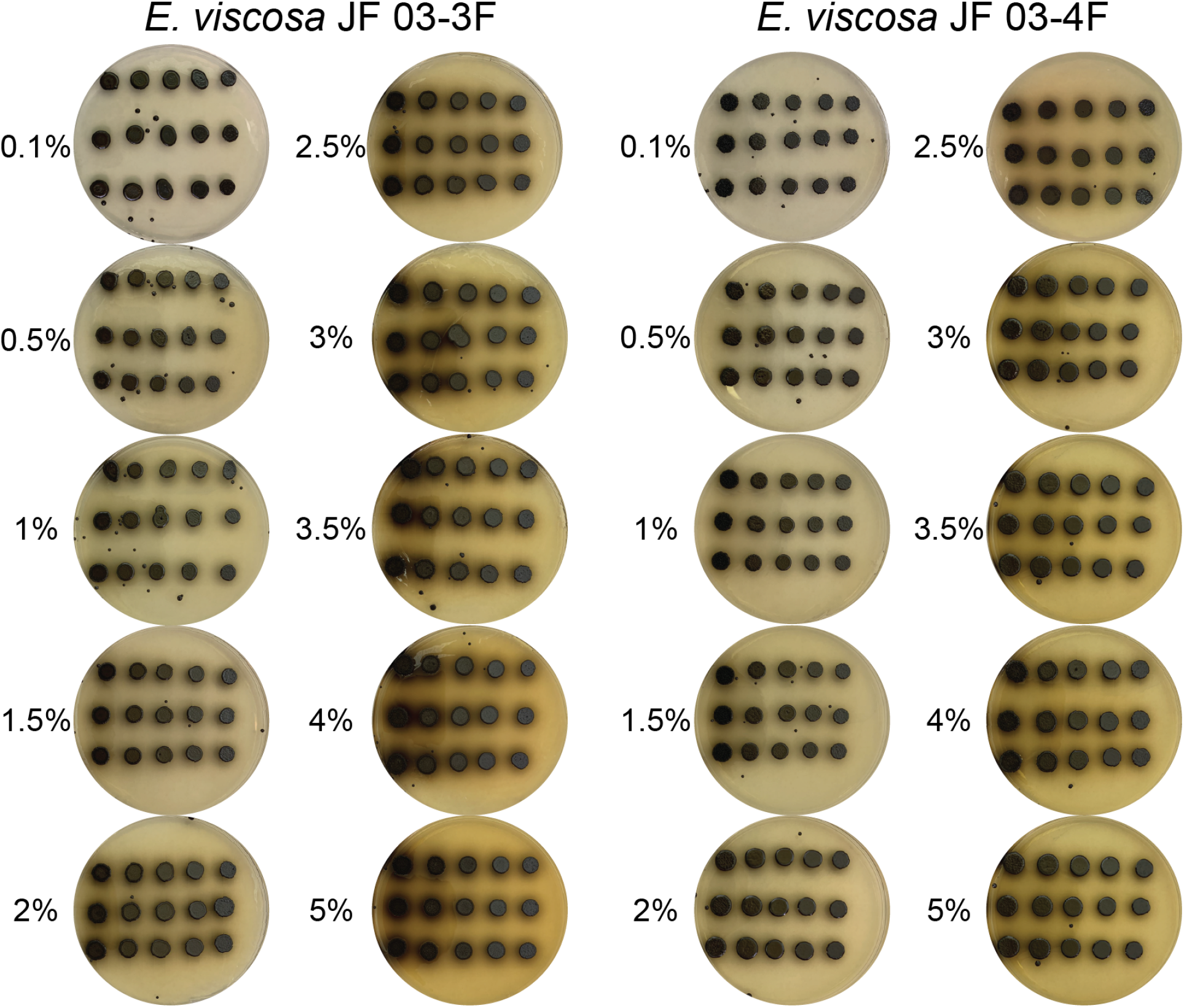
*E. viscosa* JF 03-3F and *E. viscosa* JF 03-4F grown on MEA with increasing amounts of peptone. The higher the amount of peptone in the medium, the more melanin was secreted. *E. viscosa* JF 03-3F started secreting melanin at 2%, and *E. viscosa* JF 03-4F at 4%.

To determine if it was tyrosine (the precursor to L-DOPA and HGA/tyrosine-derived melanins) in the peptone that was inducing melanin excretion at higher peptone levels, we tested differing tyrosine amounts on our fungi. When 0.5 g/L of tyrosine is added to the media, *E. viscosa* JF 03-3F and *E. viscosa* JF 03-4F are capable of melanin excretion, whereas increasing the tyrosine amount to 1 g/L melanin excretion is stopped completely in *E. viscosa* JF 03-4F and is greatly reduced in *E. viscosa* JF 03-3F. Additionally, when tyrosine in the only source of nitrogen at 0.5 g/L, no melanin excretion is observed in the wild type strains (Figure S10). Additional tests of added amino acids and ammonium chloride’s effect on melanin excretion were also performed. A summarized description of the media variants and their results on these fungi’s ability to excrete melanin can be found in Table S3. Additional details and further understanding of these results are to be described at a later date.

#### Confirmation of Melanin Excretion from Living Cells and Determining its OD Amount in the Supernatant

To confirm that the dark pigment in culture supernatants is indeed melanin, we performed a melanin extraction using previously described methods (Pal et al., 2014; Pralea et al., 2019). Although both enzymatic and chemical extraction of melanin was attempted, the chemical extraction process proved to be more efficient at recovering excreted melanin from the supernatant (Figure S11). Therefore, all extractions going forward were performed using the chemical extraction method to ensure all melanin was extracted from the solution.

Growth of *E. viscosa* JF 03-3F*, E. viscosa* JF 03-4F, and *E. dermatitidis* on YPD and MEA were observed daily for melanin excretion; 11 mL aliquots of their supernatant were removed daily, and the melanin was extracted from each sample. As the experiment progressed, it was obvious that *E. viscosa* JF 03-3F and *E. viscosa* JF 03-4F began excreting melanin on day 3 and excreted more in YPD than in MEA (Figure 15). *E. dermatitidis* on the other hand seemed to have a slower build-up of melanin, and melanin wasn’t truly obvious in the medium until day 6 in MEA though it can be seen in MEA day 3 in Figure 15. Analysis of the optical density (OD) of the final extracted melanin revealed that melanin amount increased over the course of the experiment, and that greater amounts of melanin are released in YPD for *E. viscosa* JF 03-3F and *E. viscosa* JF 03-4F compared to MEA for *E. dermatitidis* (Figures S12, S13, & S14). Interestingly, for both *E. viscosa* JF 03-3F and *E. viscosa* JF 03-4F, their peak melanin excretion was on day 6 and not on day 7 when grown in YPD (Figure 16). There was less melanin on day 7 for both strains, indicating that the melanin after day 6 was either degrading, or was being taken back up by the cells. This was only obvious with reading the OD results of the extracted melanin, as the supernatants themselves past day 5 all strikingly dark and hard to differentiate visually (Figure 15). Extracted melanin from all *E. viscosa* JF 03-3F, *E. viscosa* JF 03-4F, and *E. dermatitidis* samples (except *E. dermatitidis* in MEA on day 3) displayed the typical results from a full spectrum read on pure melanin (Figures S12, S13, & S14; Table S2). When graphed, absorbance to Wavelength, the sample should have an exponentially decreasing OD as wavelength increases, and when the log10 of the OD absorbance reads is calculated, the full spectrum reads should become be a linear regression with an R^2^ value of 0.97 or higher (Pralea et al., 2019). All *E. viscosa* JF 03-3F*, E. viscosa* JF 03-4F, and *E. dermatitidis* supernatant samples displayed these features (Table S2), and therefore we can confirm that the dark nature of their supernatants is caused by excreted melanin.

**Figure 15:**
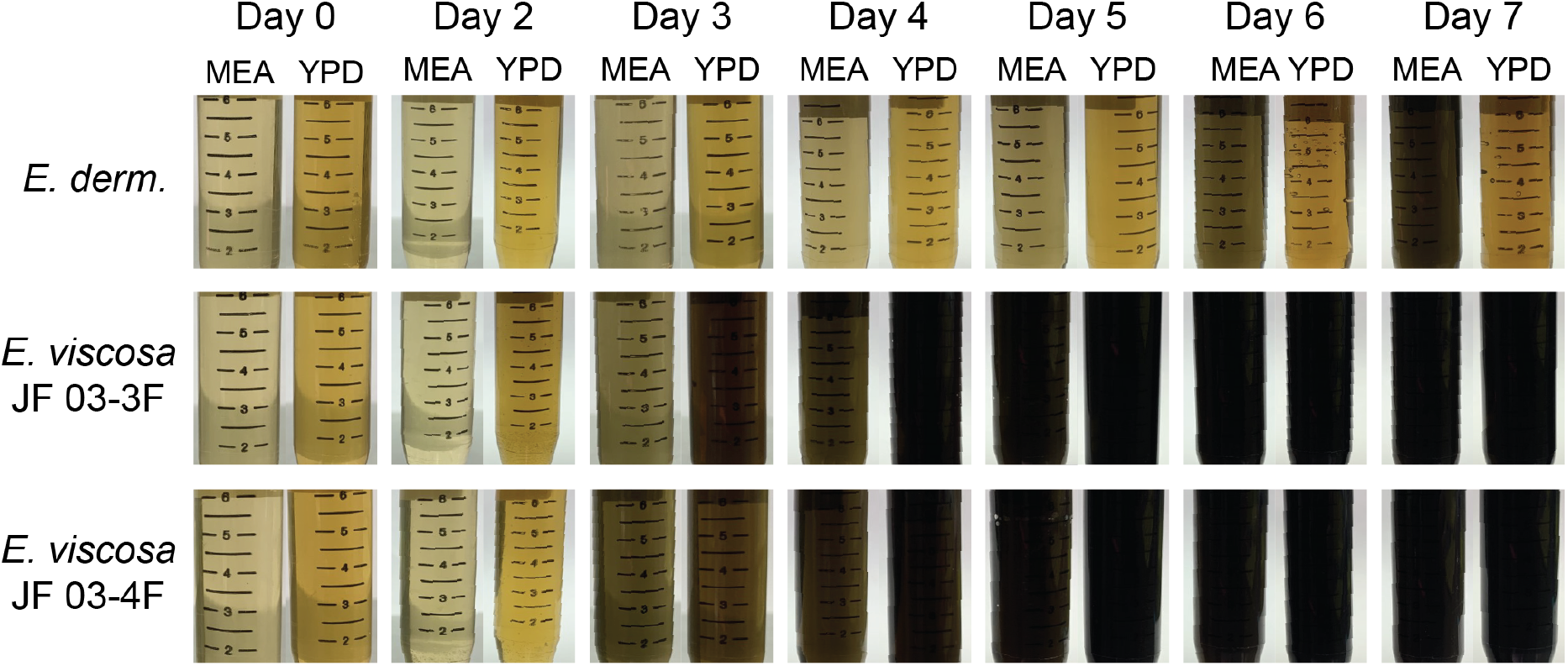
Supernatants of *E. dermatitidis (E. derm.)*, *E. viscosa* JF 03-3F, and *E. viscosa* JF 03-4F for each day to observe their accumulation of external melanin. Melanin began to accumulate in the supernatant of *E. viscosa* JF 03-3F and *E. viscosa* JF 03-4F on day 3, with a steep increase in melanin in YPD. *E. dermatitidis* began melanin secretion at day 6, and only in MEA.

**Figure 16:**
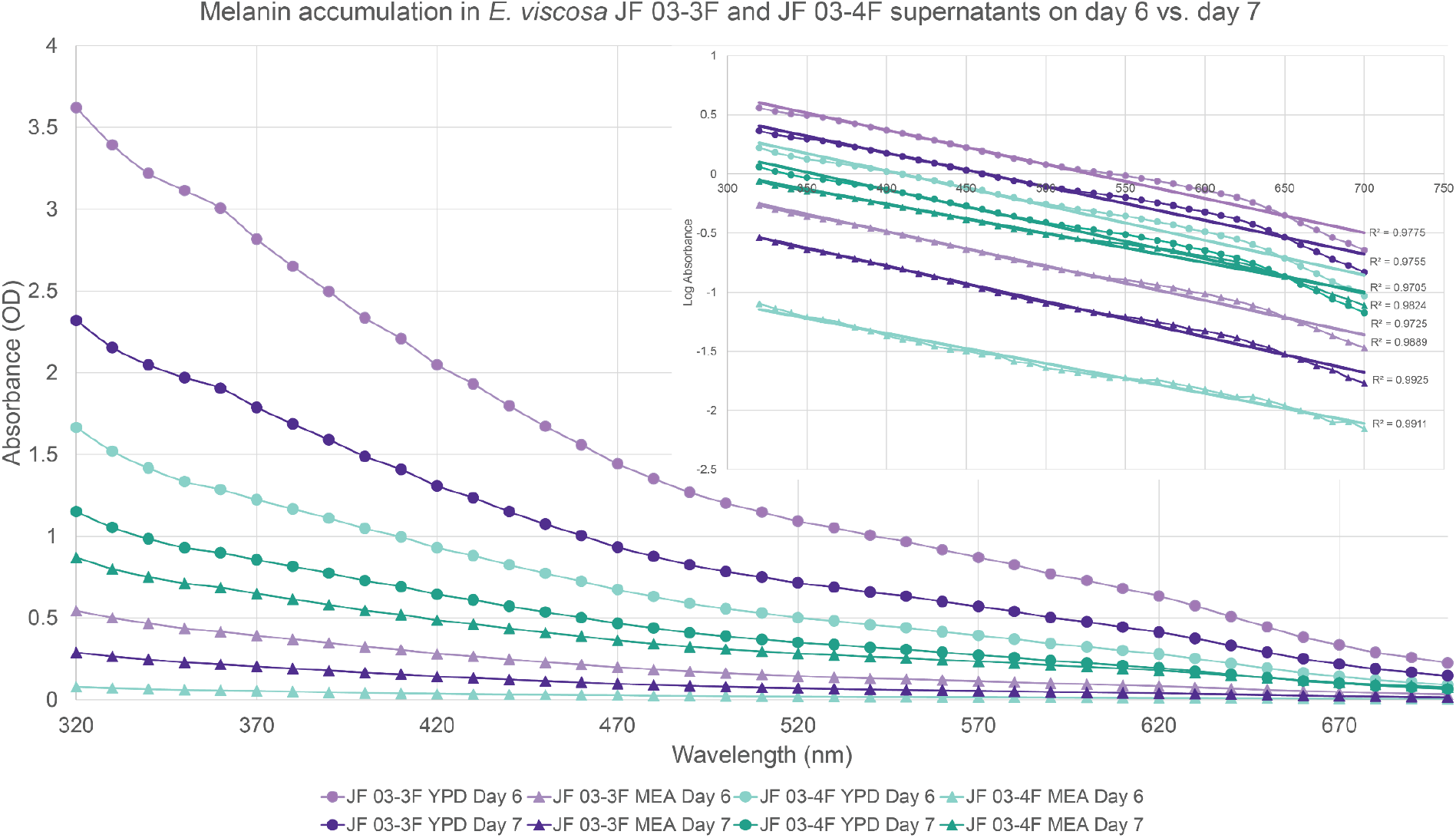
Day 6 and 7 results of daily melanin extraction from *E. viscosa* JF 03-3F and *E. viscosa* JF 03-4F. Both samples display typical melanin properties with full spectrum light, in that all samples have a linear regression when their Log Absorbance is plotted against wavelength, with an R^2^ value of 0.97 or higher (Pralea et al., 2019). The sample with the highest amount of secreted melanin was *E. viscosa* JF 03-3F in YPD on day 6. Both *E. viscosa* JF 03-3F and *E. viscosa* JF 03-4F had more secreted melanin when grown on YPD as opposed to MEA which showed lower melanin secretion for both species. Interestingly, both species had a decrease in melanin on day 7 when grown on YPD.

### Genetic Analysis of Melanin Production

#### Creation of an Albino Mutant Using Chemical Mutagenesis

At this time, molecular tools for the manipulation of *E. viscosa* JF 03-3F and *E. viscosa* JF 03-4F are not yet available. Therefore, we combined classical genetics with genome re-sequencing to initially investigate the regulation of melanin production. Following mutagenesis with the alkylating agent ethyl methanesulfonate (EMS), four distinct phenotypic classes of mutants were recovered: pink/albino, crusty, brown, and melanin over-excretion (data not shown). At this time, the most important of these phenotypes was the pink/albino phenotype found in a mutant of *E. viscosa* JF 03-4F, a mutant we named EMS 2-11 (Figure 17). Genome re-sequencing of high molecular weight DNA from mutant EMS 2- 11 and comparison to the reference *E. viscosa* JF 03-4F genome revealed a deleterious SNP causing a nonsense mutation in the gene polyketide synthase 1 (*pks1*). This mutation was on position 2,346,590 in scaffold 3, located in the *pks1* gene, which caused a C -> T mutation on the first position of the codon such that it became a stop codon, Q814STOP. Interestingly, *pks1* only functions in one of the three melanin biosynthetic pathways found in *E. viscosa* JF 03-4F, specifically the DHN/allomelanin pathway, which is typically known as the main melanin produced by polyextremotolerant fungi. Only two of the other random mutations found in EMS 2-11 were missense or nonsense mutations. One is a mutation in a transcriptional repressor EZH1, and another is in alcohol dehydrogenase GroES-like/polyketide synthase enolreductase. Although either of these mutations could theoretically contribute to the pink phenotype, it is more likely that a nonsense mutation in *pks1* is solely responsible for the loss of melanin production despite the lack of mutations in the other melanin biosynthetic pathways. This mutant is considered albino but is colored pink because of carotenoid production (Noack-Schönmann et al., 2014; Tesei et al., 2017). It has already been shown that blockage of melanin production in *E. dermatitidis* by mutations in *pks1* results in pink instead of white albino colonies because they also produce carotenoids and by-products such as flavolin that are normally masked by melanin (Geis & Szaniszlo, 1984; Geis et al., 1984).

**Figure 17:**
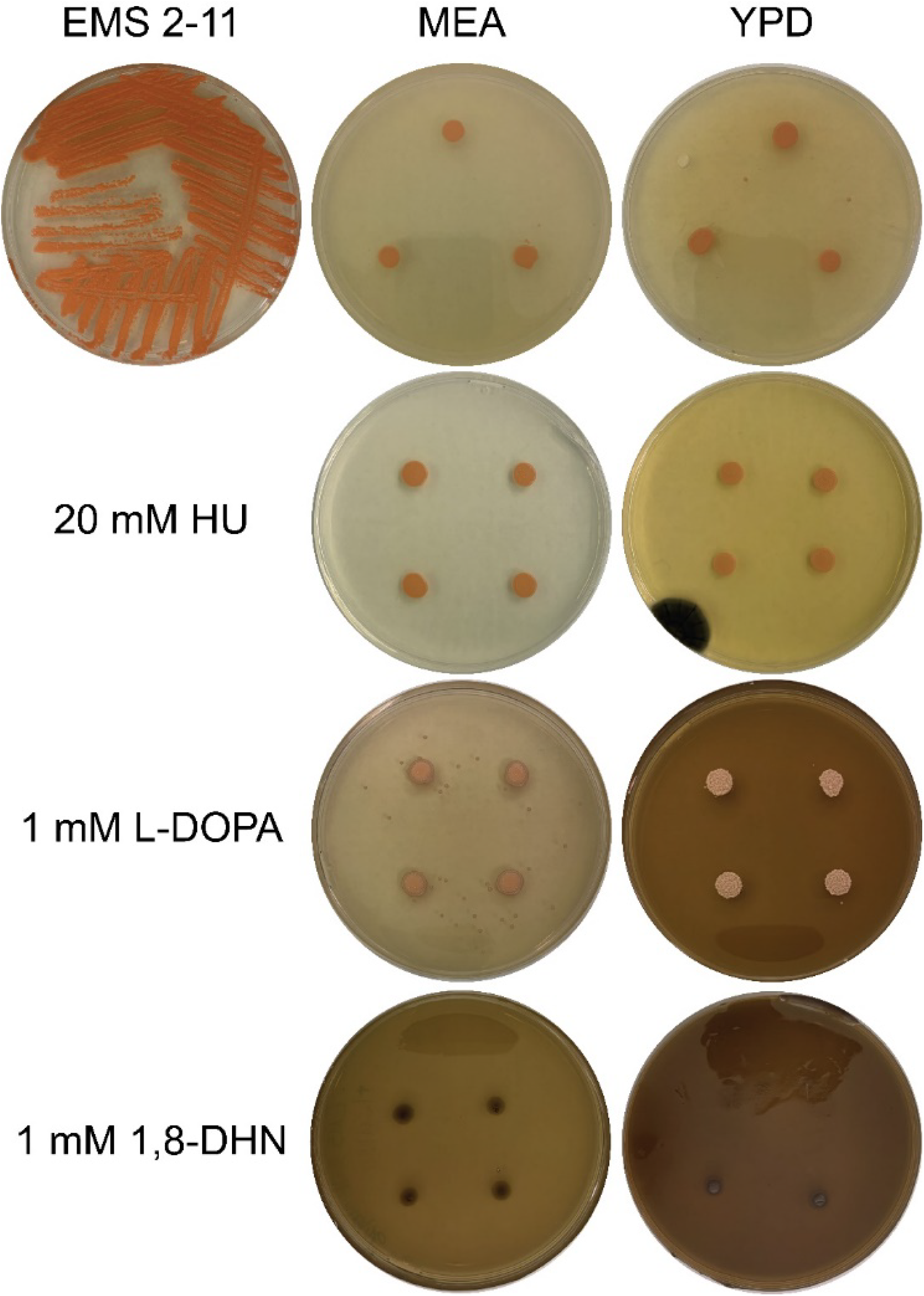
Phenotype of the EMS 2-11 mutant of *E. viscosa* JF 03-4F with *pks1* nonsense mutation Q814STOP, causing melanin production to be stopped hence the pink coloration. Attempts to recover melanin production were done with Hydroxyurea (HU), L-DOPA, and 1,8-DHN. Neither HU nor L-DOPA were able to recover melanin production in the mutant, however 1,8-DHN was able to recover melanin production in this mutant.

#### Recovery of melanin Production in an Albino Mutant: Modern use of the Classical Beadle and Tatum Biochemical-Genetics Experiment

Pks1 is a polyketide synthase that is essential for the first step in the DHN-melanin/allomelanin production pathway. However, both *E. viscosa* JF 03-4F and *E. viscosa* JF 03-3F are shown to have the genetic capabilities to produce all three forms of fungal melanins, which means theoretically any of the three pathways could be activated to produce melanin if the fungi are given the proper chemical precursors. To test if “activation” of any of the three melanin production pathways could restore melanin production to the EMS 2-11 mutant, we performed the classical Beadle and Tatum biochemical-genetics experiment (Beadle & Tatum, 1941) and substituted either 1 mM of L-DOPA, 20 mM hydroxyurea (HU), tyrosine (additive of 0.5 g/L, 1 g/L, or 0.5 g/L sole nitrogen source), or 1 mM of 1,8- DHN into the medium. L-DOPA was previously shown to recover melanization in a *pks1* deletion mutant of *E. dermatitidis*, thus we presumed that it would be taken up by our EMS 2-11 mutant and activate the DOPA melanin/pheomelanin biosynthesis pathway (Dadachova et al., 2007; Paolo et al., 2006). However, substitution of 1 mM L-DOPA into either MEA or YPD did not induce melanin production in the EMS 2-11 *pks1* mutant (Figure 17). These plates were grown in the dark for 10 days, as L-DOPA will auto polymerize into a melanin precursor in the presence of light, and still no melanin production was observed.

Addition of tyrosine, a melanin precursor molecule for L-DOPA and HGA-based melanins, also did not recover EMS 2-11’s melanin production in any of the media iterations tested (Figure S10). Hydroxyurea is another compound that in other *Exophiala* species induced melanin production (Schultzhaus et al., 2020), however addition of 20 mM of HU also did not activate melanin production in our EMS 2-11 albino mutant (Figure 17). Finally, addition to the media of 1 mM of 1,8-DHN, which is the immediate precursor to the final DHN-melanin/allomelanin molecule and is downstream of *pks1*, did result in recovery of melanized colonies. The colonies that grew on MEA+ 1 mM 1,8-DHN did not form extensive growth, but their cells were still dark, and on YPD + 1,8-DHN active dark colony growth is observed (Figure 17). These results suggest that the DHN-melanin produced by the Pks1 polyketide synthase is responsible for the bulk of melanin production by *E. viscosa* JF 03-4F.

## Discussion

A major limitation to our understanding of the roles played by BSCs in semi-arid ecosystems is our lack of insight into the microbial components of these biofilms and the nature of their functional interactions (Carr et al., 2021). As photoautotrophs, the roles of cyanobacteria and algae in BSCs are clear (Belnap, 2002), but the importance of other residents such as fungi and heterotrophic bacteria is less so. Here, we identify two isolates of a novel species of black yeasts from BSCs; *E. viscosa* two strains JF 03-3F and JF 03-4F. In addition to presenting the complete annotated genome sequence for both strains, we also provide a relatively detailed profile of their ability to utilize diverse carbon and nitrogen sources, as well as their response to different forms of stress. Most importantly, we demonstrate that *E. viscosa* JF 03-3F and *E. viscosa* JF 03-4F are capable of producing excessive amounts of melanin, including melanin that is excreted at high levels. Our results suggest that by making melanin extracellularly available as a public good, polyextremotolerant fungi provide a critical service to the broader BSC community.

### E. viscosa and its Closest Relatives

*E. viscosa* JF 03-3F and *E. viscosa* JF 03-4F are two strains of a novel fungus from the family Herpotrichielleaceae that we isolated from a lichen-dominated BSC. Distinguishing features of these isolates includes their yeast-dominated morphology, pronounced melanization, and production of extracellular material that leads to the formation of “goopy” or “slimy” colonies. Notably, numerous other melanized fungi with filamentous or polymorphic characteristics were also isolated from the same BSC communities. These fungi, which will be described in detail elsewhere, grow much more slowly than *E. viscosa* JF 03-3F or *E. viscosa* JF 03-4F and also release much less or no melanin.

Phylogenetic comparisons suggest that the closest relatives of *E. viscosa* JF 03-3F and *E. viscosa* JF 03-4F are *E. sideris*, *E. bergeri*, *P. elegans*, and *E*. *nigra*. However, through extensive phylogenetic analyses we can confirm that *E. viscosa* is a distinct species separate from these related fungi. Comparison of the two strains respective annotated genome sequences revealed that overall gene content and characteristics are similar to *E. sideris*. *E*. *sideris* belongs to the *jeanselmei*-clade of *Exophiala*; it is typically recovered from highly arsenate-polluted environments and is not known to be pathogenic (Seyedmousavi et al., 2011). The failure of *E. viscosa* JF 03-3F and *E. viscosa* JF 03-4F to grow at 37 °C reinforces the point that species within this clade are less likely to possess the capacity of causing disease in mammals.

Organization of the *MAT* locus between both strains of *E. viscosa* and *E. sideris* is very similar, as they possess a fused *MAT* gene, although they differ in the transcript directionality for the flanking *COX13* and *APC5* genes. Many *Exophiala* species are considered heterothallic, even species closely related to *E. sideris* such as *E. oligosperma* and *E. spinifera* are heterothallic (Teixeira et al., 2017; Untereiner & Naveau, 1999). As previously proposed by Teixeira et al., the proposed homothallic behavior of *E. sideris*, *E. viscosa* JF 03-3F, and *E. viscosa* JF 03-4F, may have resulted from an ancestral parasexual recombination event that altered the organization of the *MAT* locus (2017). Further investigation of these ideas will require demonstration that these *Exophiala* species undergo sexual development. Additional investigations will also be performed to determine the environmental factors for triggering sexual or asexual development in *E. viscosa*, as past attempts of established methods have failed but do not constitute evidence for absence of development.

### Phenotypic Characterization of E. viscosa JF 03-3F and E. viscosa JF 03-4F

Our results show that *E. viscosa* JF 03-3F and *E. viscosa* JF 03-4F are capable of using a diverse array of carbon sources. These include mannitol, ribitol, and glucose, which are considered the main carbon sources exchanged between photobionts and their mycobiont partners in lichens (Richardson et al., 1967; Yoshino et al., 2020). Although, *E. viscosa* JF 03-3F and *E. viscosa* JF 03-4F are able to use most nitrogen sources provided, glutamate and aspartate are clearly not being taken up or used, unlike in yeasts such as *S. cerevisiae* where glutamate is a more favorable nitrogen source used for multiple core metabolism and amino acid production processes (Ljungdahl & Daignan-Fornier, 2012). Because peptone is a favorable nitrogen source for *E. viscosa* JF 03-3F and *E. viscosa* JF 03-4F, it is likely that this species can import peptides and degrade them as organic nitrogen sources. In the BSC soil environment, organic nitrogen has been shown to be the more abundant form of nitrogen (Delgado-Baquerizo et al., 2010), but whether cyanobacteria and other nitrogen-fixing bacteria in BSCs are contributing organic or inorganic nitrogen sources is not understood (Belnap, 2002). However, since these fungi can use both inorganic and organic forms of nitrogen, they can use their metabolic flexibility to benefit them when either are present.

A characteristic feature of BSCs is the inherently stressful nature of these ecosystems. Of note, *E. viscosa* JF 03-3F and *E. viscosa* JF 03-4F were isolated from a site that can experience extremes in temperature, UV exposure, and soil wetness/osmotic pressure. Although the optimal growth range for these strains is from 15 °C to 27 °C, they are capable of slow growth at 4 °C and while they do not grow at 37 °C, they do remain viable after 72 consecutive hours of exposure to this temperature. The ability to grow at 4 °C, albeit slowly, presumably contributes to the success of fungi such *E. viscosa* JF 03-3F and *E. viscosa* JF 03-4F in adapting to BSCs in colder drylands. A broader comparison of microbiomes and associated functional traits between cold-adapted BSCs and those found in hot deserts would be an informative approach to identifying microbial species and/or physiological factors that underlie temperature adaptation.

As for resistance to UV radiation, *E. viscosa* JF 03-3F and *E. viscosa* JF 03-4F were both able to survive the high exposure of UV-C and were uniquely resistant to standard UV mutagenesis conditions. Potential mechanisms that might contribute to UV resistance include the presence of melanin or other defensive compounds (i.e., carotenoids), some of which could be public goods provided by other constituents of the BSC community, or an enhanced capacity to repair damaged DNA. This will ultimately be an interesting topic to investigate in the future.

One feature we expected to find in *E. viscosa* JF 03-3F and *E. viscosa* JF 03-4F, but we did not necessarily observe was higher metal resistance. Melanin is known to enhance resistances to metals (Bell & Wheeler, 1986; Gadd, 1994; Gorbushina, 2007; Purvis et al., 2004), and their closest relative *E. sideris* is primarily isolated from arsenate-contaminated soils (Seyedmousavi et al., 2011), so we predicted that this species would have increased metal tolerance when compared to non-melanized fungi. However, of the metals tested, only AgNO_3_ and CuCl_2_ showed a decreased zone of clearing (higher resistance) in *E. viscosa* JF 03-3F and *E. viscosa* JF 03-4F, compared to *S. cerevisiae* and *E. dermatitidis*. A potential explanation for the latter observation is that melanin has a higher affinity for copper due to the phenolic compounds within the polymer (Gadd & de Rome, 1988).

### Production and Excretion of Melanin by E. viscosa JF 03-3F and E. viscosa JF 03-4F

Melanin is considered an expensive secondary metabolite to produce because it is made up of conglomerates of polyphenols and as such contains many carbons (Schroeder et al., 2020). Therefore, it would be assumed that an organism would not want to create melanin just to export it out of the cell. Either this excreted melanin is an evolutionary trait that is beneficial to the fungi in some way, or this melanin excretion is a form of overflow metabolism in which melanin is the most streamlined way to remove an excessive buildup of precursors (De Deken, 1966). Additionally, there are not many fungi known to excrete melanin and few in the genus *Exophiala* (Romsdahl et al., 2021), so the possibility of a novel species that excretes such extreme amounts of melanin into their surroundings was intriguing.

Both fungi have the genetic makeup to produce all three types of fungal melanin: allomelanin (DHN-derived), pyomelanin (HGA/L-tyrosine derived), and pheomelanin (L-DOPA-derived); but it is still unknown how these melanin biosynthetic pathways are regulated. We were able to confirm through genetic means that they are producing DHN-derived allomelanin, but we were unable to definitively confirm the production of the other two forms of melanin. Additionally, we can speculate that tyrosine-derived melanins are not being directly produced for excretion under conditions we have tested, since using tyrosine as a sole nitrogen source did not induce melanin excretion. However, tyrosine used as an additive did induce melanin excretion at 0.5 g/L but not at 1 g/L, implying a possible tyrosine-dependent regulation of melanin production/excretion that is not entirely understood. We hope to eventually elucidate the melanin(s) that are produced by these fungi by combining genetic approaches with mass spectrometry and NMR in the future.

In our attempts to understand the regulation of melanin production by *E. viscosa* JF 03-3F and *E. viscosa* JF 03-4F we tried blocking individual pathways using known chemical blockers of DHN and L-DOPA-melanin, phthalide and kojic acid respectively. Neither blocker, nor their combination at the highest amounts used were able to block the production of melanin in either strain. If this species has the capability to produce all three melanins, then it is possible that they are producing more than one at the same time. This could be the case here, since the combination of both kojic acid and phthalide still resulted in melanized cells which would indicate that tyrosine-derived pyomelanin was the cause of melanization. Alternatively, since these fungi are fully melanized, it could be that the concentrations used on fungi that use melanin in asexual structures only was not strong enough to block melanin production in these fungi. Additionally, our melanin gene annotation revealed that both fungi contain homologs to *A. niger* and *A. fumigatus’* Arp1 and Arp2, which could be causing a bifurcation of the DHN melanin pathway and prevent phthalide from being an effective blocker of melanin production (Jørgensen et al., 2011; Wheeler & Klich, 1995).

Linked regulation of potentially multiple forms of melanin, was observed with an EMS-induced mutant in which the *pks1* gene was disrupted. Although *pks1* is the first enzyme involved in only the production of DHN-derived allomelanin, the cells that resulted from this mutation were albino. Theoretically, these cells should have still been capable of producing the two other melanins as their pathways were seemingly not disrupted by any mutations, but this was not the case. Addition of L-DOPA has been shown to induce melanization in a *pks1* deletion mutant of *E. dermatitidis* (Paolo et al., 2006), which would indicate that it is capable of producing both pheomelanin and allomelanin. However, the same experiment did not restore melanization to the *E. viscosa* JF 03-4F’s *pks1* nonsense mutant. Therefore, it is possible that; (i) *E. viscosa* JF 03-3F and *E. viscosa* JF 03-4F only produce DHN-derived allomelanin and disruption of *pks1* causes all melanin production to shut down, (ii) *pks1* (or downstream creation of DHN-derived allomelanin) is essential for the regulation of other melanin productions, or (iii) pheomelanin and pyomelanin are only produced under specific conditions that were not tested during analysis of the mutants.

Our experiments also showed that *E. viscosa* JF 03-3F and *E. viscosa* JF 03-4F actively excrete melanin when they are viable and growing. We observed that melanin began to accumulate starting as early as day 2 (by OD reads), therefore suggesting live cells are excreting melanin and not melanin coming out of dead lysed cells. However, at this time we do not have a mechanism for how melanin is being released from the cells, and something to be investigated later. We also observed a decrease in melanin concentrations after day 6 in both YPD and MEA, which raises the intriguing potential for melanin uptake or degradation by *E. viscosa* JF 03-3F and *E. viscosa* JF 03-4F. Additionally, we observed that increasing organic nitrogen in the media induced melanin excretion from both fungi, but we do not know how this phenomenon is regulated or which melanin is being excreted.

### Insights into the Niche of Polyextremotolerant Fungi Within the Biological Soil Crust Consortium

Fungal residents of BSC communities include lichenized mycobionts, lichenicolous fungi, endolichenic fungi, and free-living fungi. Although the latter are likely not directly associated with lichens themselves, they are surrounded by algae, cyanobacteria, and other bacteria in the same way that lichen-associated fungi are surrounded by a similar community of microbes. This raises the intriguing possibility that fungi such as *E. viscosa* JF 03-3F and *E. viscosa* JF 03-4F might transiently associated with autotrophs in response to specific environmental conditions. Members of the genus *Exophiala* (Class Eurotiomycetes) are phylogenetically related to the lichen-forming Verrucariales, and reside between Lecanoromycetes and Lichinomycetes (James et al., 2006). Additionally, the lifestyle of the ancestral state of Eurotiomycetes is currently debated as to whether it is lichenized or not. Originally, it was thought that the ancestral state of Eurotiomycetidae was lichenized (Lutzoni et al., 2001), but recent literature has shown that the ancestral state was more likely non-lichenized (Lücking & Nelsen, 2018; Nelsen et al., 2020). However, many have recognized the similarities between taxa residing in Eurotiomycetes that are non-lichenized to their lichenized relatives. Particularly, members of Eurotiomycetes and Lecanoromycetes share commonality in their ecological locations (Gostinčar et al., 2012; Muggia et al., 2013) and in their type 1 polyketide synthase (PKS) genes (Boustie & Grube, 2005), which are thought to be acquired by lichen-forming fungi via horizontal gene transfer from actinobacteria (Schmitt & Lumbsch, 2009). As such, it is likely that *E. viscosa* JF 03-3F and *E. viscosa* JF 03-4F and other related fungi have lichen-like qualities, which allow them to interact with algae, cyanobacteria, and other bacteria to form lichen-like symbioses. The idea that polyextremotolerant fungi can engage in lichen-like associations with microbes has been postulated (Gostinčar et al., 2012; Muggia et al., 2013), but direct experimentation is still needed to confirm these ideas. As these are two strains of a newly identified species, more extensive ecological and molecular investigation is needed to determine the extent to which this taxon is found in BSCs and to elucidate their species interaction networks.

Correlation between environmental organic nitrogen availability and excreted melanin amounts could have strong implications for interactions between these fungi found in BSCs and their nitrogen-fixing community partners. The presumed source of nitrogen used to enhance the excretion of melanin would be from nitrogen-fixing prokaryotes such as cyanobacteria, other bacteria, and archaea in the community. Additionally, organic nitrogen was found to be more abundant than inorganic nitrogen in BSC soils, and more abundant than in other soil ecosystems (Delgado-Baquerizo et al., 2010). However, the nitrogen source that is released into the environment by nitrogen-fixing bacteria in BSCs is not known. It could be ammonia/ammonium, or it could be an organic form of nitrogen since cyanobacteria and diazotrophs are known to release both forms of nitrogen (Berthelot et al., 2015; Gimenez et al., 2016; Vitousek et al., 2002). Our results suggest that the availability of these nitrogen-fixers to produce a pool of organic nitrogen could subsequently induce the excretion of melanin that then provides protective services to the broader BSC community. Excreted melanin could function at the community-level to maintain respiration and growth under varying temperature conditions while also mitigating the effects of desiccation and exposure to UV, extending the window of time allotted for optimal photosynthetic capabilities (Carr et al., 2021; Honegger, 1997; Honegger & Haisch, 2001; Lange & Tenhunen, 1981; Nybakken et al., 2004). Moreover, the excreted melanin could simultaneously serve as an external source of carbon used by the fungi and possibly others when the environment does not allow for photosynthesis. In either scenario, the seemingly wasteful excretion of melanin by *E. viscosa* JF 03-3F and *E. viscosa* JF 03-4F instead represents the optimal sharing of an expensive public good for the overall betterment of the BSC community in which they reside.

The availability of complete, annotated genome sequences for *E. viscosa* JF 03-3F and *E. viscosa* JF 03-4F will enable more detailed investigations of the mechanisms that support the adaptation of these fungi to extreme environments. This in turn should provide greater insight into the roles that these fungi play in the BSC community. Of particular interest will be the regulatory mechanisms that coordinate the production of different types of melanin in response to environmental and chemical inputs. Moreover, the broad applicability of melanin as a bioproduct has triggered growing interest in microorganisms that excrete this polymer. In this context, *E. viscosa* JF 03-3F and *E. viscosa* JF 03-4F could potentially serve as excellent platforms for additional engineering focused on optimizing or tailoring the type of melanin produced in responses to specific applications.

## Acknowledgements

The authors would like to acknowledge Dr. Rajib Saha for providing a pre-print review of this work, with his guidance we were able to make this manuscript as clear and concise as possible, Dr. Christian Elowsky for acquiring beautiful microscopic and plate photos of these new strains, Dr. Tania Kurbessoian for assistance with phylogenetic analyses, and Darwyn S. Coxson for assistance in obtaining soil samples for isolation of these fungi. The work (proposal: 10.46936/10.25585/60001081) conducted by the U.S. Department of Energy Joint Genome Institute (https://ror.org/04xm1d337), a DOE Office of Science User Facility, is supported by the Office of Science of the U.S. Department of Energy under Contract No. DE-AC02-05CH11231. Funding for this work was provided by NASA grant number 80NSSC17K0737, and NSF PRFB fellowship number 2209217.

## Citations

Altschul, S. F., Gish, W., Miller, W., Myers, E. W., & Lipman, D. J. (1990). Basic local alignment search tool. J Mol Biol, 215(3), 403–410. https://doi.org/10.1016/S0022-2836(05)80360-2

Ametrano, C. G., Selbmann, L., & Muggia, L. (2017). A standardized approach for co-culturing Dothidealean rock-inhabiting fungi and lichen photobionts in vitro. Symbiosis, 73(1), 35–44.

Ashburner, M., Ball, C. A., Blake, J. A., Botstein, D., Butler, H., Cherry, J. M., Sherlock, G. (2000). Gene ontology: tool for the unification of biology. The Gene Ontology Consortium. Nat Genet, 25(1), 25–29. https://doi.org/10.1038/75556

Bairoch, A. (2000). The ENZYME database in 2000. Nucleic Acids Res, 28(1), 304–305. https://doi.org/10.1093/nar/28.1.304

Bairoch, A., Apweiler, R., Wu, C. H., Barker, W. C., Boeckmann, B., Ferro, S., Yeh, L. S. (2005). The Universal Protein Resource (UniProt). Nucleic Acids Res, 33(Database issue), D154-159. https://doi.org/10.1093/nar/gki070

Bateman, A., Coin, L., Durbin, R., Finn, R. D., Hollich, V., Griffiths-Jones, S., Eddy, S. R. (2004). The Pfam protein families database. Nucleic Acids Res, 32(Database issue), D138-141. https://doi.org/10.1093/nar/gkh121

Bates, S. T., Garcia-Pichel, F., & Nash III, T. (2010). Fungal components of biological soil crusts: insights from culture-dependent and culture-independent studies. Bibliotheca Lichenologica, 105, 197–210.

Bates, S. T., Reddy, G. S., & Garcia-Pichel, F. (2006). Exophiala crusticola anam. nov.(affinity Herpotrichiellaceae), a novel black yeast from biological soil crusts in the Western United States. International journal of systematic and evolutionary microbiology, 56(11), 2697–2702.

Beadle, G. W., & Tatum, E. L. (1941). Genetic Control of Biochemical Reactions in Neurospora. Proceedings of the National Academy of Sciences, 27(11), 499–506. https://doi.org/10.1073/pnas.27.11.499

Bell, A. A., & Wheeler, M. H. (1986). Biosynthesis and functions of fungal melanins. Annual review of phytopathology, 24(1), 411–451.

Belnap, J. (2002). Nitrogen fixation in biological soil crusts from southeast Utah, USA. Biology and fertility of soils, 35(2), 128–135.

Belnap, J. (2003). The world at your feet: desert biological soil crusts. Frontiers in Ecology and the Environment, 1(4), 181–189.

Belnap, J., Büdel, B., & Lange, O. L. (2001). Biological soil crusts: characteristics and distribution. In Biological soil crusts: structure, function, and management (pp 3–30). Springer.

Belnap, J., & Eldridge, D. (2001). Disturbance and recovery of biological soil crusts. In Biological soil crusts: structure, function, and management (pp 363–383). Springer.

Belnap, J., & Lange, O. L. (2003). Biological soil crusts: structure, function, and management (Vol. 150). Springer.

Berthelot, H., Bonnet, S., Camps, M., Grosso, O., & Moutin, T. (2015). Assessment of the dinitrogen released as ammonium and dissolved organic nitrogen by unicellular and filamentous marine diazotrophic cyanobacteria grown in culture. Frontiers in Marine Science, 2, 80.

Birney, E., & Durbin, R. (2000). Using GeneWise in the Drosophila annotation experiment. Genome Res, 10(4), 547–548. https://doi.org/10.1101/gr.10.4.547

Bligh, E. G., & Dyer, W. J. (1959). A rapid method of total lipid extraction and purification. Canadian journal of biochemistry and physiology, 37(8), 911–917.

Boustie, J., & Grube, M. (2005). Lichens—a promising source of bioactive secondary metabolites. Plant Genetic Resources, 3(2), 273–287. https://doi.org/10.1079/pgr200572

Bowker, M. A., Reed, S., Belnap, J., & Phillips, S. (2002). Temporal variation in community composition, pigmentation, and Fv/Fm of desert cyanobacterial soil crusts. Microbial Ecology, 13-25.

Bowker, M. A., Soliveres, S., & Maestre, F. T. (2010). Competition increases with abiotic stress and regulates the diversity of biological soil crusts. Journal of Ecology, 98(3), 551–560.

Bowman, E. A., & Arnold, A. E. (2021). Drivers and implications of distance decay differ for ectomycorrhizal and foliar endophytic fungi across an anciently fragmented landscape. The ISME Journal, 15(12), 3437–3454. https://doi.org/10.1038/s41396-021-01006-9

Cao, W., Zhou, X., McCallum, N. C., Hu, Z., Ni, Q. Z., Kapoor, U., Mantanona, A. J. (2021). Unraveling the structure and function of melanin through synthesis. Journal of the American Chemical Society, 143(7), 2622–2637.

Carr, E. C., Harris, S. D., Herr, J. R., & Riekhof, W. R. (2021). Lichens and biofilms: Common collective growth imparts similar developmental strategies. Algal Research, 54, 102217.

Castresana, J. (2000). Selection of conserved blocks from multiple alignments for their use in phylogenetic analysis. Molecular biology and evolution, 17(4), 540–552.

Chen, Z., Martinez, D. A., Gujja, S., Sykes, S. M., Zeng, Q., Szaniszlo, P. J., Cuomo, C. A. (2014). Comparative Genomic and Transcriptomic Analysis of Wangiella dermatitidis, A Major Cause of Phaeohyphomycosis and a Model Black Yeast Human Pathogen. G3 Genes|Genomes|Genetics, 4(4), 561–578. https://doi.org/10.1534/g3.113.009241

Cordero, R. J., & Casadevall, A. (2017). Functions of fungal melanin beyond virulence. Fungal Biology Reviews, 31(2), 99–112.

Couradeau, E., Giraldo-Silva, A., De Martini, F., & Garcia-Pichel, F. (2019). Spatial segregation of the biological soil crust microbiome around its foundational cyanobacterium, Microcoleus vaginatus, and the formation of a nitrogen-fixing cyanosphere. Microbiome, 7(1). https://doi.org/10.1186/s40168-019-0661-2

Cubero, O. F., Crespo, A., Fatehi, J., & Bridge, P. D. (1999). DNA extraction and PCR amplification method suitable for fresh, herbarium-stored, lichenized, and other fungi. Plant Systematics and Evolution, 216(3), 243–249.

Dadachova, E., Bryan, R. A., Huang, X., Moadel, T., Schweitzer, A. D., Aisen, P., Casadevall, A. (2007). Ionizing radiation changes the electronic properties of melanin and enhances the growth of melanized fungi. PloS one, 2(5), e457.

De Deken, R. H. (1966). The Crabtree Effect: A Regulatory System in Yeast. Journal of General Microbiology, 44(2), 149–156. https://doi.org/10.1099/00221287-44-2-149

De Hoog, G., Vicente, V., Caligiorne, R., Kantarcioglu, S., Tintelnot, K., Gerrits van den Ende, A., & Haase, G. (2003). Species diversity and polymorphism in the Exophiala spinifera clade containing opportunistic black yeast-like fungi. Journal of Clinical Microbiology, 41(10), 4767–4778.

Delgado-Baquerizo, M., Castillo-Monroy, A. P., Maestre, F. T., & Gallardo, A. (2010). Plants and biological soil crusts modulate the dominance of N forms in a semi-arid grassland. Soil Biology and Biochemistry, 42(2), 376–378. https://doi.org/10.1016/j.soilbio.2009.11.003

Donlan, R. M., & Costerton, J. W. (2002). Biofilms: survival mechanisms of clinically relevant microorganisms. Clinical microbiology reviews, 15(2), 167–193.

Edberg, S., Chaskes, S., Alture-Werber, E., & Singer, J. (1980). Esculin-based medium for isolation and identification of Cryptococcus neoformans. Journal of Clinical Microbiology, 12(3), 332–335.

Edberg, S. C., Pittman, S., & Singer, J. (1977). Esculin hydrolysis by Enterobacteriaceae. Journal of Clinical Microbiology, 6(2), 111–116.

Edgar, R. C. (2004). MUSCLE: multiple sequence alignment with high accuracy and high throughput. Nucleic Acids Research, 32(5), 1792–1797. https://doi.org/10.1093/nar/gkh340

Frases, S., Salazar, A., Dadachova, E., & Casadevall, A. (2007). Cryptococcus neoformans can utilize the bacterial melanin precursor homogentisic acid for fungal melanogenesis. Applied and environmental microbiology, 73(2), 615–621.

Gadd, G. M. (1994). Interactions of fungi with toxic metals. The Genus Aspergillus, 361–374.

Gadd, G. M., & de Rome, L. (1988). Biosorption of copper by fungal melanin. Applied microbiology and biotechnology, 29(6), 610–617.

Geis, P. A., & Szaniszlo, P. J. (1984). Carotenoid pigments of the dematiaceous fungus Wangiella dermatitidis. Mycologia, 76(2), 268–273.

Geis, P. A., Wheeler, M. H., & Szaniszlo, P. J. (1984). Pentaketide metabolites of melanin synthesis in the dematiaceous fungus Wangiella dermatitidis. Archives of microbiology, 137(4), 324–328.

Gessler, N., Egorova, A., & Belozerskaya, T. (2014). Melanin pigments of fungi under extreme environmental conditions. Applied Biochemistry and Microbiology, 50(2), 105–113.

Gimenez, A., Baklouti, M., Bonnet, S., & Moutin, T. (2016). Biogeochemical fluxes and fate of diazotroph- derived nitrogen in the food web after a phosphate enrichment: modeling of the VAHINE mesocosms experiment. Biogeosciences, 13(17), 5103–5120. https://doi.org/10.5194/bg-13-5103-2016

Gnerre, S., MacCallum, I., Przybylski, D., Ribeiro, F. J., Burton, J. N., Walker, B. J., Jaffe, D. B. (2011). High-quality draft assemblies of mammalian genomes from massively parallel sequence data. Proceedings of the National Academy of Sciences, 108(4), 1513–1518. https://doi.org/10.1073/pnas.1017351108

Gorbushina, A. A. (2007). Life on the rocks. Environmental microbiology, 9(7), 1613–1631.

Gostinčar, C., Grube, M., De Hoog, S., Zalar, P., & Gunde-Cimerman, N. (2009). Extremotolerance in fungi: evolution on the edge. FEMS microbiology ecology, 71(1), 2–11.

Gostinčar, C., Grube, M., & Gunde-Cimerman, N. (2011). Evolution of fungal pathogens in domestic environments? Fungal biology, 115(10), 1008–1018.

Gostinčar, C., Muggia, L., & Grube, M. (2012). Polyextremotolerant black fungi: oligotrophism, adaptive potential, and a link to lichen symbioses. Frontiers in Microbiology, 3, 390.

Grabherr, M. G., Haas, B. J., Yassour, M., Levin, J. Z., Thompson, D. A., Amit, I., Zeng, Q. (2011). Full-length transcriptome assembly from RNA-Seq data without a reference genome. Nature biotechnology, 29(7), 644–652.

Grigoriev, I. V., Nikitin, R., Haridas, S., Kuo, A., Ohm, R., Otillar, R., Shabalov, I. (2014). MycoCosm portal: gearing up for 1000 fungal genomes. Nucleic Acids Research, 42(D1), D699–D704. https://doi.org/10.1093/nar/gkt1183

Grigoriev, I. V., Nordberg, H., Shabalov, I., Aerts, A., Cantor, M., Goodstein, D., Dubchak, I. (2012). The genome portal of the Department of Energy Joint Genome Institute. Nucleic Acids Res, 40(Database issue), D26-32. https://doi.org/10.1093/nar/gkr947

Grube, M., Cernava, T., Soh, J., Fuchs, S., Aschenbrenner, I., Lassek, C., Sensen, C. W. (2015). Exploring functional contexts of symbiotic sustain within lichen-associated bacteria by comparative omics. The ISME journal, 9(2), 412–424.

Haase, G., Sonntag, L., Melzer-Krick, B., & de Hoog, G. S. (1999). Phylogenetic inference by SSU-gene analysis of members of the Herpotrichiellaceae with special reference to human pathogenic species. Stud Mycol, 43, 80–97.

Hom, E. F., & Murray, A. W. (2014). Niche engineering demonstrates a latent capacity for fungal-algal mutualism. Science, 345(6192), 94–98.

Honegger, R. (1997). Metabolic interactions at the mycobiont-photobiont interface in lichens. In Plant relationships (pp. 209–221). Springer.

Honegger, R., & Haisch, A. (2001). Immunocytochemical location of the (1→ 3)(1→ 4)-β-glucan lichenin in the lichen-forming ascomycete Cetraria islandica (Icelandic moss). New phytologist, 150(3), 739–746.

Huelsenbeck, J. P., & Ronquist, F. (2001). MRBAYES: Bayesian inference of phylogenetic trees. Bioinformatics, 17(8), 754–755.

Jalmi, P., Bodke, P., Wahidullah, S., & Raghukumar, S. (2012). The fungus Gliocephalotrichum simplex as a source of abundant, extracellular melanin for biotechnological applications. World Journal of Microbiology and Biotechnology, 28(2), 505–512. https://doi.org/10.1007/s11274-011-0841-0

James, T. Y., Kauff, F., Schoch, C. L., Matheny, P. B., Hofstetter, V., Cox, C. J., Miadlikowska, J. (2006). Reconstructing the early evolution of Fungi using a six-gene phylogeny. Nature, 443(7113), 818–822.

Jurka, J. (2000). Repbase update: a database and an electronic journal of repetitive elements. Trends Genet, 16(9), 418–420. https://doi.org/10.1016/s0168-9525(00)02093-x

Jørgensen, T. R., Park, J., Arentshorst, M., van Welzen, A. M., Lamers, G., vanKuyk, P. A., Ram, A. F. J. (2011). The molecular and genetic basis of conidial pigmentation in Aspergillus niger. Fungal Genetics and Biology, 48(5), 544–553. https://doi.org/10.1016/j.fgb.2011.01.005

Katoh, K., Misawa, K., Kuma, K. i., & Miyata, T. (2002). MAFFT: a novel method for rapid multiple sequence alignment based on fast Fourier transform. Nucleic acids research, 30(14), 3059–3066.

Kearse, M., Moir, R., Wilson, A., Stones-Havas, S., Cheung, M., Sturrock, S., Drummond, A. (2012). Geneious Basic: an integrated and extendable desktop software platform for the organization and analysis of sequence data. Bioinformatics, 28(12), 1647–1649. https://doi.org/10.1093/bioinformatics/bts199

Kobayashi, N., Muramatsu, T., Yamashina, Y., Shirai, T., Ohnishi, T., & Mori, T. (1993). Melanin reduces ultraviolet-induced DNA damage formation and killing rate in cultured human melanoma cells. Journal of investigative dermatology, 101(5), 685–689.

Koonin, E. V., Fedorova, N. D., Jackson, J. D., Jacobs, A. R., Krylov, D. M., Makarova, K. S., Natale, D. A. (2004). A comprehensive evolutionary classification of proteins encoded in complete eukaryotic genomes. Genome Biol, 5(2), R7. https://doi.org/10.1186/gb-2004-5-2-r7

Krogh, A., Larsson, B., von Heijne, G., & Sonnhammer, E. L. (2001). Predicting transmembrane protein topology with a hidden Markov model: application to complete genomes. J Mol Biol, 305(3), 567–580. https://doi.org/10.1006/jmbi.2000.4315

Kumar, S., Stecher, G., Li, M., Knyaz, C., & Tamura, K. (2018). MEGA X: Molecular Evolutionary Genetics Analysis across Computing Platforms. Mol Biol Evol, 35(6), 1547–1549. https://doi.org/10.1093/molbev/msy096

Kuo, A., Bushnell, B., & Grigoriev, I. V. (2014). Fungal genomics: sequencing and annotation. Advances in Botanical Research, 70, 1–52.

Kõljalg, U., Nilsson, R. H., Abarenkov, K., Tedersoo, L., Taylor, A. F. S., Bahram, M., Larsson, K.-H. (2013). Towards a unified paradigm for sequence-based identification of fungi. Molecular Ecology, 22(21), 5271–5277. https://doi.org/10.1111/mec.12481

Lan, S., Wu, L., Zhang, D., & Hu, C. (2012). Successional stages of biological soil crusts and their microstructure variability in Shapotou region (China). Environmental Earth Sciences, 65(1), 77–88.

Lange, O., & Tenhunen, J. (1981). Moisture content and CO2 exchange of lichens. II. Depression of net photosynthesis in Ramalina maciformis at high water content is caused by increased thallus carbon dioxide diffusion resistance. Oecologia, 51(3), 426–429.

Letunic, I., & Bork, P. (2021). Interactive Tree Of Life (iTOL) v5: an online tool for phylogenetic tree display and annotation. Nucleic acids research, 49(W1), W293–W296.

Ljungdahl, P. O., & Daignan-Fornier, B. (2012). Regulation of amino acid, nucleotide, and phosphate metabolism in Saccharomyces cerevisiae. Genetics, 190(3), 885–929.

Lutzoni, F., Pagel, M., & Reeb, V. (2001). Major fungal lineages are derived from lichen symbiotic ancestors. Nature, 411(6840), 937–940.

Lücking, R., & Nelsen, M. P. (2018). Chapter 23 - Ediacarans, Protolichens, and Lichen-Derived Penicillium: A Critical Reassessment of the Evolution of Lichenization in Fungi. In M. Krings, C. J. Harper, N. R. Cúneo, & G. W. Rothwell (Eds.), Transformative Paleobotany (pp. 551–590). Academic Press. https://doi.org/10.1016/B978-0-12-813012-4.00023-1

Maier, S., Muggia, L., Kuske, C. R., & Grube, M. (2016). Bacteria and non-lichenized fungi within biological soil crusts. In Biological Soil Crusts: An Organizing Principle in Drylands (pp. 81–100). Springer.

McFarland, J. (1907). The nephelometer: an instrument for estimating the number of bacteria in suspensions used for calculating the opsonic index and for vaccines. Journal of the American Medical Association, 49(14), 1176–1178.

Mitchison-Field, L. M. Y., Vargas-Muñiz, J. M., Stormo, B. M., Vogt, E. J. D., Van Dierdonck, S., Pelletier, J. F., Gladfelter, A. S. (2019). Unconventional Cell Division Cycles from Marine-Derived Yeasts. Current Biology, 29(20), 3439–3456.e3435. https://doi.org/10.1016/j.cub.2019.08.050

Molina, L., & Molina, M. (1986). Absolute absorption cross sections of ozone in the 185-to 350-nm wavelength range. Journal of Geophysical Research: Atmospheres, 91(D13), 14501–14508.

Moussa, T. A., Al-Zahrani, H. S., Kadasa, N. M., Moreno, L. F., Gerrits van den Ende, A., de Hoog, G. S., & Al-Hatmi, A. M. (2017). Nomenclatural notes on Nadsoniella and the human opportunist black yeast genus Exophiala. Mycoses, 60(6), 358–365.

Muggia, L., Gueidan, C., Knudsen, K., Perlmutter, G., & Grube, M. (2013). The Lichen Connections of Black Fungi. Mycopathologia, 175(5-6), 523–535. https://doi.org/10.1007/s11046-012-9598-8

Nelsen, M. P., Lücking, R., Boyce, C. K., Lumbsch, H. T., & Ree, R. H. (2020). No support for the emergence of lichens prior to the evolution of vascular plants. Geobiology, 18(1), 3–13. https://doi.org/10.1111/gbi.12369

Nielsen, H., Brunak, S., & von Heijne, G. (1999). Machine learning approaches for the prediction of signal peptides and other protein sorting signals. Protein Eng, 12(1), 3–9. https://doi.org/10.1093/protein/12.1.3

Nilsson, R. H., Larsson, K. H., Taylor, A. F. S., Bengtsson-Palme, J., Jeppesen, T. S., Schigel, D., Abarenkov, K. (2019). The UNITE database for molecular identification of fungi: handling dark taxa and parallel taxonomic classifications. Nucleic Acids Res, 47(D1), D259–D264. https://doi.org/10.1093/nar/gky1022

Noack-Schönmann, S., Bus, T., Banasiak, R., Knabe, N., Broughton, W. J., Den Dulk-Ras, H., Gorbushina, A. A. (2014). Genetic transformation of Knufia petricola A95-a model organism for biofilm-material interactions. Amb Express, 4(1), 1–6. https://doi.org/10.1186/s13568-014-0080-5

Nunes da Rocha, U., Cadillo-Quiroz, H., Karaoz, U., Rajeev, L., Klitgord, N., Dunn, S., Garcia-Pichel, F. (2015). Isolation of a significant fraction of non-phototroph diversity from a desert biological soil crust. Frontiers in microbiology, 6, 277.

Nybakken, L., Solhaug, K. A., Bilger, W., & Gauslaa, Y. (2004). The lichens Xanthoria elegans and Cetraria islandica maintain a high protection against UV-B radiation in Arctic habitats. Oecologia, 140(2), 211–216.

Ogata, H., Goto, S., Sato, K., Fujibuchi, W., Bono, H., & Kanehisa, M. (1999). KEGG: Kyoto Encyclopedia of Genes and Genomes. Nucleic Acids Res, 27(1), 29–34. https://doi.org/10.1093/nar/27.1.29

Pal, A. K., Gajjar, D. U., & Vasavada, A. R. (2014). DOPA and DHN pathway orchestrate melanin synthesis in Aspergillus species. Medical mycology, 52(1), 10–18.

Paolo, W. F., Dadachova, E., Mandal, P., Casadevall, A., Szaniszlo, P. J., & Nosanchuk, J. D. (2006). Effects of disrupting the polyketide synthase gene WdPKS1 in Wangiella [Exophiala] dermatitidis on melanin production and resistance to killing by antifungal compounds, enzymatic degradation, and extremes in temperature. BMC microbiology, 6(1), 1–16. https://doi.org/10.1186/1471-2180-6-55

Pearson, W. R. (2013). An Introduction to Sequence Similarity (“Homology”) Searching. Current Protocols in Bioinformatics, 42(1), 3.1.1–3.1.8. https://doi.org/10.1002/0471250953.bi0301s42

Perez-Cuesta, U., Aparicio-Fernandez, L., Guruceaga, X., Martin-Souto, L., Abad-Diaz-de-Cerio, A., Antoran, A., Rementeria, A. (2020). Melanin and pyomelanin in Aspergillus fumigatus: from its genetics to host interaction. International Microbiology, 23(1), 55–63.

Pralea, I.-E., Moldovan, R.-C., Petrache, A.-M., Ilieș, M., Hegheș, S.-C., Ielciu, I., Radu, M. (2019). From extraction to advanced analytical methods: The challenges of melanin analysis. International journal of molecular sciences, 20(16), 3943.

Price, A. L., Jones, N. C., & Pevzner, P. A. (2005). De novo identification of repeat families in large genomes. Bioinformatics, 21 *Suppl 1*, i351–358. https://doi.org/10.1093/bioinformatics/bti1018

Price, M. N., Dehal, P. S., & Arkin, A. P. (2009). FastTree: computing large minimum evolution trees with profiles instead of a distance matrix. Molecular biology and evolution, 26(7), 1641–1650.

Puillandre, N., Brouillet, S., & Achaz, G. (2021). ASAP: assemble species by automatic partitioning. Molecular Ecology Resources, 21(2), 609–620. https://doi.org/10.1111/1755-0998.13281

Purvis, O. W., Bailey, E. H., McLean, J., Kasama, T., & Williamson, B. J. (2004). Uranium biosorption by the lichen Trapelia involuta at a uranium mine. Geomicrobiology Journal, 21(3), 159–167.

Pócs, T. (2009). Cyanobacterial crust types, as strategies for survival in extreme habitats. Acta Botanica Hungarica, 51(1-2), 147–178.

Płonka, P., & Grabacka, M. (2006). Melanin synthesis in microorganisms: biotechnological and medical aspects. Acta Biochimica Polonica, 53(3).

Rajeev, L., Da Rocha, U. N., Klitgord, N., Luning, E. G., Fortney, J., Axen, S. D., Kerfeld, C. A. (2013). Dynamic cyanobacterial response to hydration and dehydration in a desert biological soil crust. The ISME journal, 7(11), 2178–2191.

Richardson, D., Smith, D., & Lewis, D. (1967). Carbohydrate movement between the symbionts of lichens. Nature, 214(5091), 879–882.

Ridgway, R. (1912). Color standards and color nomenclature. The author.

Romsdahl, J., Schultzhaus, Z., Cuomo, C. A., Dong, H., Abeyratne-Perera, H., Hervey, W. J., & Wang, Z. (2021). Phenotypic Characterization and Comparative Genomics of the Melanin-Producing Yeast Exophiala lecanii-corni Reveals a Distinct Stress Tolerance Profile and Reduced Ribosomal Genetic Content. Journal of Fungi, 7(12), 1078. https://doi.org/10.3390/jof7121078

Ronquist, F., & Huelsenbeck, J. P. (2003). MrBayes 3: Bayesian phylogenetic inference under mixed models. Bioinformatics, 19(12), 1572–1574.

Rosas, Á. L., Nosanchuk, J. D., Gómez, B. L., Edens, W. A., Henson, J. M., & Casadevall, A. (2000). Isolation and serological analyses of fungal melanins. *Journal of immunological methods*, *244*(1-2), 69-80. Salamov, A. A., & Solovyev, V. V. (2000). Ab initio gene finding in Drosophila genomic DNA. Genome Res, 10(4), 516–522. https://doi.org/10.1101/gr.10.4.516

Schmitt, I., & Lumbsch, H. T. (2009). Ancient Horizontal Gene Transfer from Bacteria Enhances Biosynthetic Capabilities of Fungi. PLoS ONE, 4(2), e4437. https://doi.org/10.1371/journal.pone.0004437

Schreier, W. J., Gilch, P., & Zinth, W. (2015). Early events of DNA photodamage. Annual review of physical chemistry, 66, 497–519.

Schroeder, W. L., Harris, S. D., & Saha, R. (2020). Computation-driven analysis of model polyextremo- tolerant fungus exophiala dermatitidis: defensive pigment metabolic costs and human applications. IScience, 23(4), 100980.

Schultzhaus, Z., Romsdahl, J., Chen, A., Tschirhart, T., Kim, S., Leary, D., & Wang, Z. (2020). The response of the melanized yeast Exophiala dermatitidis to gamma radiation exposure. Environmental microbiology, 22(4), 1310–1326.

Seyedmousavi, S., Badali, H., Chlebicki, A., Zhao, J., Prenafeta-Boldu, F. X., & De Hoog, G. S. (2011). Exophiala sideris, a novel black yeast isolated from environments polluted with toxic alkyl benzenes and arsenic. Fungal biology, 115(10), 1030–1037.

Teixeira, M. M., Moreno, L. F., Stielow, B. J., Muszewska, A., Hainaut, M., Gonzaga, L., de Hoog, G. S. (2017). Exploring the genomic diversity of black yeasts and relatives (Chaetothyriales, Ascomycota). Studies in mycology, 86, 1–28. https://doi.org/10.1016/j.simyco.2017.01.001

Ter-Hovhannisyan, V., Lomsadze, A., Chernoff, Y. O., & Borodovsky, M. (2008). Gene prediction in novel fungal genomes using an ab initio algorithm with unsupervised training. Genome Res, 18(12), 1979–1990. https://doi.org/10.1101/gr.081612.108

Tesei, D., Tafer, H., Poyntner, C., Piñar, G., Lopandic, K., & Sterflinger, K. (2017). Draft Genome Sequences of the Black Rock Fungus Knufia petricola and Its Spontaneous Nonmelanized Mutant. Genome Announc, 5(44). https://doi.org/10.1128/genomeA.01242-17

Tragiannidis, A., Fegeler, W., Rellensmann, G., Debus, V., Müller, V., Hoernig-Franz, I., Groll, A. (2012). Candidaemia in a European Paediatric University Hospital: a 10-year observational study. Clinical Microbiology and Infection, 18(2), E27–E30.

Untereiner, W. A. (1994). A simple method for the in vitro production of pseudothecia in species of Capronia. Mycologia, 86(2), 290–295. https://doi.org/10.1080/00275514.1994.12026410

Untereiner, W. A., & Naveau, F. A. (1999). Molecular systematics of the Herpotrichiellaceae with an assessment of the phylogenetic positions of Exophiala dermatitidis and Phialophora americana. Mycologia, 91(1), 67–83.

Untereiner, W. A., & Naveau, F. A. (1999). Molecular systematics of the Herpotrichiellaceae with an assessment of the phylogenetic positions of*Exophiala dermatitidis*and*Phialophora americana*. Mycologia, 91(1), 67–83. https://doi.org/10.1080/00275514.1999.12060994

Vitousek, P. M., Cassman, K., Cleveland, C., Crews, T., Field, C. B., Grimm, N. B., Sprent, J. I. (2002). Biogeochemistry, 57(1), 1–45. https://doi.org/10.1023/a:1015798428743

Weber, S. (2005). Light-driven enzymatic catalysis of DNA repair: a review of recent biophysical studies on photolyase. Biochimica et Biophysica Acta (BBA)-Bioenergetics, 1707(1), 1–23.

Wheeler, M. H., & Klich, M. A. (1995). The Effects of Tricyclazole, Pyroquilon, Phthalide, and Related Fungicides on the Production of Conidial Wall Pigments by Penicillium and Aspergillus Species. Pesticide Biochemistry and Physiology, 52(2), 125–136. https://doi.org/10.1006/pest.1995.1037

White, T. J., Bruns, T., Lee, S., & Taylor, J. (1990). 38 - AMPLIFICATION AND DIRECT SEQUENCING OF FUNGAL RIBOSOMAL RNA GENES FOR PHYLOGENETICS. In M. A. Innis, D. H. Gelfand, J. J. Sninsky, & T. J. White (Eds.), PCR Protocols (pp. 315–322). Academic Press. https://doi.org/10.1016/B978-0-12-372180-8.50042-1

Winston, F. (2008). EMS and UV mutagenesis in yeast. Current protocols in molecular biology, 82(1), 13.13 B. 11-13.13 B. 15. https://doi.org/10.1002/0471142727.mb1303bs82

Yoshino, K., Yamamoto, K., Masumoto, H., Degawa, Y., Yoshikawa, H., Harada, H., & Sakamoto, K. (2020). Polyol-assimilation capacities of lichen-inhabiting fungi. The Lichenologist, 52(1), 49–59.

Zanne, A. E., Abarenkov, K., Afkhami, M. E., Aguilar-Trigueros, C. A., Bates, S., Bhatnagar, J. M., Crowther, T. W. (2020). Fungal functional ecology: bringing a trait-based approach to plant-associated fungi. Biological Reviews, 95(2), 409–433.

Zdobnov, E. M., & Apweiler, R. (2001). InterProScan--an integration platform for the signature-recognition methods in InterPro. Bioinformatics, 17(9), 847–848. https://doi.org/10.1093/bioinformatics/17.9.847

Zeng, J. S., & De Hoog, G. S. (2008). *Exophiala spinifera*and its allies: diagnostics from morphology to DNA barcoding. Medical Mycology, 46(3), 193–208. https://doi.org/10.1080/13693780701799217

Zerbino, D. R., & Birney, E. (2008). Velvet: algorithms for de novo short read assembly using de Bruijn graphs. Genome research, 18(5), 821–829.

Zhou, K., Salamov, A., Kuo, A., Aerts, A. L., Kong, X., & Grigoriev, I. V. (2015). Alternative splicing acting as a bridge in evolution. Stem Cell Investig, 2, 19. https://doi.org/10.3978/j.issn.2306-9759.2015.10.01

Zupančič, J., Novak Babič, M., Zalar, P., & Gunde-Cimerman, N. (2016). The black yeast Exophiala dermatitidis and other selected opportunistic human fungal pathogens spread from dishwashers to kitchens. PLoS One, 11(2), e0148166.

