## Supplemental File 2 for "Characterization of a novel polyextremotolerant fungus, Exophiala viscosa, with insights into its melanin regulation and ecological niche"

ITS-only ASAP analysis:

Partition 1

Asap-Score: 1.000000

P-value: 2.814371e-01

Threshold distance: 1.32%

Nb subsets with recursion:18 (without recursion: 17)

------------------------------------------------------------

Subset[ 1 ] n: 2 ;id: Exophiala_viscosa_JF_03-4F Exophiala_viscosa_JF_03-3F

Subset[ 2 ] n: 1 ;id: Exophiala_nigra

Subset[ 3 ] n: 1 ;id: Exophiala_spinifera

Subset[ 4 ] n: 1 ;id: Exophiala_bergeri

Subset[ 5 ] n: 1 ;id: Exophiala_dermatitidis

Subset[ 6 ] n: 2 ;id: Exophiala_sideris Phaeoannellomyces_elegans

Subset[ 7 ] n: 1 ;id: Exophiala_lecanii-corni

Subset[ 8 ] n: 1 ;id: Exophiala_cancerae

Subset[ 9 ] n: 1 ;id: Exophiala_castellanii

Subset[ 10 ] n: 1 ;id: Exophiala_crusticola

Subset[ 11 ] n: 1 ;id: Exophiala_salmonis

Subset[ 12 ] n: 1 ;id: Exophiala_nishimurae

Subset[ 13 ] n: 1 ;id: Exophiala_pisciphila

Subset[ 14 ] n: 1 ;id: Exophiala_aquamarina

Subset[ 15 ] n: 1 ;id: Exophiala_xenobiotica

Subset[ 16 ] n: 1 ;id: Exophiala_oligosperma

Subset[ 17 ] n: 1 ;id: Exophiala_jeanselmei

Subset[ 18 ] n: 1 ;id: Endocarpon_pusillum

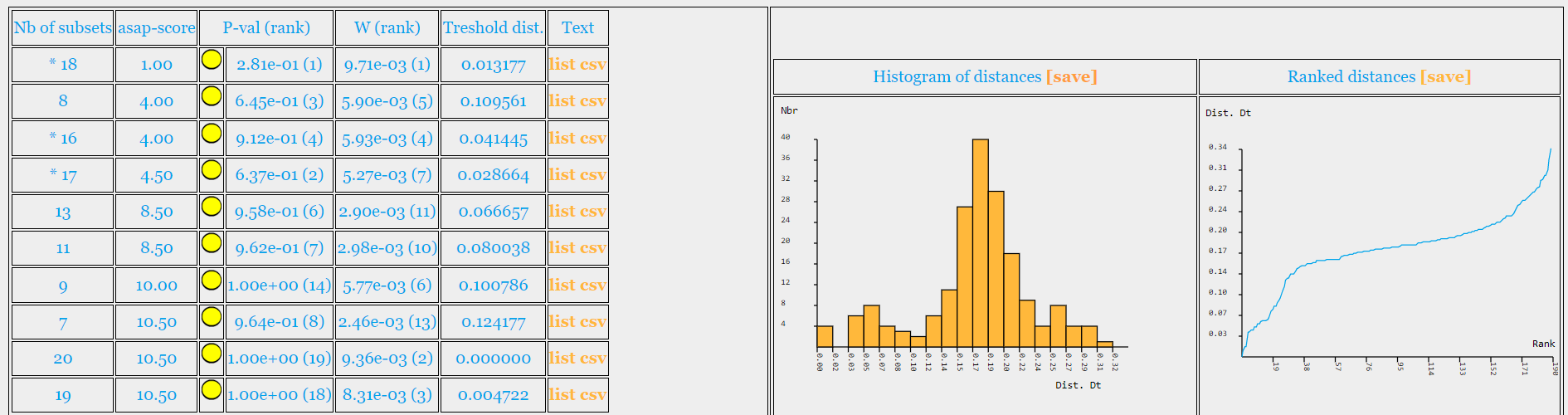

18S-28S-ITS concatenation ASAP analysis:

Partition 1

Asap-Score: 2.000000

P-value: 1.477046e-01

Threshold distance: 0.75%

Nb subsets with recursion:18 (without recursion: 17)

------------------------------------------------------------

Subset[ 1 ] n: 2 ;id: Exophiala viscosa JF 03-3F Exophiala viscosa JF 03-4F

Subset[ 2 ] n: 1 ;id: Exophiala nigra

Subset[ 3 ] n: 2 ;id: Exophiala sideris Phaeoannellomyces elegans

Subset[ 4 ] n: 1 ;id: Exophiala bergeri

Subset[ 5 ] n: 1 ;id: Exophiala spinifera

Subset[ 6 ] n: 1 ;id: Exophiala aquamarina

Subset[ 7 ] n: 1 ;id: Exophiala cancerae

Subset[ 8 ] n: 1 ;id: Exophiala castellanii

Subset[ 9 ] n: 1 ;id: Exophiala dermatitidis

Subset[ 10 ] n: 1 ;id: Exophiala jeanselmei

Subset[ 11 ] n: 1 ;id: Exophiala lecanii-corni

Subset[ 12 ] n: 1 ;id: Exophiala nishimurae

Subset[ 13 ] n: 1 ;id: Exophiala oligosperma

Subset[ 14 ] n: 1 ;id: Exophiala pisciphila

Subset[ 15 ] n: 1 ;id: Exophiala salmonis

Subset[ 16 ] n: 1 ;id: Exophiala xenobiotica

Subset[ 17 ] n: 1 ;id: Exophiala crusticola

Subset[ 18 ] n: 1 ;id: Endocarpon pusillum

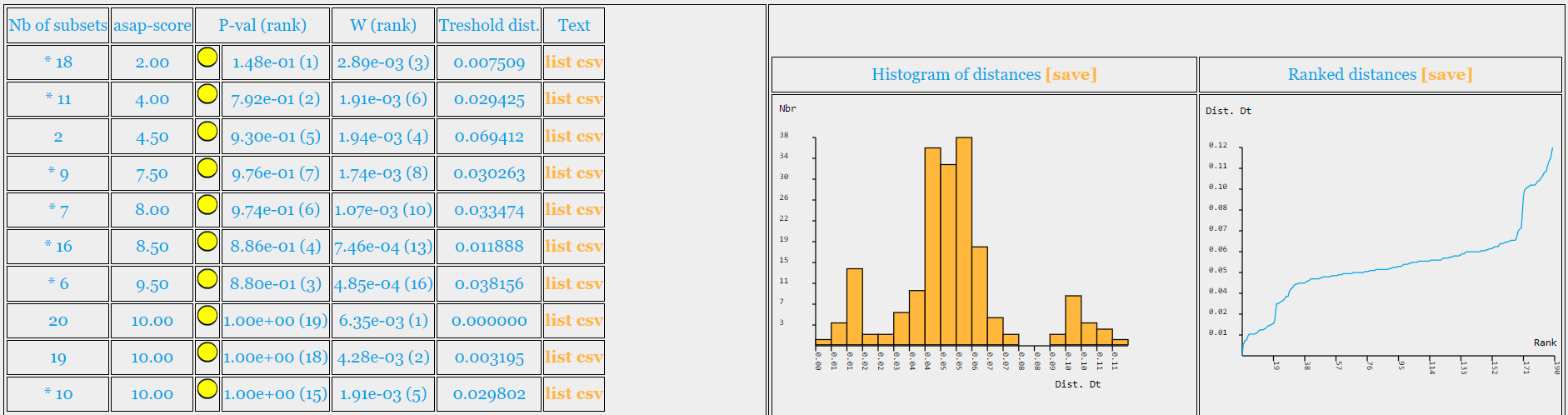

18S-28S-ITS-rpb1 concatenation ASAP analysis:

Partition 1

Asap-Score: 2.500000

P-value: 9.321357e-01

Threshold distance: 2.56%

Nb subsets with recursion:16 (without recursion: 15)

------------------------------------------------------------

Subset[ 1 ] n: 2 ;id: Exophiala viscosa JF 03-3F Exophiala viscosa JF 03-4F

Subset[ 2 ] n: 1 ;id: Exophiala sideris

Subset[ 3 ] n: 1 ;id: Exophiala bergeri

Subset[ 4 ] n: 1 ;id: Exophiala nigra

Subset[ 5 ] n: 1 ;id: Exophiala spinifera

Subset[ 6 ] n: 2 ;id: Exophiala aquamarina Exophiala pisciphila

Subset[ 7 ] n: 1 ;id: Exophiala salmonis

Subset[ 8 ] n: 1 ;id: Exophiala cancerae

Subset[ 9 ] n: 1 ;id: Exophiala castellanii

Subset[ 10 ] n: 1 ;id: Exophiala dermatitidis

Subset[ 11 ] n: 1 ;id: Exophiala jeanselmei

Subset[ 12 ] n: 1 ;id: Exophiala lecanii-corni

Subset[ 13 ] n: 1 ;id: Exophiala nishimurae

Subset[ 14 ] n: 1 ;id: Exophiala oligosperma

Subset[ 15 ] n: 1 ;id: Exophiala xenobiotica

Subset[ 16 ] n: 1 ;id: Endocarpon pusillum

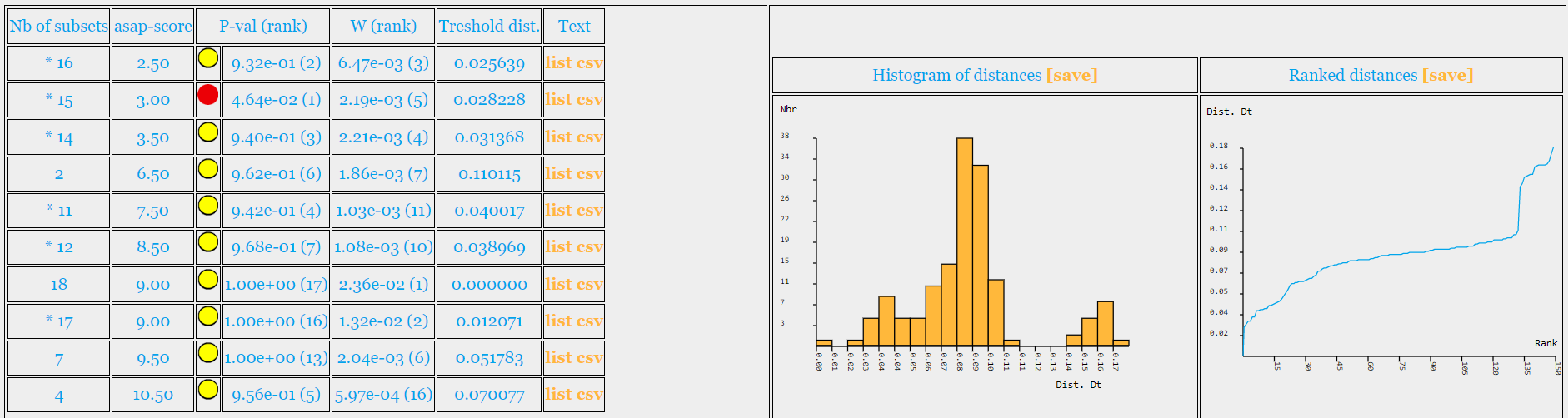
