## Supplemental Figures and Tables for "Characterization of a novel polyextremotolerant fungus, Exophiala viscosa, with insights into its melanin regulation and ecological niche"

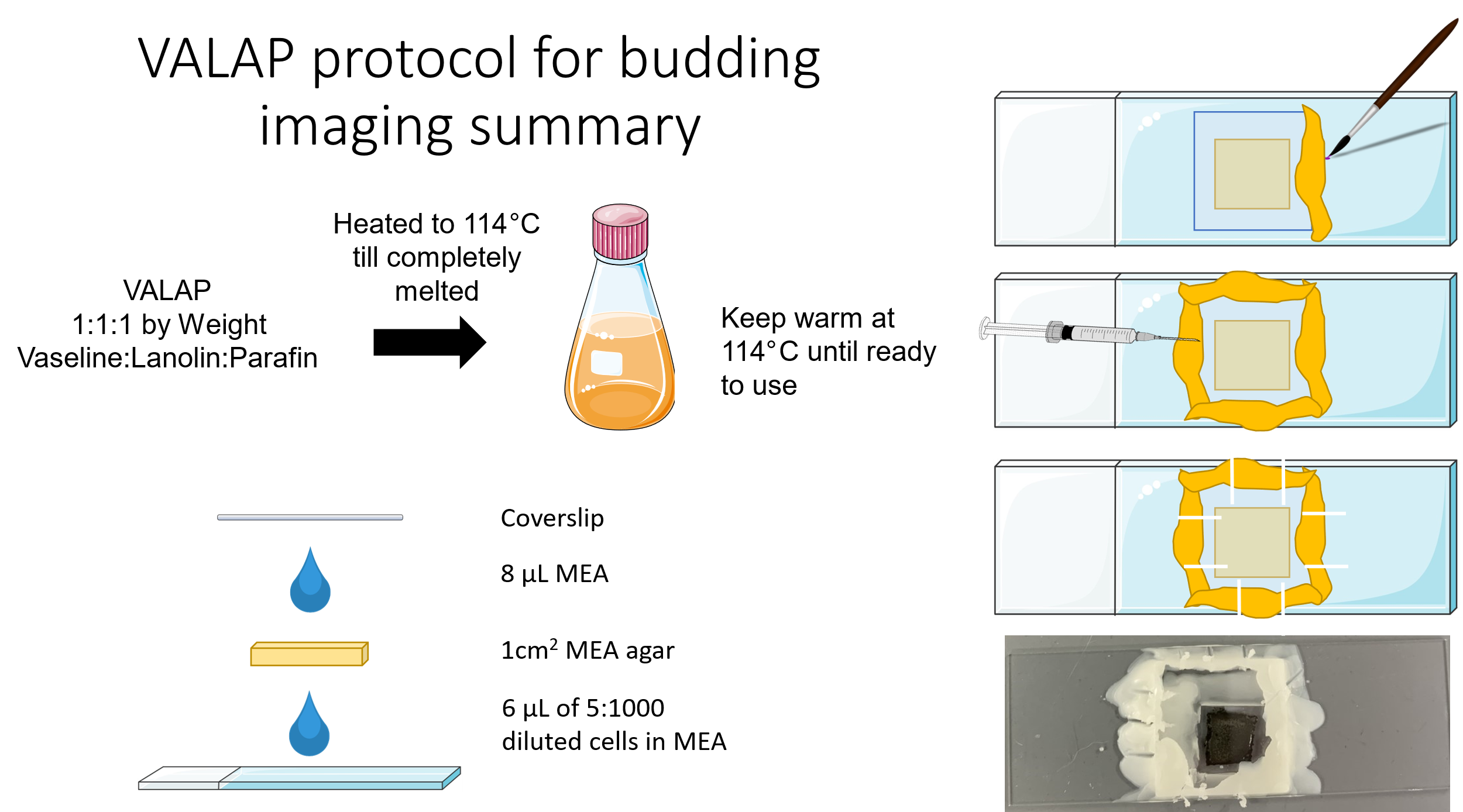

**Supplementary Figure 1:** Visual summary of the VALAP method for setting up microscope slides for long-term microscopy of budding patterns of yeast cells. VALAP is a 1:1:1 ratio by weight of Vaseline:Lanolin:Parafin which is heated to 114 °C to completely melt the mixture. After the mixture has melted, it must be maintained at 114 °C and not higher or else the mixture will become useless. To prepare the microscope slide, 6 μL of 5:1000 diluted cells is added directly onto the slide. Then a 1 cm2 slab of MEA agar is placed on top of the diluted cells. Additionally, 8 μL of liquid MEA is placed on top of the agar slab, and the cover slip is set on top of the liquid MEA. The warm VALAP is then applied to all of the outside edges of the coverslip using a paintbrush. Then using a needle syringe, air holes are carefully poked into the VALAP perimeter to allow for gas exchange while the cells are growing.

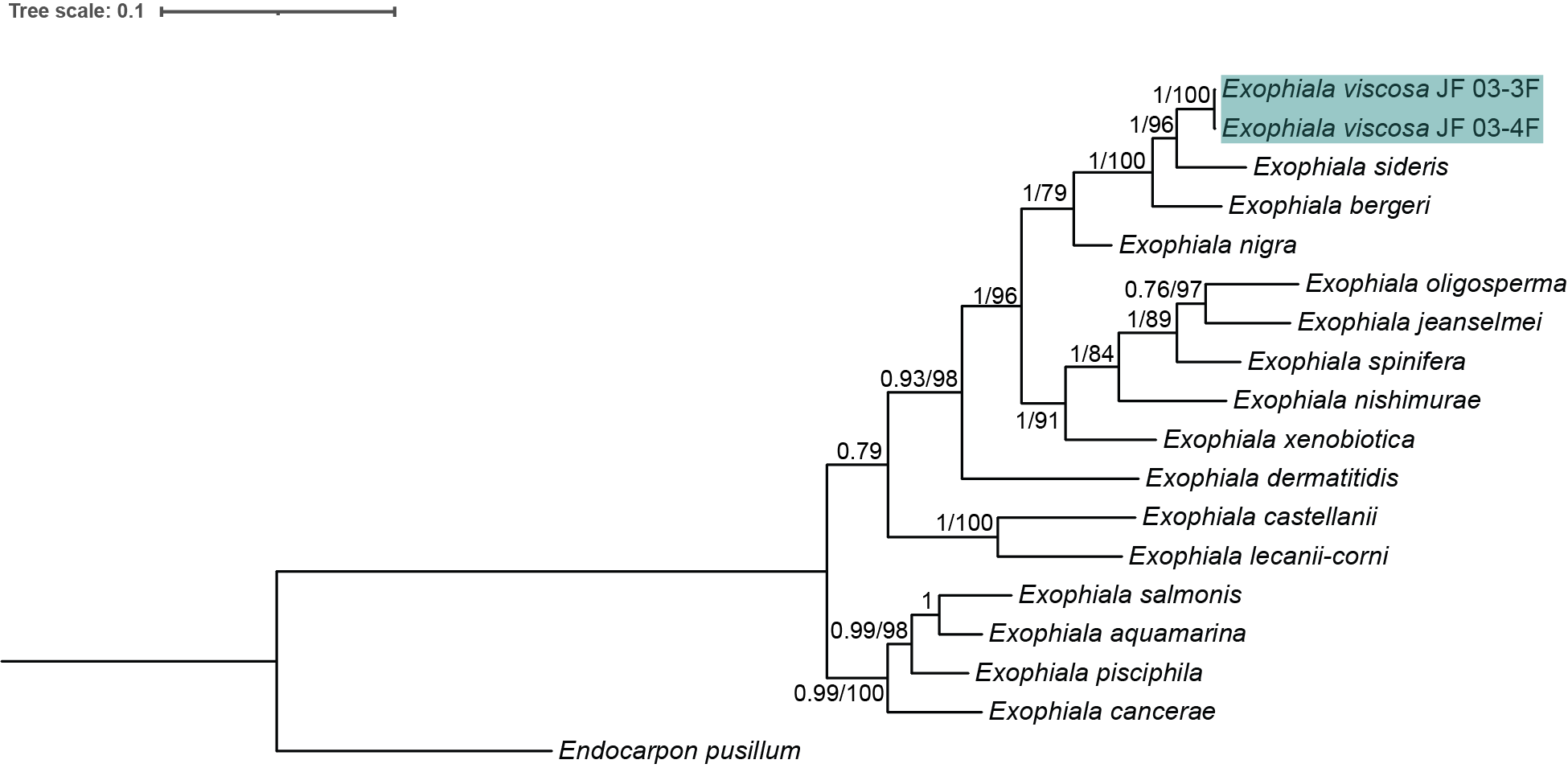

**Supplementary Figure 2**: Rooted Bayesian phylogenetic tree of the concatenation of the 18S, 28S, ITS, and RPB1 regions of available Exophiala species with the addition of E. viscosa JF 03-3F and E. viscosa JF 03-4F to identify the phylogenetic location of these new species within the genus Exophiala. The tree with the highest log likelihood is shown. The numbers above the branches are the Bayesian posterior probability/maximum likelihood bootstrap values, with posterior probabilities > 75% and bootstrap values > 80 shown. Location of E. viscosa JF 03-3F and E. viscosa JF 03-4F within Exophiala places them closest to E. sideris with 100% posterior probability and a bootstrap value of 96.

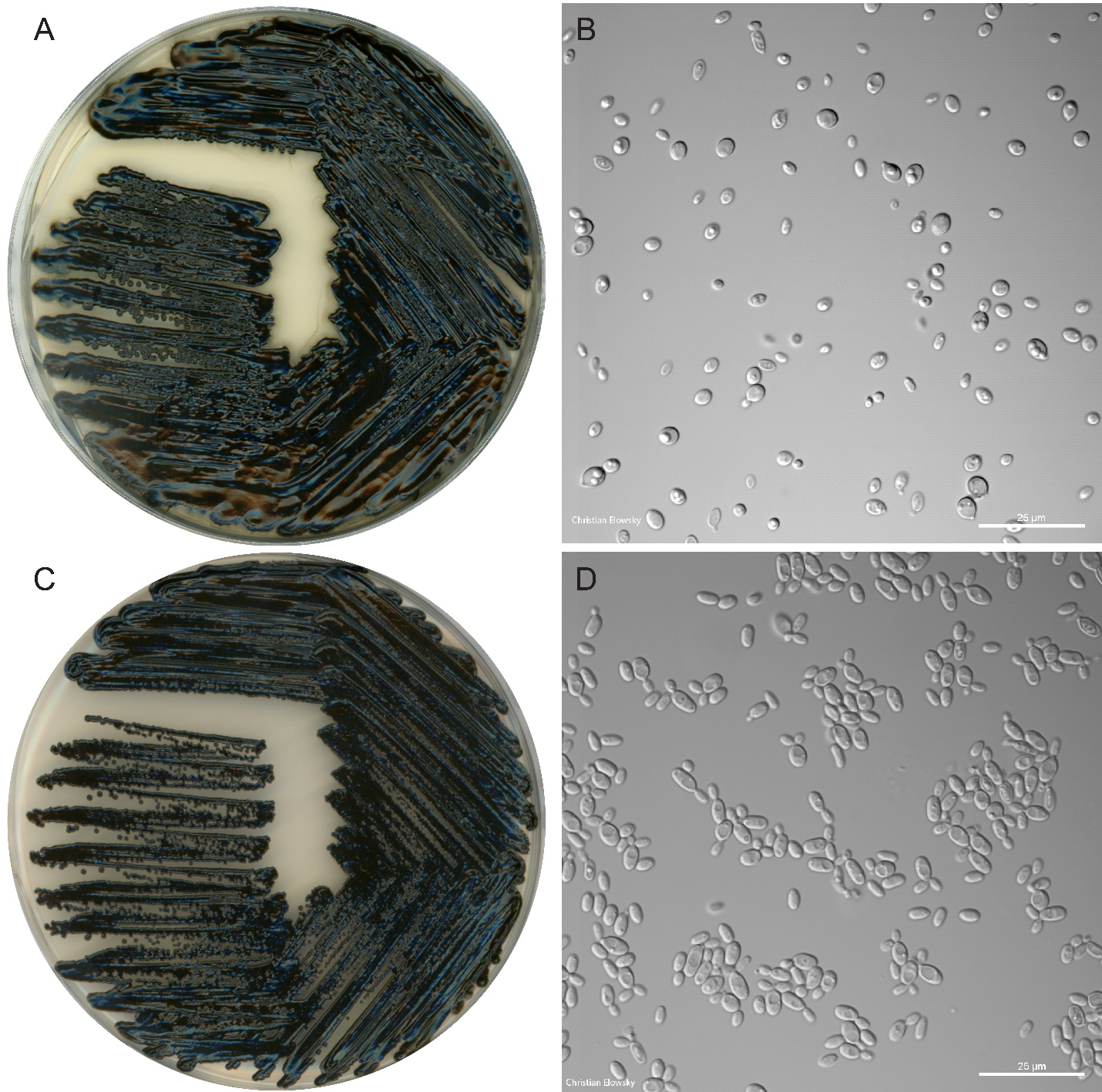

**Supplementary Figure 3:** A) E. viscosa JF 03-3F plate morphology; grown on an MEA plate for 10 days. B) E. viscosa JF 03-3F cell morphology; grown in liquid MEA for 5 days; 60x objective lens. C) E. viscosa JF 03-4F plate morphology; grown on a MEA plate for 10 days. D) E. viscosa JF 03-4F cell morphology; grown in liquid MEA for 5 days; 60x objective lens. (Both plate photos and microscopy photos were taken by Christian Elowsky)

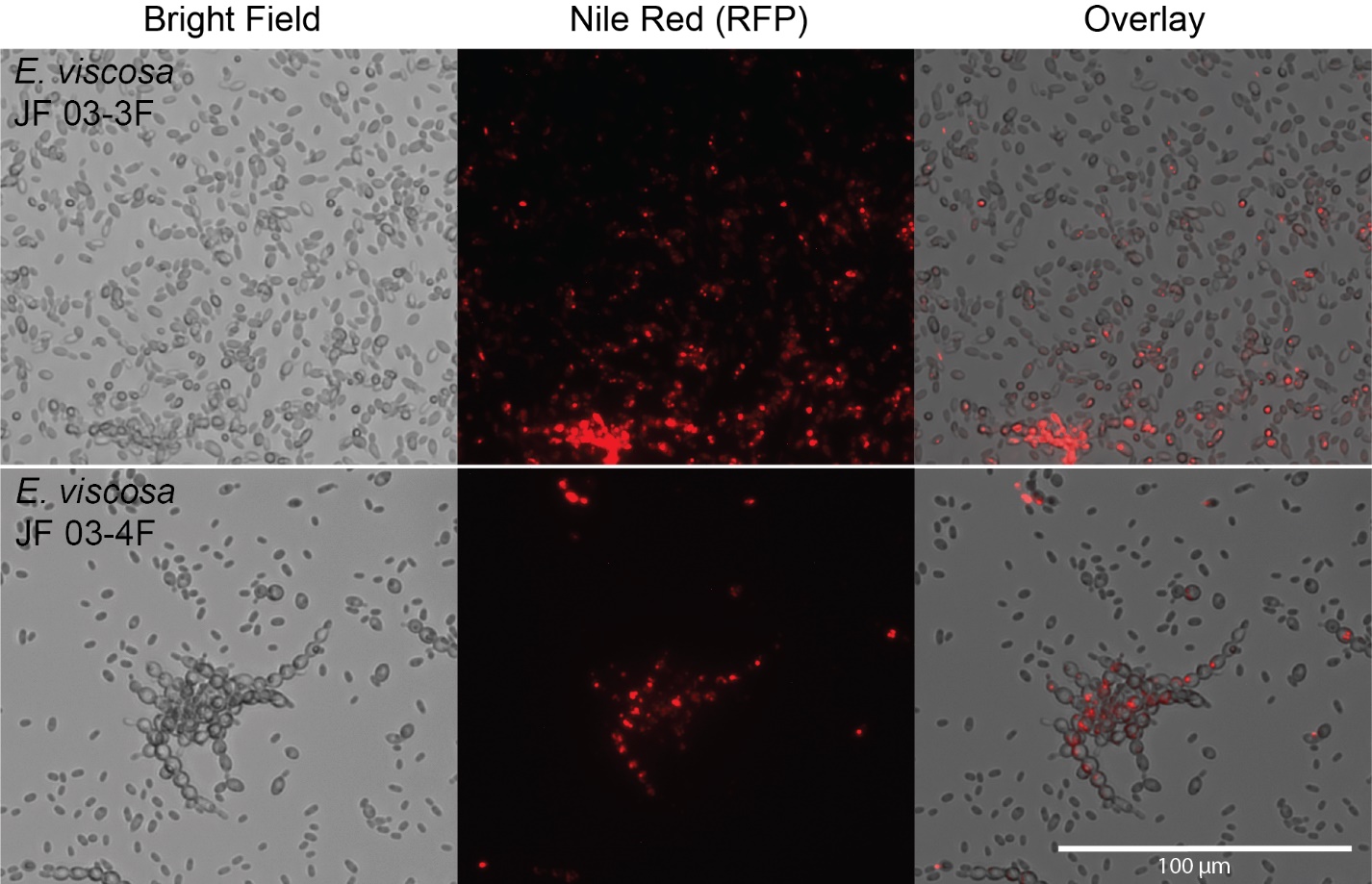

**Supplementary Figure 4:** Nile red micrographs of E. viscosa JF 03-3F and E. viscosa JF 03-4F to display their lipid bodies. Both E. viscosa JF 03-3F and E. viscosa JF 03-4F harbor lipid bodies. Cells were grown in liquid MEA for 3 days; Nile red solution was 50 μg/mL; 1 μL of Nile red was used for every 10 μL of cells. Scale bar represents scale for all images.

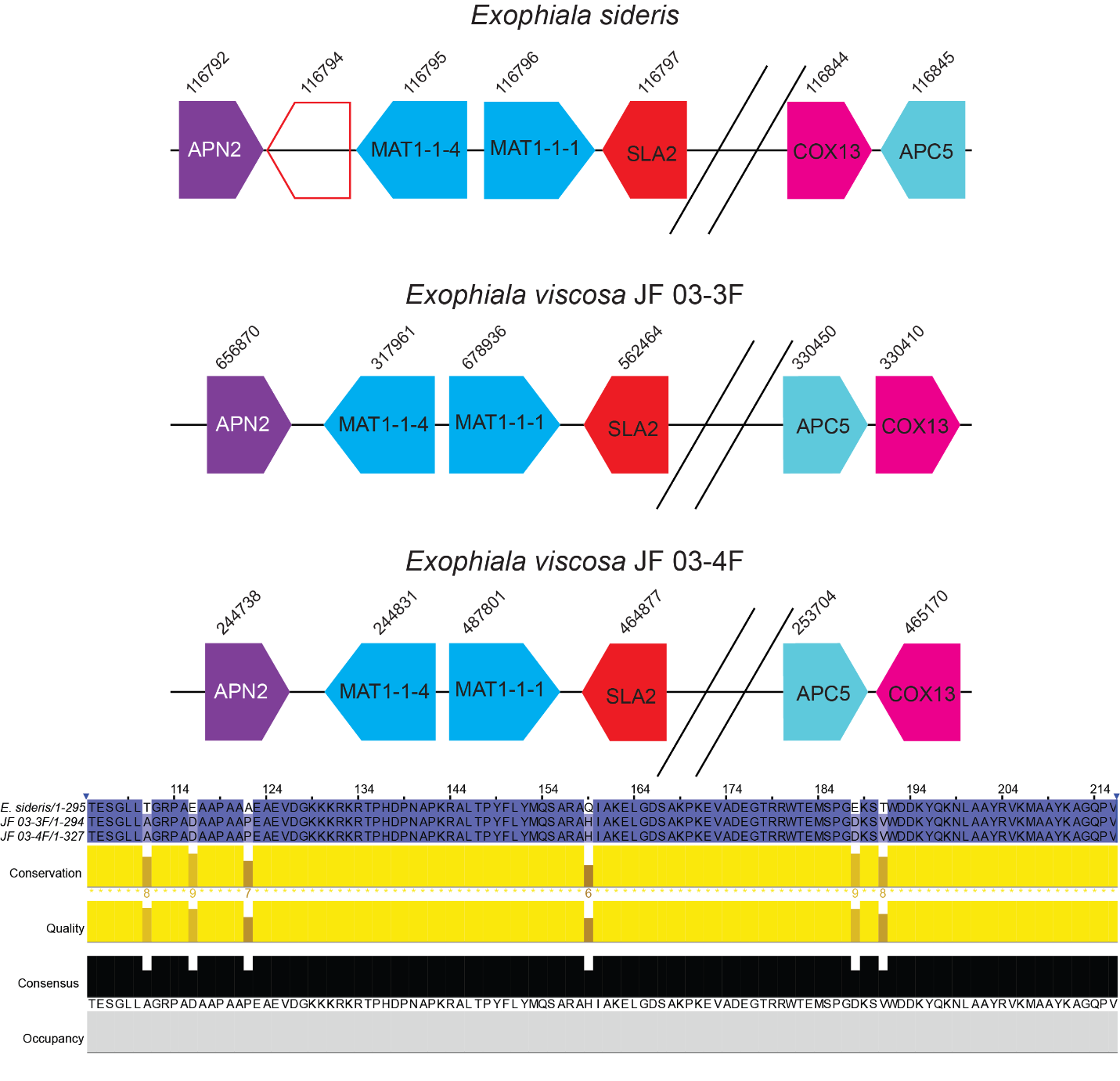

**Supplementary Figure 5:** MAT loci gene order for E. sideris, E. viscosa JF 03-3F, and E. viscosa JF 03-4F. In all three species the same genes are present within the MAT locus. E. sideris is indicated to have a gene for a hypothetical protein between APN2 and MAT1-1-4, whereas E. viscosa JF 03-3F and E. viscosa JF 03-4F were not predicted to have that gene. Additionally, all three species have COX 13 and APC5 downstream of their MAT loci, but the gene order or orientation is different amongst the three species. Numbers above the genes represent their protein ID numbers in the JGI Mycocosm database. Bottom part shows a portion of the protein sequences of the MAT1-1-1 for E. sideris, E. viscosa JF 03-3F, and E. viscosa JF 03-4F. All three species share high homology in this protein, dark blue amino acids mean all three species have the same amino acid in that position.

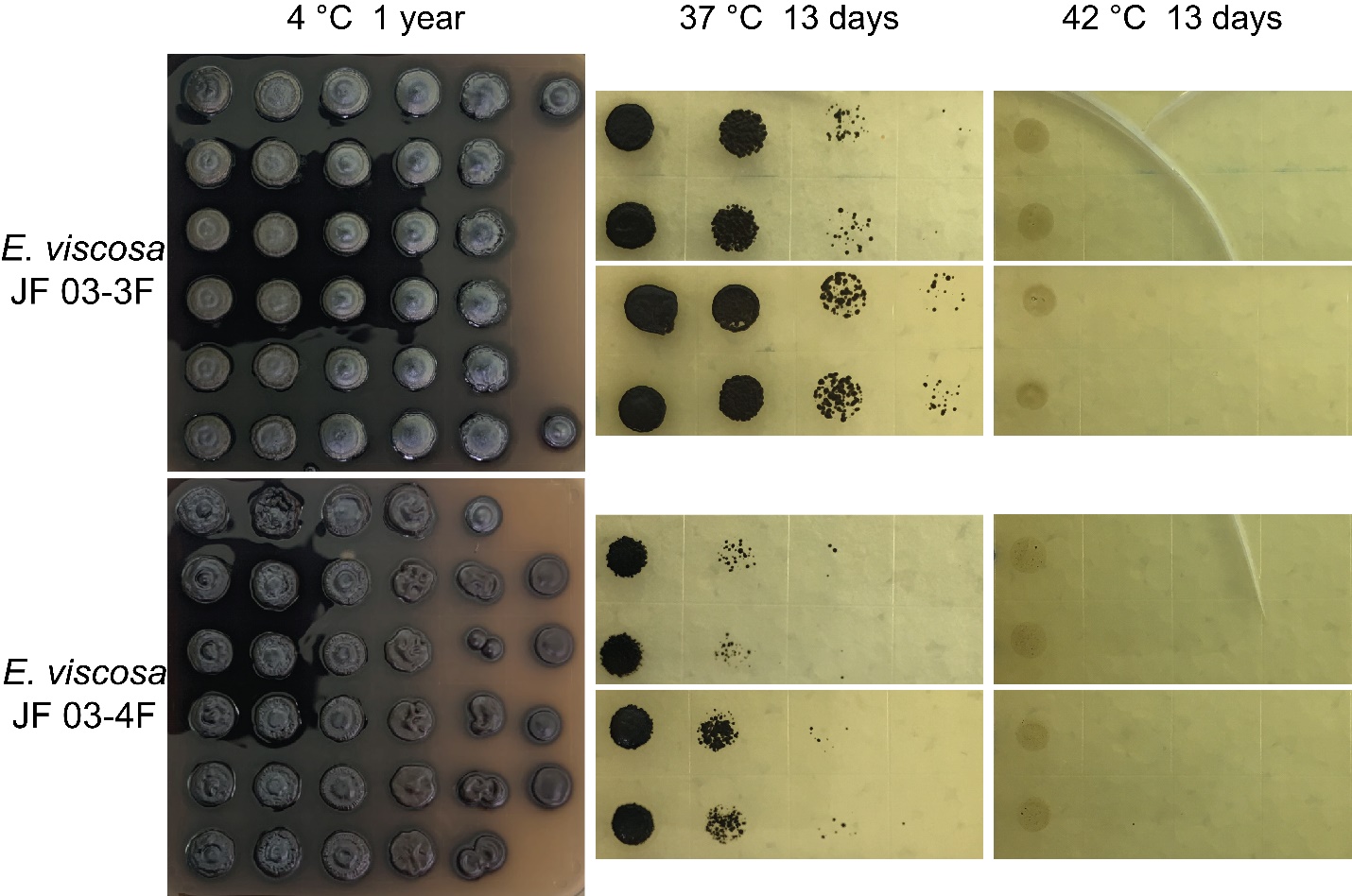

**Supplementary Figure 6:** Growth of E. viscosa JF 03-3F and E. viscosa JF 03-4F at the lowest and highest temperatures tested, after prolonged periods of time. Growth at 4 °C continued for a year in both species, indicating that they can grow at these lower temperatures for extended periods of time. Additionally, we observed that while neither species was capable of active growth at 37 °C, 48 hours was not too long of an exposure time to kill these cells. Whereas at 42 °C neither species was capable of growth, and both were killed after 48 hours of exposure.

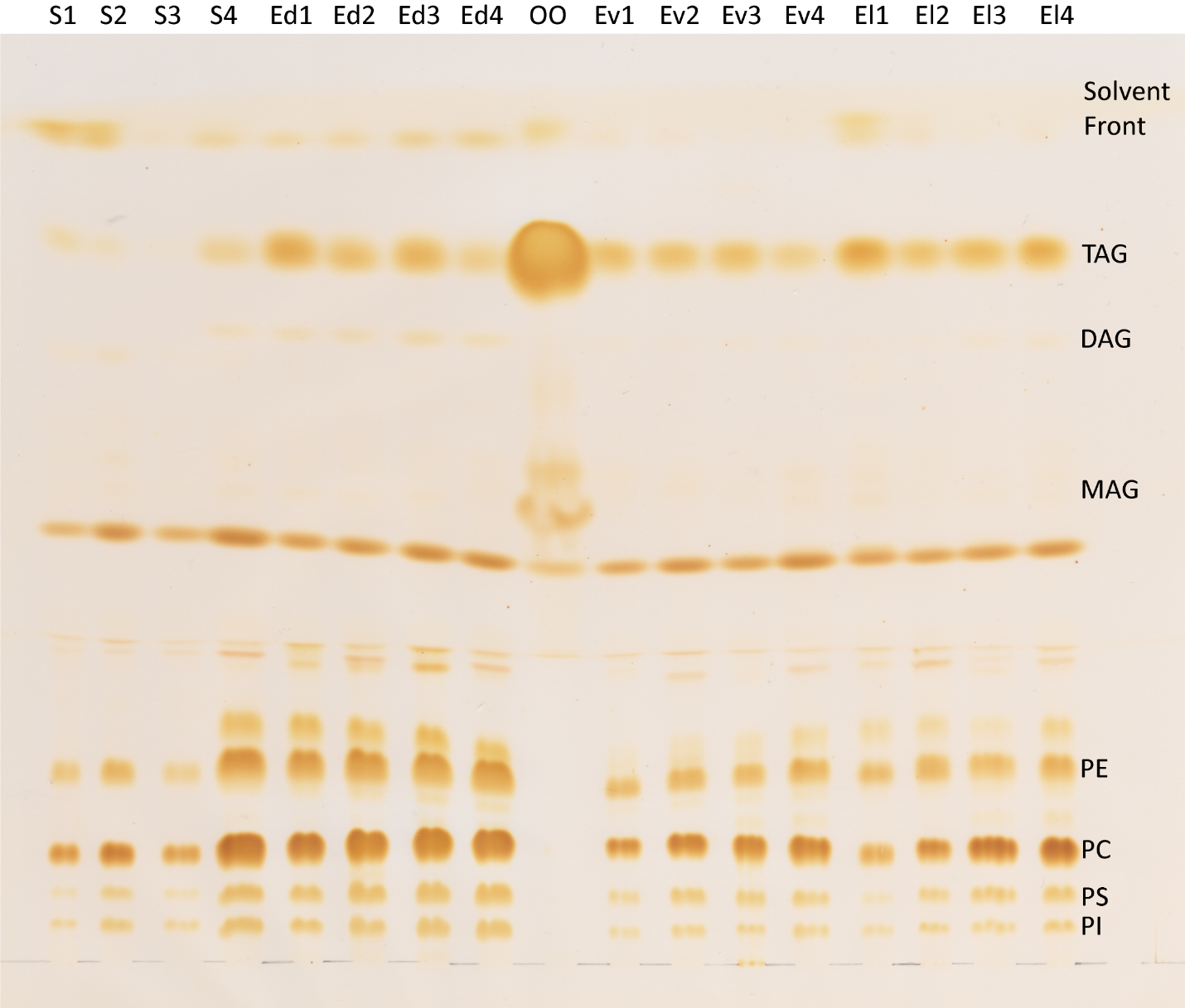

**Supplementary Figure 7:** Lipid profile of S. cerevisiae (Sc), E. dermatitidis (Ed), E. viscosa JF 03-3F JF 03-3F (Ev), and E. viscosa JF 03-4F (El) using four different medias (1: MEA, 2: MEA + 2% peptone, 3: MEA + 2% glycerol, 4: MEA + 2% glycerol + 2% peptone). Differences in fermentable vs. non-fermentable carbon sources and amount of nitrogen source did not alter the amount or types of lipids produced by either E. viscosa JF 03-3F or E. viscosa JF 03-4F. These fungi also showed no unique lipid production or any extreme accumulations of any lipids when compared to other fungi. Lipid codes: PI = Phosphotidylinositol; PS = Phosphotidylserine; PC = Phosphotidylcholine; PE = Phosphotidylethanolamine; MAG = Monoacylglycerol; DAG = Diacylglycerol; TAG = Triacylglycerol.

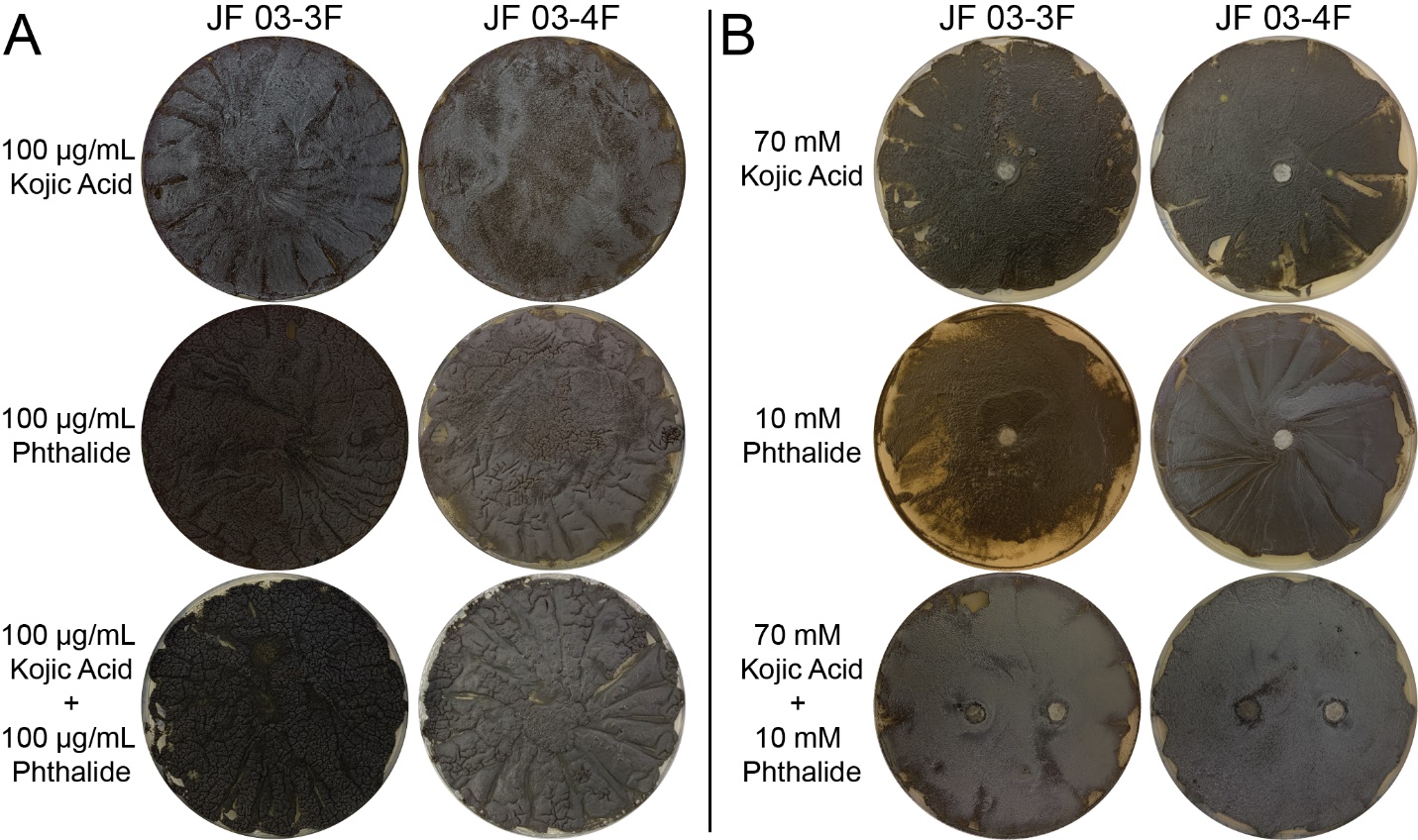

**Supplementary Figure 8:** (A) E. viscosa JF 03-3F and E. viscosa JF 03-4F lawns grown on MEA media contain either 100 μg/mL of kojic acid, 100 μg/mL of phthalide, or both in hopes of blocking melanin production through chemical means. Neither the individual melanin blockers nor their combined efforts were able to block melanin production in either fungus. (B) The same chemicals were used to attempt a different method of melanin blocking but at different concentrations, 10 mg/mL for kojic acid and 10 mM for phthalide. These compounds were added to filter discs and placed on lawns of E. viscosa JF 03-3F and E. viscosa JF 03-4F. Neither the individual compounds nor the combined compounds blocked melanin production.

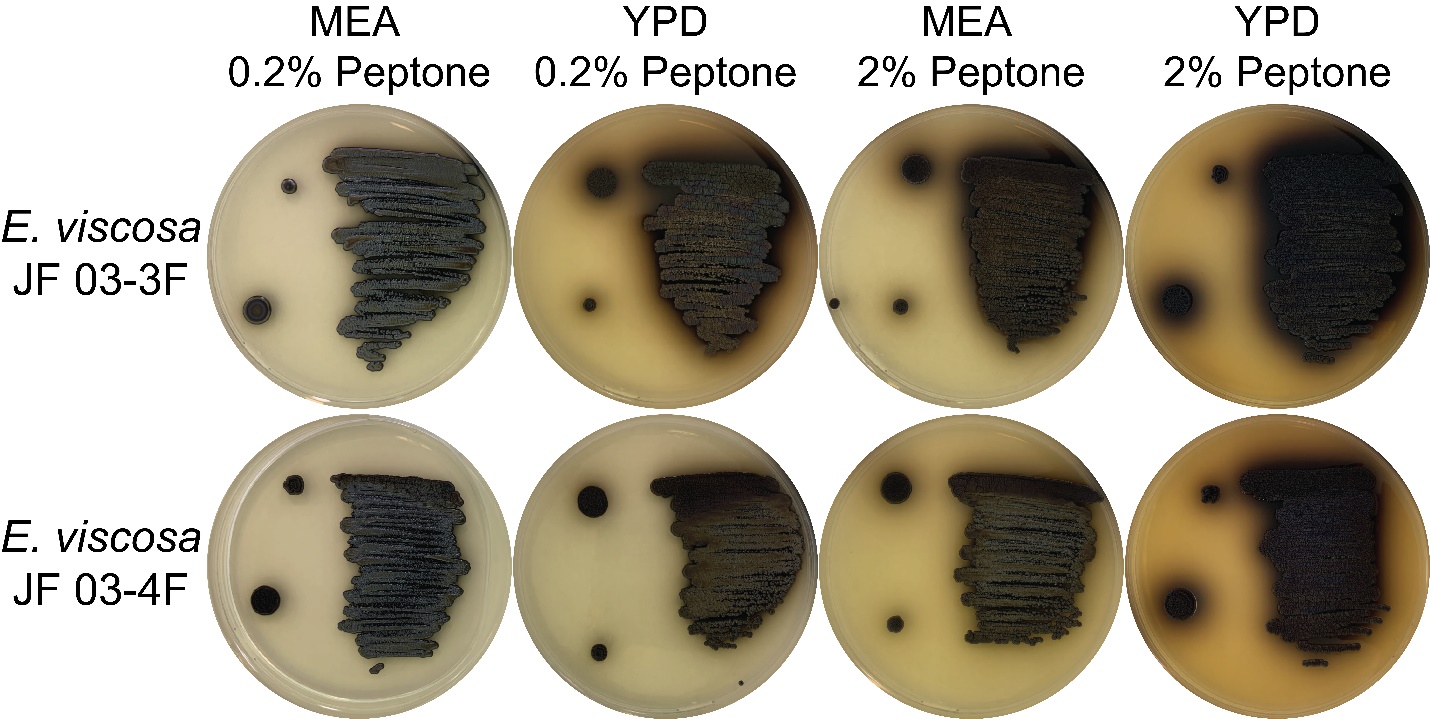

**Supplementary Figure 9:** E. viscosa JF 03-3F and E. viscosa JF 03-4F grown on MEA and YPD with different concentrations of peptone. E. viscosa JF 03-3F is capable of melanin excretion on MEA with 2% peptone, which is the same amount of peptone in regular YPD. E. viscosa JF 03-4F was not as capable of secreting melanin in the MEA + 2% peptone, but there is a slight amount of excreted melanin. E. viscosa JF 03-3F was also capable of secreting melanin on YPD with 0.2% peptone, indicating that yeast extract might have more available nitrogen than malt extract.

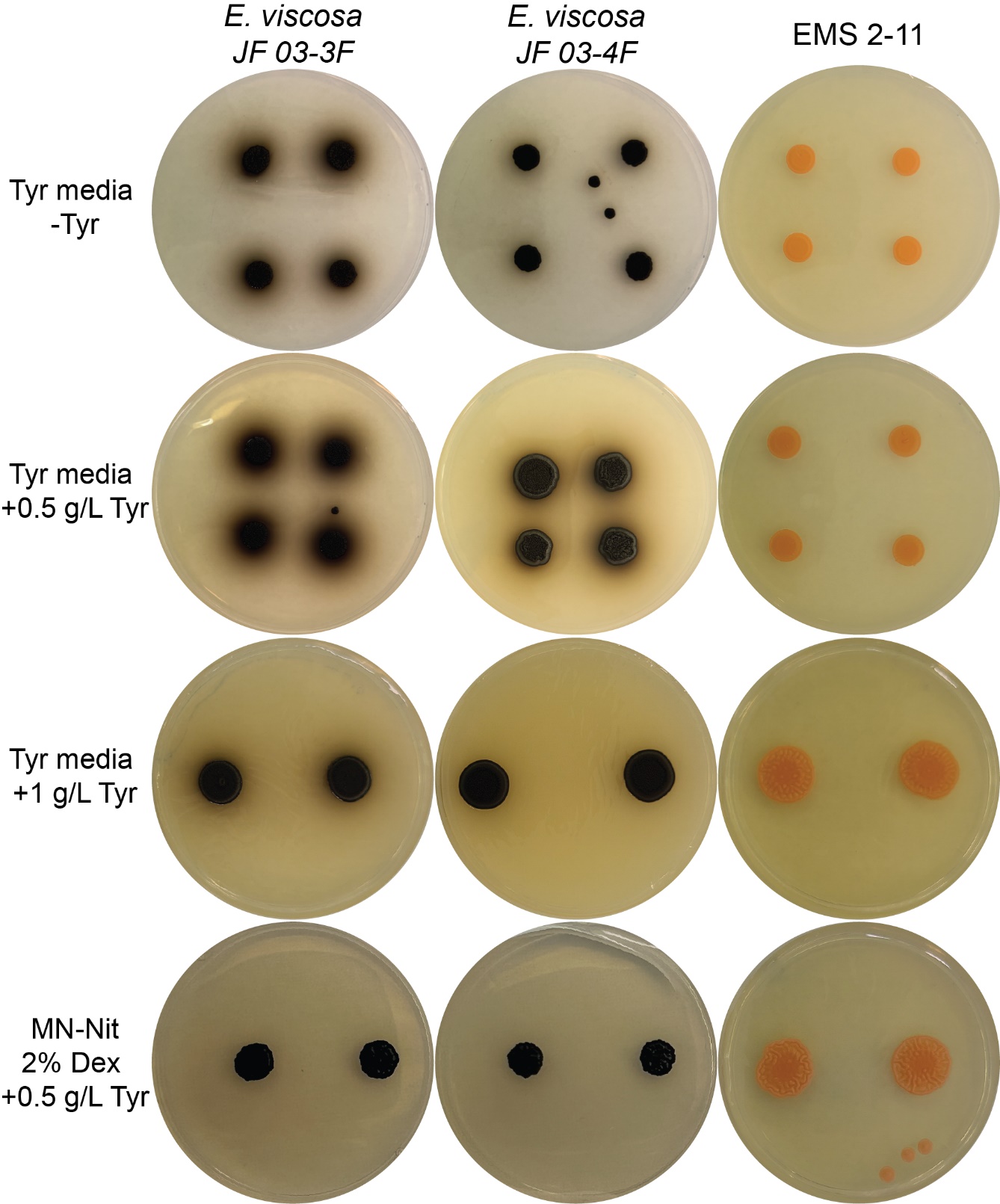

**Supplementary Figure 10:** E. viscosa JF 03-3F, E. viscosa JF 03-4F, and the albino mutant of E. viscosa JF 03-4F EMS 2-11 grown on Tyrosine media (Tyr media: 2% dextrose; 1% peptone; 0.1% yeast extract (Jalmi et al., 2012)) with or without tyrosine added at 0.5 g/L and 1 g/L, and on MN-Nitrate 2% dextrose +0.5 g/L tyrosine; day 19 of growth. This was done to determine if the tyrosine in peptone could be causing the melanin excretion, or if tyrosine alone can induce melanin excretion. If tyrosine alone was causing the excretion via the tyrosine-dependent biosynthetic pathways, then EMS 2-11 would also be excreting melanin since it is only mutated in pks1 which affects allomelanin production and does not require tyrosine as a precursor. While adding 0.5 g/L of tyrosine does seem to induce melanin excretion, adding 1 g/L seems to halt melanin excretion. Growth on tyrosine as a sole nitrogen source also does not seem to induce melanin excretion in either the wild type strains or the albino mutant. The peptone vs. tyrosine melanin excretion phenomenon, and the melanin excretion phenomenon overall is still being studied.

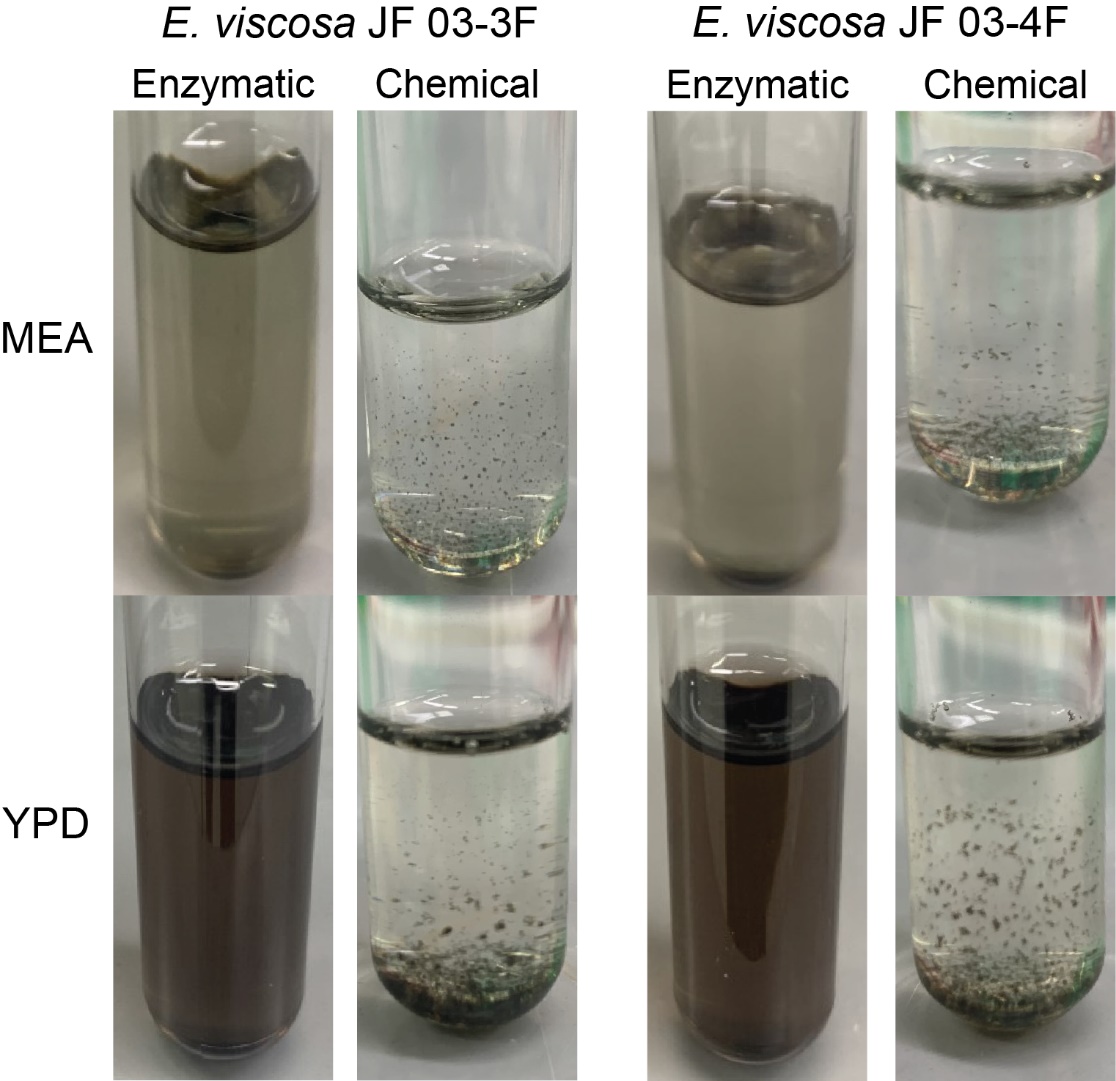

**Supplementary Figure 11:** Extraction of melanin from supernatants of E. viscosa JF 03-3F and E. viscosa JF 03-4F using both enzymatic and chemical methods described in (Pralea et al., 2019). Enzymatic extraction methods were incapable of extracting all the melanin, leaving behind a dark supernatant in the last step. However, melanin extracted by chemical extraction methods had complete extraction of the secreted melanin.

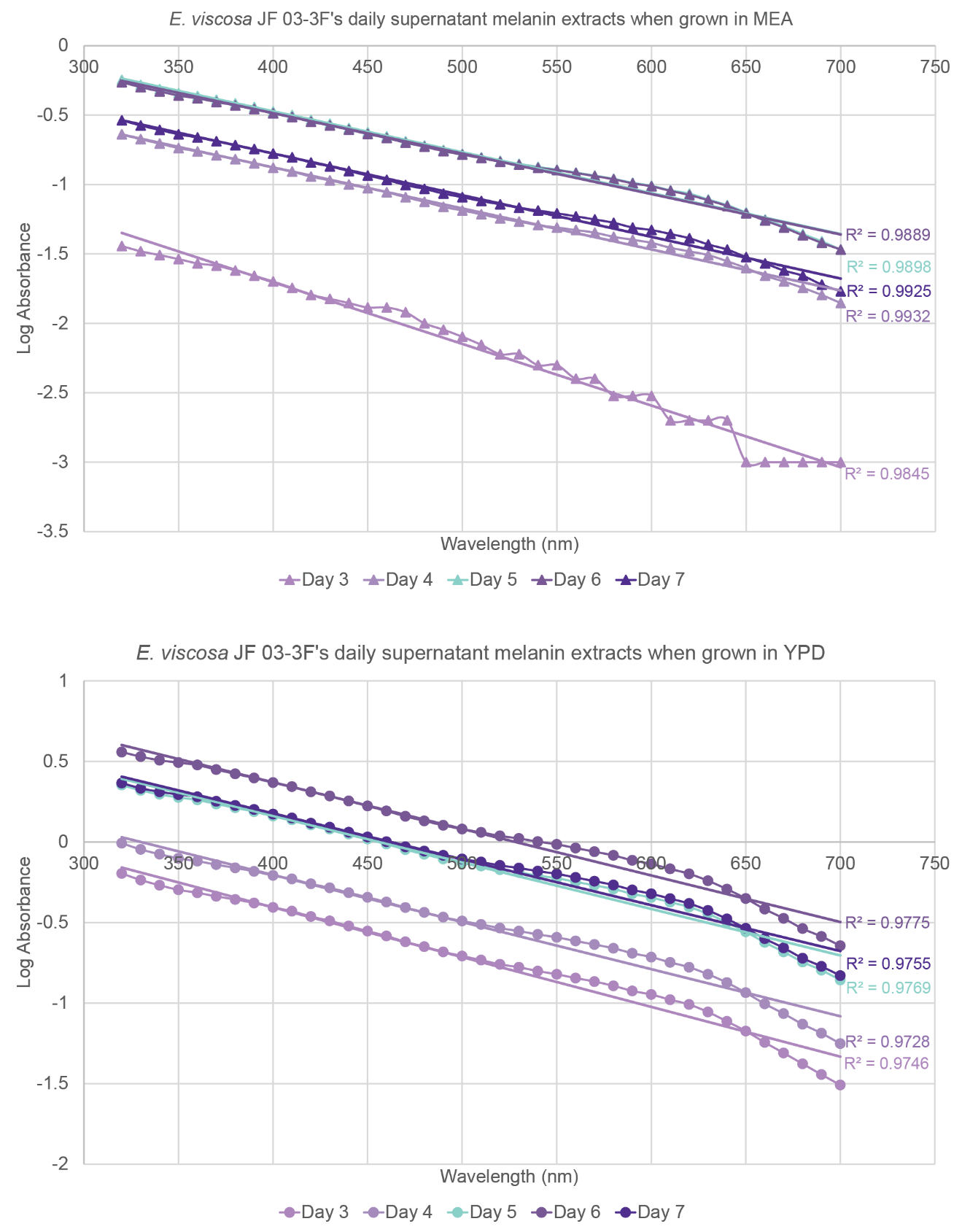

**Supplementary Figure 12:** Log absorbance values of the extracted melanin from the supernatant of E. viscosa JF 03-3F. All R^2^ values of both MEA and YPD from every day are R^2^ ≥ 0.97, meaning that the compound that was extracted was melanin for all samples.

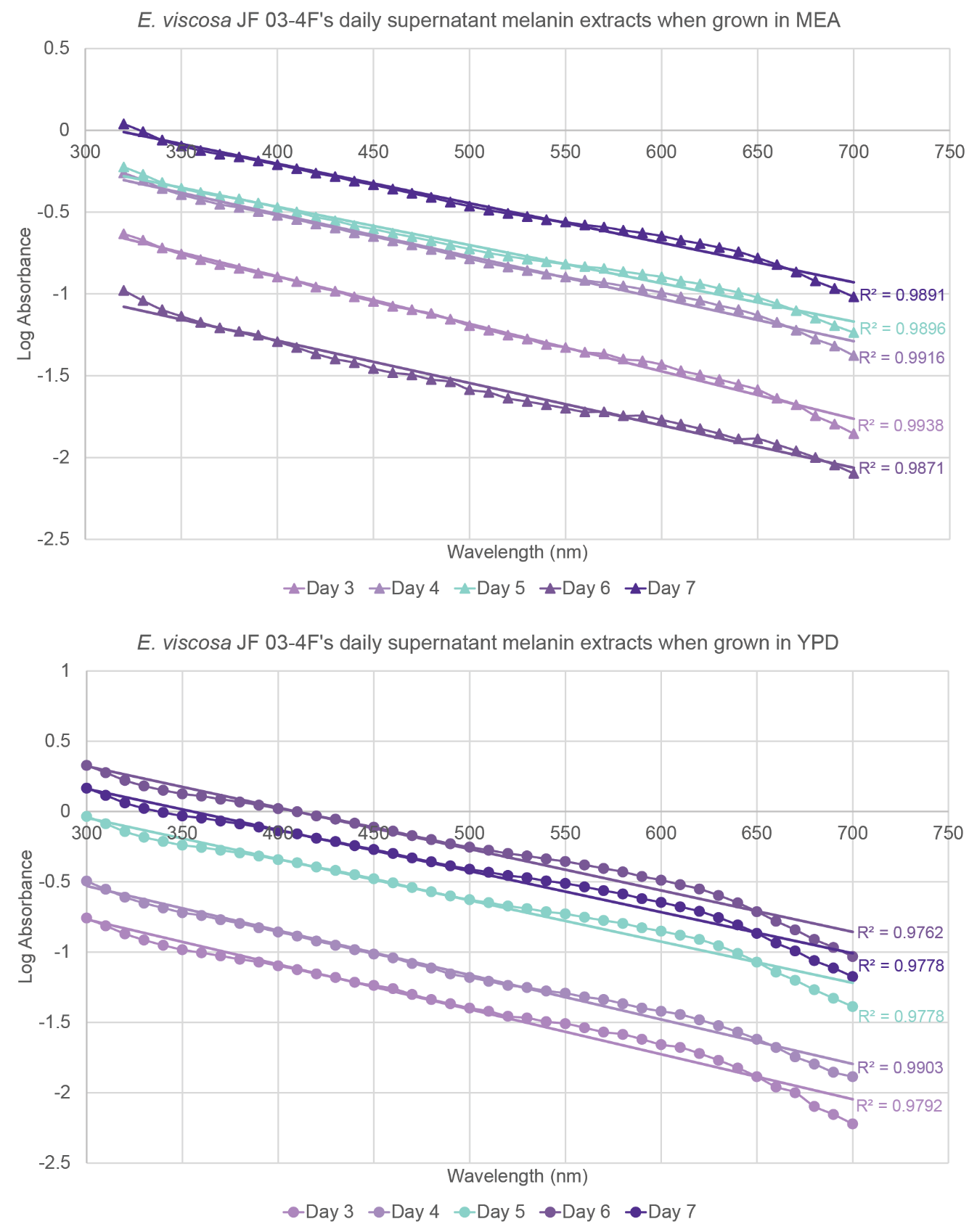

**Supplementary Figure 13**: Log absorbance values of the extracted melanin from the supernatant of E. viscosa JF 03-4F. All R^2^ values of both MEA and YPD from every day are R^2^ ≥ 0.97, meaning that the compound that was extracted was melanin for all samples.

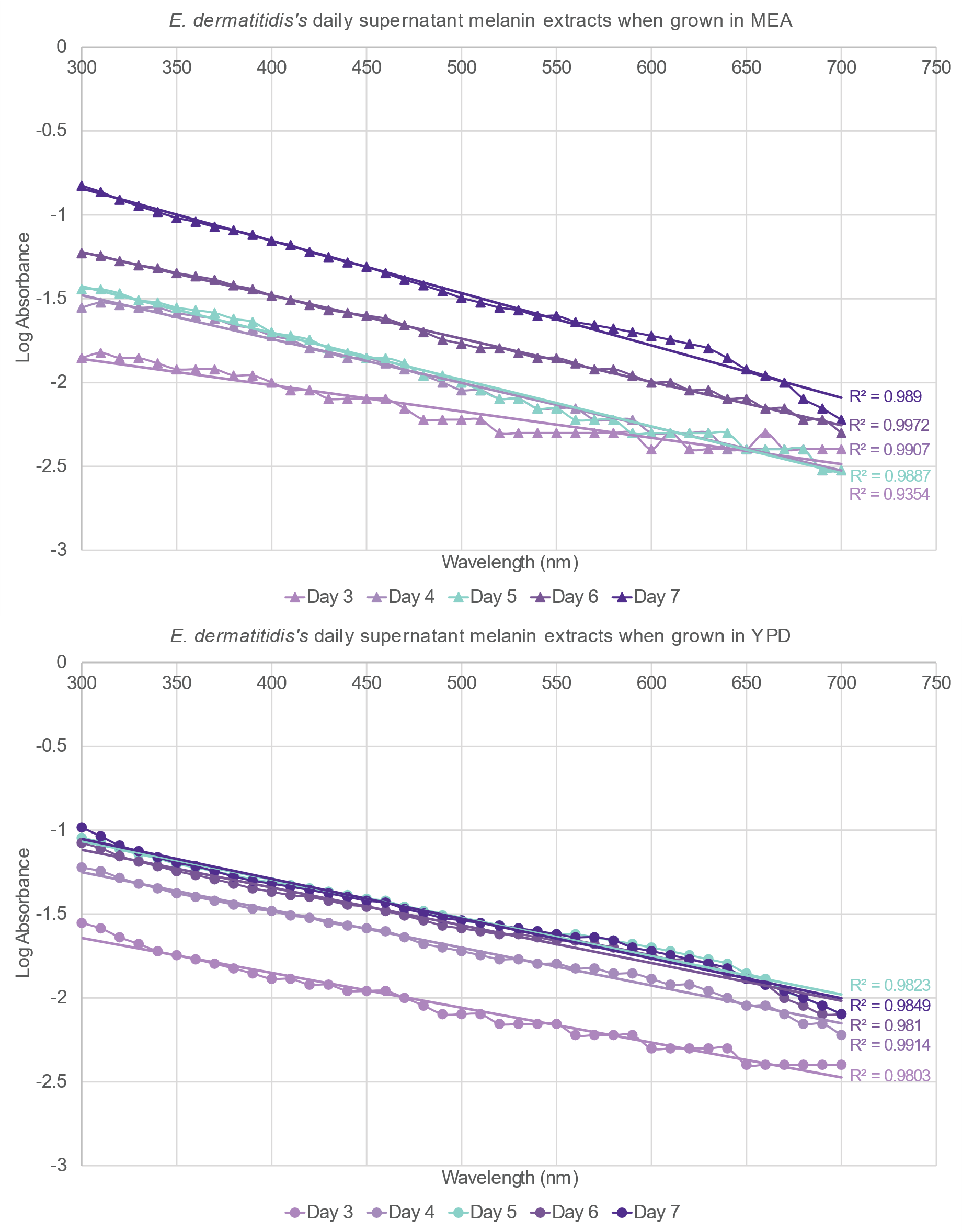

**Supplementary Figure 14**: Log absorbance values of the extracted melanin from the supernatant of E. dermatitidis. All R^2^ values of both MEA and YPD from every day are R^2^ ≥ 0.97, meaning that the compound that was extracted was melanin for all samples.

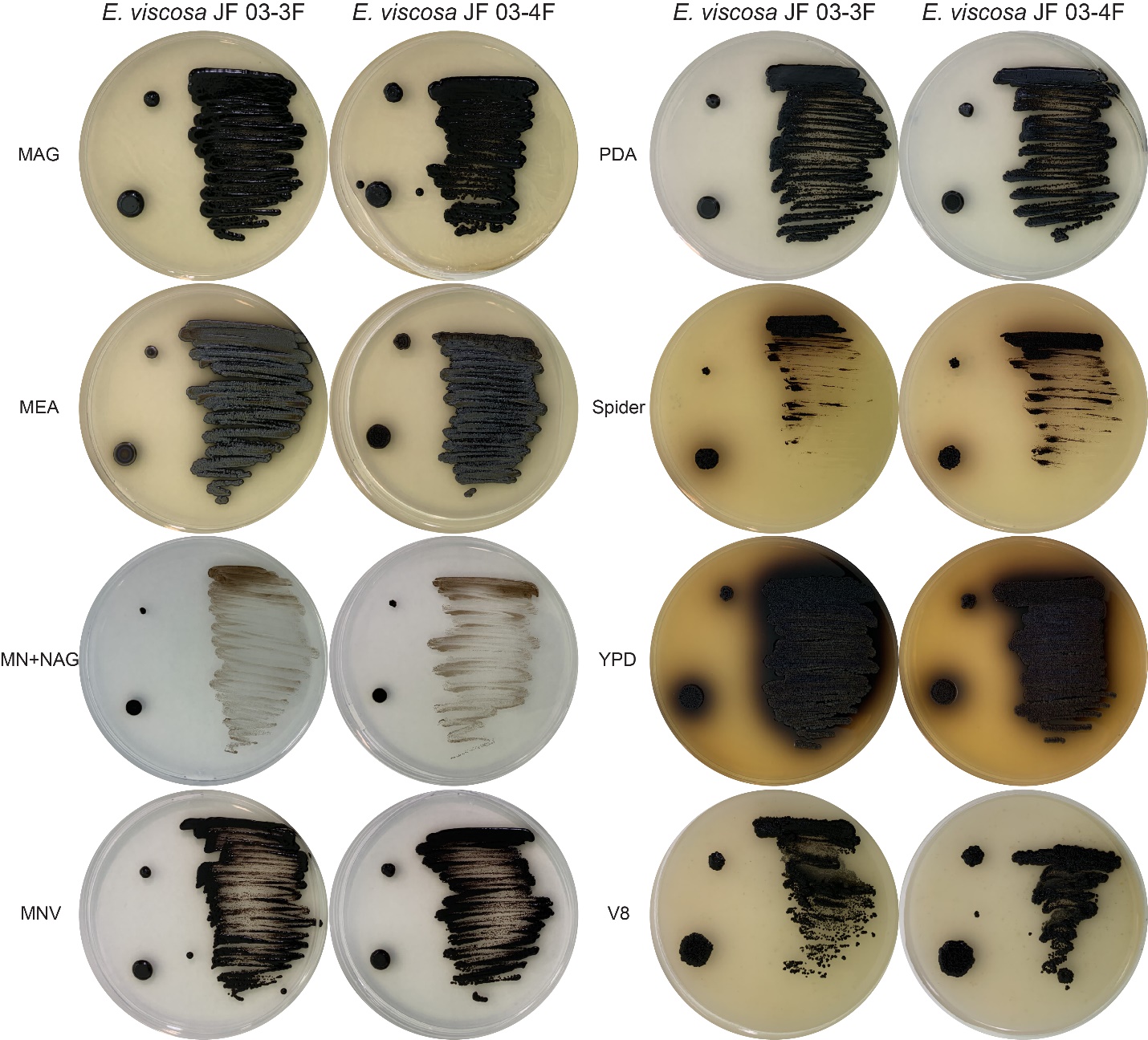

**Supplementary Figure 15:** Original photos of E. viscosa JF 03-3F and E. viscosa JF 03-4F on different media types.

**Supplemental Table 1:** R^2^ values for serially-diluted cell’s OD vs. labeled columns to determine best wavelength for OD measurements

| **R^2^ Values** | **E. viscosa** JF 03-3F | | | | | **E. viscosa** JF 03-4F | | | | |
| --- | --- | --- | --- | --- | --- | --- | --- | --- | --- | --- |
| **λ** | **25% > % Diff** | **50% > % Diff** | **2.19x10^6 > cells** | **4.38x10^6 > cells** | **Actual % Diff 12.5% > consistency** | **25% > % Diff** | **50% > % Diff** | **2.29x10^6 > Cells** | **4.58x10^6 > cells** | **Actual % Diff 12.5% >**  **consistency** |
| 300 | N/A | N/A | 0.99846 | N/A | 0.00458 | N/A | N/A | N/A | N/A | 0.01026 |
| 310 | N/A | N/A | 0.99842 | N/A | 0.00640 | N/A | N/A | N/A | N/A | 0.01743 |
| 320 | N/A | N/A | 0.99811 | N/A | 0.01865 | N/A | N/A | N/A | N/A | 0.05374 |
| 330 | N/A | N/A | 0.99784 | N/A | 0.02876 | 0.98023 | 0.83057 | 0.98023 | 0.83057 | 0.01681 |
| 340 | N/A | N/A | 0.99762 | N/A | 0.01722 | 0.97850 | 0.84832 | 0.97850 | 0.84832 | 0.00108 |
| 350 | N/A | N/A | 0.99731 | N/A | 0.03011 | 0.97763 | 0.86215 | 0.97763 | 0.86215 | 0.02053 |
| 360 | 0.99072 | 0.82656 | 0.99702 | 0.99072 | 0.02590 | 0.97810 | 0.86867 | 0.97810 | 0.86867 | 0.02595 |
| 370 | 0.98952 | 0.84428 | 0.99661 | 0.98952 | 0.00347 | 0.97727 | 0.87773 | 0.97727 | 0.87773 | 0.06769 |
| 380 | 0.98851 | 0.86061 | 0.99617 | 0.98851 | 0.00016 | 0.97703 | 0.88556 | 0.97703 | 0.88556 | 0.08173 |
| 390 | 0.98835 | 0.87409 | 0.99571 | 0.98835 | 0.00354 | 0.97637 | 0.89575 | 0.97637 | 0.89575 | 0.10751 |
| 400 | 0.98662 | 0.89107 | 0.99514 | 0.98662 | 0.01706 | 0.97418 | 0.91200 | 0.97418 | 0.91200 | 0.14472 |
| 410 | 0.98525 | 0.90268 | 0.99469 | 0.98525 | 0.04220 | 0.97217 | 0.92601 | 0.97217 | 0.92601 | 0.16328 |
| 420 | 0.98445 | 0.91489 | 0.99407 | 0.98445 | 0.04658 | 0.96926 | 0.94179 | 0.96926 | 0.94179 | 0.18480 |
| 430 | 0.98281 | 0.92644 | 0.99359 | 0.98281 | 0.06500 | 0.96728 | 0.95122 | 0.96728 | 0.95122 | 0.21924 |
| 440 | 0.98036 | 0.94096 | 0.99269 | 0.98036 | 0.10829 | 0.96531 | 0.95960 | 0.96531 | 0.95960 | 0.22432 |
| 450 | 0.97762 | 0.95450 | 0.99165 | 0.97762 | 0.13724 | 0.96379 | 0.95549 | 0.96379 | 0.95549 | 0.21414 |
| 460 | 0.97466 | 0.96527 | 0.99057 | 0.97466 | 0.17287 | 0.96231 | 0.95349 | 0.96231 | 0.95349 | 0.22154 |
| 470 | 0.97182 | 0.96242 | 0.98980 | 0.97182 | 0.19849 | 0.96148 | 0.95299 | 0.96148 | 0.95299 | 0.21765 |
| 480 | 0.96979 | 0.95724 | 0.98916 | 0.96979 | 0.21771 | 0.96066 | 0.95026 | 0.96066 | 0.95026 | 0.23401 |
| 490 | 0.96828 | 0.95392 | 0.98879 | 0.96828 | 0.25068 | 0.96011 | 0.94857 | 0.96011 | 0.94857 | 0.23096 |
| 500 | 0.96728 | 0.95162 | 0.98841 | 0.96728 | 0.28489 | 0.96003 | 0.94743 | 0.96003 | 0.94743 | 0.21163 |
| 510 | 0.96655 | 0.94995 | 0.98815 | 0.96655 | 0.31610 | 0.95968 | 0.94661 | 0.95968 | 0.94661 | 0.22175 |
| 520 | 0.96615 | 0.94870 | 0.98814 | 0.96615 | 0.28625 | 0.95934 | 0.94493 | 0.95934 | 0.94493 | 0.20395 |
| 530 | 0.96585 | 0.94793 | 0.98818 | 0.96585 | 0.28547 | 0.95938 | 0.94451 | 0.95938 | 0.94451 | 0.20779 |
| 540 | 0.96566 | 0.94702 | 0.98828 | 0.96566 | 0.25322 | 0.95976 | 0.94433 | 0.95976 | 0.94433 | 0.19645 |
| 550 | 0.96565 | 0.94668 | 0.98825 | 0.96565 | 0.29927 | 0.95963 | 0.94374 | 0.95963 | 0.94374 | 0.17600 |
| 560 | 0.96615 | 0.94728 | 0.98847 | 0.96615 | 0.26053 | 0.95968 | 0.94292 | 0.95968 | 0.94292 | 0.19001 |
| 570 | 0.96665 | 0.94763 | 0.98864 | 0.96665 | 0.25148 | 0.95996 | 0.94301 | 0.95996 | 0.94301 | 0.17008 |
| 580 | 0.96699 | 0.94835 | 0.98884 | 0.96699 | 0.22055 | 0.96034 | 0.94279 | 0.96034 | 0.94279 | 0.15510 |
| 590 | 0.96755 | 0.94880 | 0.98911 | 0.96755 | 0.20711 | 0.96057 | 0.94254 | 0.96057 | 0.94254 | 0.16411 |
| 600 | 0.96802 | 0.94920 | 0.98934 | 0.96802 | 0.19794 | 0.96077 | 0.94249 | 0.96077 | 0.94249 | 0.14509 |
| 610 | 0.96842 | 0.94989 | 0.98967 | 0.96842 | 0.21037 | 0.96113 | 0.94235 | 0.96113 | 0.94235 | 0.13176 |
| 620 | 0.96888 | 0.95030 | 0.98973 | 0.96888 | 0.20081 | 0.96125 | 0.94203 | 0.96125 | 0.94203 | 0.14060 |
| 630 | 0.96918 | 0.95013 | 0.98998 | 0.96918 | 0.21601 | 0.96139 | 0.94144 | 0.96139 | 0.94144 | 0.12192 |
| 640 | 0.96949 | 0.95067 | 0.99005 | 0.96949 | 0.18258 | 0.96167 | 0.94142 | 0.96167 | 0.94142 | 0.13532 |
| 650 | 0.97059 | 0.94987 | 0.99018 | 0.97059 | 0.20795 | 0.96193 | 0.94087 | 0.96193 | 0.94087 | 0.11326 |
| 660 | 0.97084 | 0.95000 | 0.99040 | 0.97084 | 0.17514 | 0.96205 | 0.94076 | 0.96205 | 0.94076 | 0.12267 |
| 670 | 0.97096 | 0.94988 | 0.99050 | 0.97096 | 0.17077 | 0.96210 | 0.94031 | 0.96210 | 0.94031 | 0.11035 |
| 680 | 0.97094 | 0.95005 | 0.99061 | 0.97094 | 0.19038 | 0.96245 | 0.93989 | 0.96245 | 0.93989 | 0.10061 |
| 690 | 0.97090 | 0.94991 | 0.99072 | 0.97090 | 0.18331 | 0.96260 | 0.93971 | 0.96260 | 0.93971 | 0.10992 |
| 700 | 0.97072 | 0.94937 | 0.99071 | 0.97072 | 0.15878 | 0.96272 | 0.93939 | 0.96272 | 0.93939 | 0.10001 |

**Supplemental Table 2:** R^2^ values and slopes after plotting the log absorbance of the supernatant samples against wavelength

|  | *E. viscosa* JF 03-3F | | *E. viscosa* JF 03-4F | | *E. dermatitidis* | |
| --- | --- | --- | --- | --- | --- | --- |
|  | R^2^ | Slope | R^2^ | Slope | R^2^ | Slope |
| **YPD** | | | | | | |
| Day 3 | 0.9746 | -0.0031 | 0.9792 | -0.0032 | 0.9803 | -0.0021 |
| Day 4 | 0.9727 | -0.0029 | 0.9903 | -0.0031 | 0.9914 | -0.0023 |
| Day 5 | 0.9768 | -0.0029 | 0.9778 | -0.0029 | 0.9823 | -0.0023 |
| Day 6 | 0.9774 | -0.0029 | 0.9762 | -0.0029 | 0.9810 | -0.0023 |
| Day 7 | 0.9755 | -0.0028 | 0.9778 | -0.0029 | 0.9849 | -0.0024 |
| **MEA** | | | | | | |
| Day 3 | 0.9845 | -0.0044 | 0.9938 | -0.0029 | 0.9354 | -0.0016 |
| Day 4 | 0.9932 | -0.0029 | 0.9916 | -0.0026 | 0.9907 | -0.0026 |
| Day 5 | 0.9898 | -0.0029 | 0.9896 | -0.0023 | 0.9887 | -0.0028 |
| Day 6 | 0.9889 | -0.0029 | 0.9871 | -0.0026 | 0.9972 | -0.0026 |
| Day 7 | 0.9925 | -0.003 | 0.9891 | -0.0024 | 0.9890 | -0.0031 |

Jalmi, P., Bodke, P., Wahidullah, S., & Raghukumar, S. (2012). The fungus Gliocephalotrichum simplex as a source of abundant, extracellular melanin for biotechnological applications. *World Journal of Microbiology and Biotechnology*, *28*(2), 505-512. <https://doi.org/10.1007/s11274-011-0841-0>

**Supplemental Table 3:** Media variations tested on E. viscosa strains for inducing melanin excretion and their outcomes.

| Base media | Component Removed | Added nitrogen | Added carbon | Added other | Days of growth | Melanin excreted | |
| --- | --- | --- | --- | --- | --- | --- | --- |
|  |  |  |  |  |  | JF 03-3F | JF 03-4F |
| MN | N/A | 2% peptone | 2% dextrose | N/A | 16 | No | No |
| MN | NaNO_3_ | 2% peptone | 2% dextrose | N/A | 16 | No | No |
| MN | NaNO_3_ | 0.5 g/L Tyr | 2% dextrose | N/A | 16 | No | No |
| MN | Trace elements | 2% peptone | N/A | N/A | 45 | No | No |
| MN | Trace elements | 0.5 g/L Tyr | N/A | N/A | 45 | No | No |
| MEA | Peptone | 2% w/v NH_4_SO_4_ | N/A | N/A | 19 | No | No |
| MEA | Peptone | 10x AA stock | N/A | N/A | 10 | No | No |
| MEA | Peptone | 0.5 g/L Tyr | N/A | N/A | 26 | Yes | Yes |
| MEA | Peptone | 1 g/L Tyr | N/A | N/A | 26 | Yes | No |
| YPD | Peptone | 20x AA stock | N/A | N/A | 32 | Yes | Yes |
| YPD | Peptone | 2% w/v NH_4_SO_4_ | N/A | N/A | 32 | No | No |
| YPD | N/A | N/A | N/A | Trace Elements | 17 | No | No |
| BBM | N/A | 0.5 g/L Tyr | 2% dextrose | N/A | 26 | No | No |
| BBM | N/A | 20x AA | 2% dextrose | N/A | 26 | No | Yes |
| BBM | N/A | 20x AA;  0.5 g/L Tyr | 2% dextrose | N/A | 26 | No | Yes |
